# Patterned embryonic invagination evolved in response to mechanical instability

**DOI:** 10.1101/2023.03.30.534554

**Authors:** Bruno C. Vellutini, Marina B. Cuenca, Abhijeet Krishna, Alicja Szałapak, Carl D. Modes, Pavel Tomancak

## Abstract

Mechanical forces are crucial for driving and shaping tissue morphogenesis during embryonic development, but their relevance for the evolution of development remains poorly understood. Here, we show that a morphogenetic innovation present in fly embryos—a deep epithelial fold known as the cephalic furrow—plays a mechanical role during *Drosophila* gastrulation. By integrating *in vivo* experiments and *in silico* simulations, we find that the formation of the cephalic furrow prevents mechanical instabilities at the head–trunk epithelium by absorbing the compressive stresses generated by concurrent morphogenetic movements of gastrulation, the expansion of mitotic domains and the germ band extension. Furthermore, by comparing the expression of known and novel genes involved in cephalic furrow formation between fly species, we find that the presence of the cephalic furrow is linked to changes in the expression of *buttonhead* transcription factor at the head–trunk boundary. These data suggest that the genetic control of cephalic furrow formation was established through the integration of a new player into the ancestral head–trunk patterning system, and that mechanical instability may have been the selective pressure associated with the evolution of the cephalic furrow. Our findings uncover empirical evidence for how mechanical forces can influence the evolution of morphogenetic innovations in early development.

## Main

Morphogenesis is a physical process that shapes embryonic tissues through cell-generated mechanical forces.^1,2^ These forces drive tissues to bend, extend, and invaginate—movements that also push and pull on their neighboring regions. Such physical interactions provide essential information to embryonic cells throughout development and ultimately shape the final morphology of tissues and organs.^3^ However, how mechanical forces influence the evolution of morphogenesis in early embryonic development remains unclear. To investigate the interplay between genetics and mechanics during the evolution of morphogenesis, we studied a prominent but enigmatic epithelial fold that forms at the head–trunk boundary of flies during gastrulation—the cephalic furrow.^4,5^

Cephalic furrow formation is under strict genetic control. In *Drosophila*, it is one of the earliest morphogenetic events of gastrulation, beginning as paired lateral indentations which quickly invaginate to form a deep epithelial fold at the boundary between the head (procephalon) and the trunk (germ band).^4–6^ The position of the cephalic furrow along the anteroposterior axis is determined with remarkable accuracy^7^ by the zygotic expression of two transcription factors, *buttonhead* (*btd*) and *even skipped* (*eve*), whose domains overlap at the head–trunk boundary by a narrow row of blastoderm cells.^8^ These so-called initiator cells shorten along the apical–basal axis through lateral myosin contractility.^9^ The shortening of initiators drives the initial infolding of the tissue while the mechanical coupling between cells ensures the propagation of a precise, bidirectional morphogenetic wave of tissue folding.^9,10^ The resulting fold spans the entire lateral surface, from dorsal to ventral, making the cephalic furrow a landmark of *Drosophila* gastrulation.^4,6^

Despite its prominence and underlying genetic control, the cephalic furrow has no obvious function during development. Unlike other embryonic invaginations such as the ventral furrow, which gives rise to mesodermal precursors, and the midgut invaginations, which give rise to endodermal tissues, the cephalic furrow does not give rise to any specific structure; it simply unfolds after a couple of hours, leaving no trace.^4^ While the cephalic furrow has been thought to serve as a temporary tissue storage,^11^ or as a tissue anchor during gastrulation,^12,13^ these hypotheses have not been investigated *in vivo* or considered in a phylogenetic context. Remarkably, recent evidence indicates that the cephalic furrow is an evolutionary novelty of some dipteran flies,^14^ making it an ideal model for investigating how patterned morphogenetic processes in early embryonic development evolve.

Our work integrates genetics and mechanics to uncover the developmental role and evolutionary origins of the cephalic furrow. First, we analyze how perturbing cephalic furrow formation impacts gastrulation in *Drosophila* by live imaging mutant embryos, and find that the absence of the cephalic furrow increases the mechanical instability of the blastoderm epithelium. Using a combination of *in vivo* experiments and *in silico* simulations, we identify the expansion of mitotic domains and germ band extension as the sources of mechanical stress and show that the cephalic furrow prevents this epithelial instability by absorbing compressive stresses. This suggests the cephalic furrow plays a mechanical role during gastrulation. Next, to uncover the changes in genetic patterning associated with the evolution of the cephalic furrow, we compared the expression of head–trunk genes between species with and without the cephalic furrow. While the expression of the patterning genes is mostly conserved, we find the *btd–eve* overlap, which is required for the cephalic furrow specification in *Drosophila*, is absent in species without the cephalic furrow. This suggests that changes in the expression of *btd* at the head–trunk boundary may be associated with the evolution of the cephalic furrow. Taken together, these data suggest that the evolution of the cephalic furrow patterning system occurred through the cooption of a novel genetic player, and that the underlying selective pressure may have been the mechanical instability during gastrulation. Our findings reveal an interplay between genetic patterning and mechanical forces during the evolution of morphogenesis in early development.

## Analyses of cephalic furrow mutants

To understand the physical consequences of perturbing the formation of the cephalic furrow in *Drosophila*, we investigated the tissue dynamics at the head–trunk boundary in known cephalic furrow mutants—*btd*, *eve*, and *prd*.^8,15^ We generated fluorescent lines carrying a loss of function allele of these genes, and imaged the embryos with high temporal resolution using *in toto* lightsheet microscopy, characterizing differences in timing and dynamics of developmental events during gastrulation (Fig. 1a).

**Figure 1:**
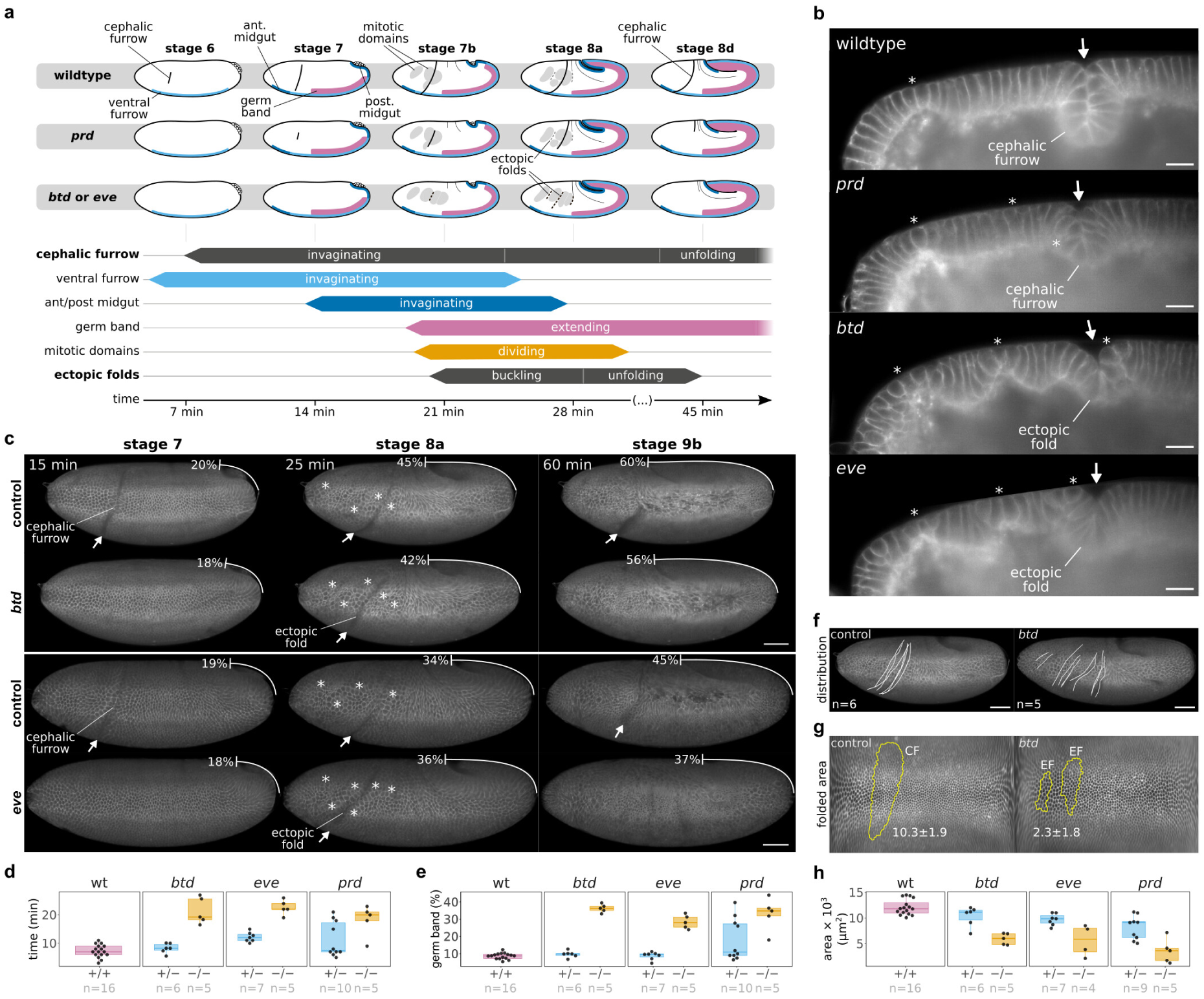
Formation of ectopic folds in cephalic furrow mutants. **a**, Overview of key developmental events in the different genetic backgrounds. Cephalic furrow formation is delayed in *prd* mutants and absent in *btd* and *eve* mutants. Ectopic folds form around the head–trunk boundary, appearing later and unfolding quicker than the cephalic furrow. Their formation coincides with the appearance of mitotic domains and rapid phase of germ band extension.^62^ **b**, Profile views of wildtype and *prd* embryos at early stage 8 and *btd* and *eve* embryos at late stage 8, showing the divergent morphologies of the cephalic furrow and ectopic folds. Scale bars = 20 µm. **c**, Lateral view of sibling controls (heterozygotes) and mutant embryos (*btd* or *eve* homozygotes). Arrows indicate epithelial folds. Asterisks indicate the position of mitotic domains. Percentages indicate the extent of germ band extension relative to egg length. Scale bars = 50 µm. **d**, Timing of formation of cephalic furrow and ectopic folds in different genetic backgrounds. The cephalic furrow forms about 7 minutes after gastrulation in wildtype and *btd* heterozygotes and is delayed in *eve* and *prd* heterozygotes and in *prd* homozygotes. Ectopic folds form about 20 min after gastrulation in *btd* and *eve* homozygotes. Each point represents one embryo. **e**, Percentage of germ band extension at the time of cephalic furrow and ectopic fold formation. The cephalic furrow appears at 10% germ band extension. Ectopic folds form at 30–35% germ band extension. **f**, Variability in the positioning of the cephalic furrow in sibling controls (*btd* heterozygotes) and ectopic folds in *btd* mutants. Scale bars = 50 µm. **g**, Folded area (yellow outline) of the cephalic furrow (CF) in a *btd* heterozygote (left) and of ectopic folds (EF) in a *btd* homozygote (right). Lateral view of a cartographic projection of the embryos. The numbers indicate the average folded area in µm^2^×10^3^ for each condition. **h**, Quantification of the total folded area in cephalic furrow mutants. The membrane marker in panels **b**, **c**, **f**, and **g** is Gap43-mCherry. Staging based on a standard developmental table.^63^

### Disruption of initiator cell behavior

We first analyzed how the behavior of initiator cells is perturbed in the three mutant backgrounds. In wildtype embryos, the initiator cells shorten and undergo anisotropic apical constriction minutes before the end of cellularization^5^ (Extended Data Fig. 1a,b). In *prd* mutants, both behaviors are delayed in initiator cells, and the adjacent cells lack the typical arched profile present in the wildtype invagination^5^ (Extended Data Fig. 1a,b). Moreover, the tissue only invaginates after gastrulation and the resulting fold is abnormal compared to a wildtype invagination. In *btd* and *eve* mutants, the cephalic furrow phenotypes are stronger. Initiator cells in *btd* embryos exhibit a reduced degree of apical constriction but do not shorten (Extended Data Fig. 1a,b, Supplementary Video 1). In contrast, *eve* mutants show neither apical constriction nor cell shortening; the epithelium remains flat for several minutes after gastrulation (Extended Data Fig. 1a,b). These observations reveal that initiator cell behavior in *prd* mutants is not only delayed but also perturbed, and that the cellular mechanism that drives cephalic furrow formation is severely disrupted in *btd* and *eve* mutants.

### Formation of ectopic folds

The absence of initiator shortening in *btd* and *eve* is associated with a strong phenotype: the formation of late epithelial folds near the canonical site of cephalic furrow invagination (Fig. 1b,c, Extended Data Fig. 1a). While the presence of late folds was already observed in *eve* mutants^8^ and, more recently, in *btd* mutants,^9^ their origin and relation to the cephalic furrow have not been investigated. Therefore, to understand the mechanisms causing these epithelial folds, we characterized their formation in different genetic backgrounds.

These ectopic folds, as they will be referred to from hereon, appear around the head–trunk boundary of *btd* and *eve* embryos, about 20 min after gastrulation when the germ band is already extended 35% of the egg length (Fig. 1a–e, Supplementary Video 2, Supplementary Video 3, Table 1). Ectopic folds can superficially resemble a cephalic furrow, but they lack the typical symmetric morphology of the wildtype invagination.^5^ Instead, ectopic folds in *btd* and *eve* mutants have a loose, often asymmetric cleft without wedge- or arched-shaped cells (Fig. 1b, Supplementary Video 4, Supplementary Video 5), and occupy ^1^ of the area and ^1^ of the depth of the cephalic furrow (Fig. 1g,h, Extended Data Fig. 2b,f, Table 2, Table 3). Ectopic folds also fold and unfold faster, lacking the typical kinetics of cephalic furrow formation (Fig. 1a,b, Extended Data Fig. 1a, Extended Data Fig. 2c–e, Supplementary Video 6). Finally, the formation of ectopic folds is more variable than that of the cephalic furrow, as their position along the head–trunk boundary differs between individual mutant embryos (Fig. 1f, Extended Data Fig. 2a, Supplementary Fig. 1, Supplementary Video 7, Supplementary Video 8). Altogether, these differences in morphology, kinetics, timing, position, and variability, suggest that ectopic folds and cephalic furrow form via distinct mechanisms.

**Table 1:**
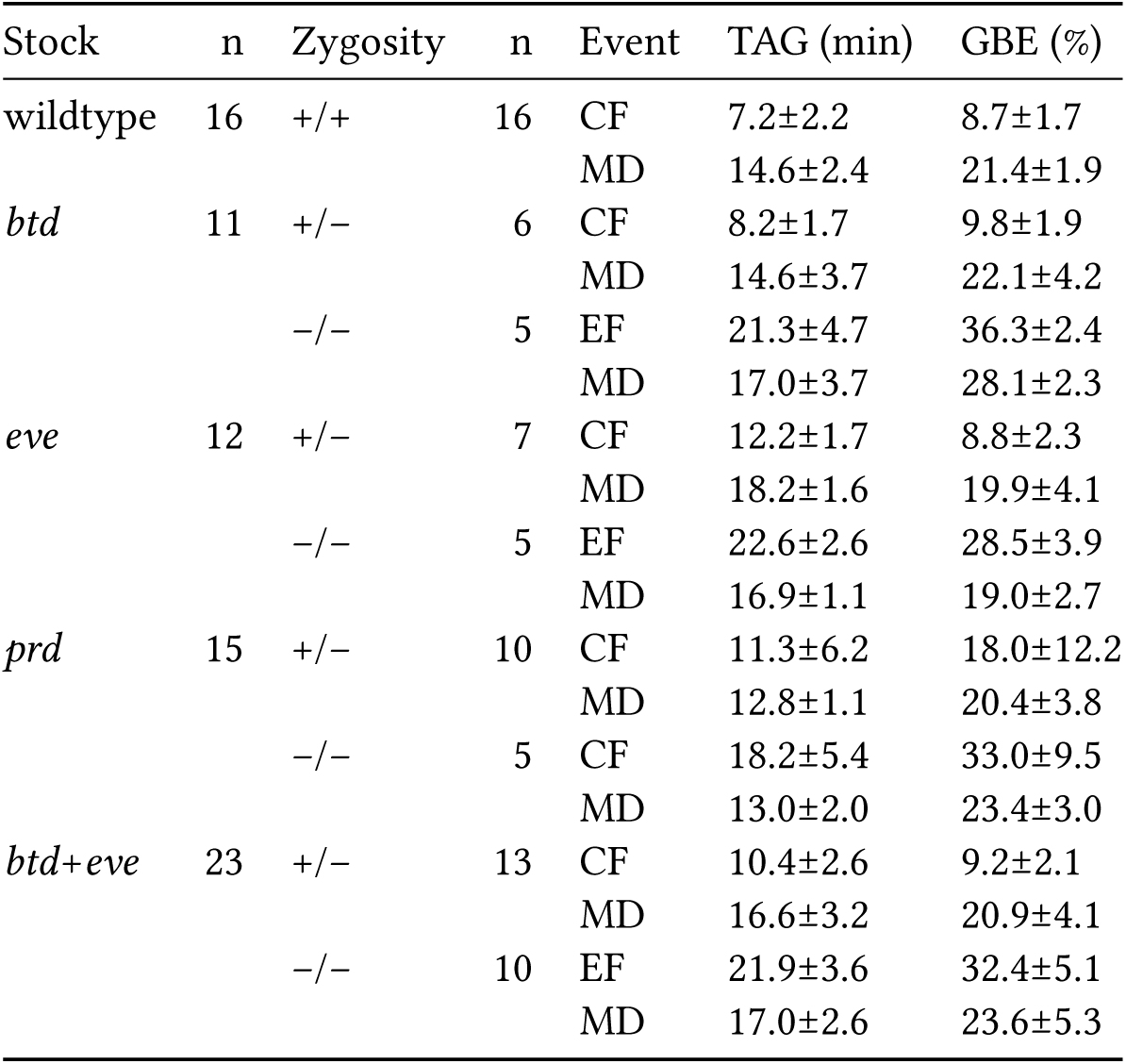
Relative timing differences between developmental events in cephalic furrow mutants. We measured the time after gastrulation (TAG) and the percentage of germ band extension (GBE) at the moment of formation of the cephalic furrow (CF), mitotic domains (MD), and ectopic folds (EFs). For a more general comparison, we also pooled the data for mutants where the cephalic furrow is absent (*btd*+*eve*).

**Table 2:**
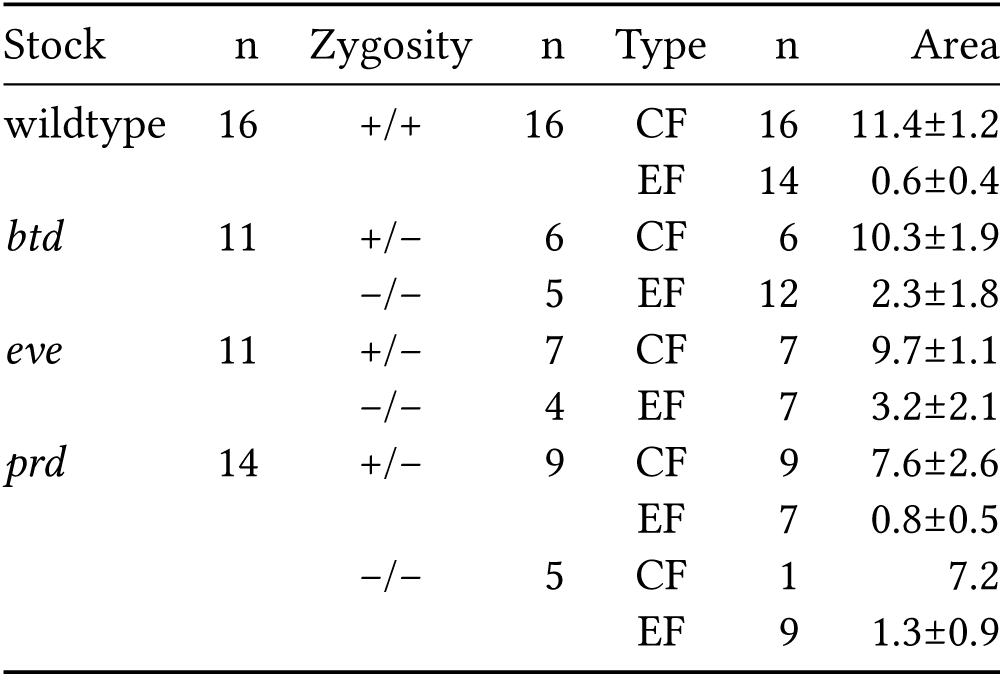
Surface area of ectopic folds in cephalic furrow mutants. Values in µm^2^×10^3^.

**Table 3:** Maximum depth of ectopic folds in cephalic furrow mutants. Values in µm.

| Stock | n | Zygoty | n | Type | n | Depth |
| --- | --- | --- | --- | --- | --- | --- |
| *btd* | 39 | +/- | 32 | CF | 52 | 71.6±8.0 |
| | | | | EF | 6 | 32.5±3.6 |
| | | -/- | 7 | EF | 28 | 52.5±12.1 |
| *eve* | 24 | +/- | 20 | CF | 34 | 59.0±6.8 |
| | | | | EF | 4 | 36.1±4.4 |
| | | -/- | 4 | EF | 15 | 42.1±11.7 |

When applying our ectopic folding scoring pipeline, we noted that heterozygote and wildtype embryos can also form ectopic folds located anterior or posterior to the cephalic furrow. The frequency of ectopic folding in these embryos is lower than in homozygotes, but not negligible. While nearly every *btd* and *eve* homozygote embryos (>92%) show one or more ectopic fold per side (2.2±0.4 and 1.8±0.6, respectively) (Extended Data Fig. 2h, Supplementary Video 8), between 18–27% of heterozygotes and about 78% of wildtype embryos have one ectopic fold in addition to the cephalic furrow (Extended Data Fig. 2i,j, Table 4). However, we find a fundamental difference. The area of the ectopic folds in heterozygote and wildtype embryos is significantly smaller (about 4x) compared to the area of ectopic folds in *btd* and *eve* embryos (Extended Data Fig. 2g,k–n, Table 2). This data shows the head–trunk interface of *Drosophila* during gastrulation is a region prone for the formation of ectopic folds and provides evidence that the absence of the cephalic furrow leads to an increase in the magnitude of these folding events.

**Table 4:** Folding statistics in cephalic furrow mutants. We calculated the percentage of embryos having a cephalic furrow (CF) or ectopic folds (EF) for each stock and genotype (Frequency), including the position of folding along the head–trunk boundary (Ant=anterior, Mid=middle, Post=posterior). In addition, we calculated the average number of folds per embryo side (Folds). For example, 28 out of 36 wildtype embryos show ectopic folds (77.8%); 42.9% of these embryos have folds at the anterior region and 71.4% form them at posterior to the head–trunk boundary; each embryo forms 1.1±0.3 folds on each side. The *n* includes datasets imaged from the lateral and dorsal sides.

| Stock | n | Zygoty | n | Type | n | Frequency | Ant | Mid | Post | Folds |
| --- | --- | --- | --- | --- | --- | --- | --- | --- | --- | --- |
| wildtype | 36 | +/+ | 36 | CF | 36 | 100% | 0% | 100% | 0% | $1.0 \pm 0.0$ |
| | | | | EF | 28 | 77.8% | 42.9% | 0% | 71.4% | $1.1 \pm 0.3$ |
| *btd* | 46 | +/- | 33 | CF | 33 | 100% | 0% | 100% | 0% | $1.0 \pm 0.0$ |
| | | | | EF | 6 | 18.2% | 0% | 0% | 100% | $1.0 \pm 0.0$ |
|  |  | -/- | 13 | CF | 0 | 0% | 0% | 0% | 0% | - |
| | | | | EF | 12 | 92.3% | 50% | 100% | 75% | $2.2 \pm 0.4$ |
| *eve* | 36 | +/- | 26 | CF | 26 | 100% | 0% | 100% | 0% | $1.0 \pm 0.0$ |
| | | | | EF | 7 | 26.9% | 14.3% | 0% | 85.7% | $1.0 \pm 0.0$ |
|  |  | -/- | 10 | CF | 0 | 0% | 0% | 0% | 0% | - |
| | | | | EF | 10 | 100% | 40% | 70% | 90% | $1.8 \pm 0.6$ |
| *prd* | 40 | +/- | 26 | CF | 26 | 100% | 0% | 100% | 0% | $1.0 \pm 0.0$ |
| | | | | EF | 7 | 26.9% | 71.4% | 0% | 57.1% | $1.3 \pm 0.5$ |
| | | -/- | 14 | CF | 7 | 50.0% | 0% | 100% | 0% | $1.0 \pm 0.0$ |
| | | | | EF | 10 | 71.4% | 50% | 80% | 70% | $1.9 \pm 0.8$ |
| *stg* | 46 | +/- | 33 | CF | 33 | 100% | 0% | 100% | 0% | $1.0 \pm 0.0$ |
| | | | | EF | 12 | 36.4% | 8.3% | 0% | 91.7% | $1.0 \pm 0.0$ |
| | | -/- | 13 | CF | 13 | 100% | 0% | 100% | 0% | $1.0 \pm 0.0$ |
| | | | | EF | 3 | 23.1% | 0% | 0% | 100% | $1.0 \pm 0.0$ |

### Evidence of tissue compression

The variability in ectopic folding suggests that, unlike the cephalic furrow, the ectopic folds are not under genetic control and form as the result of physical interactions in the tissue. Our analysis shows that the formation of ectopic folds coincides spatially and temporally with two other processes of gastrulation: the expansion of mitotic domains and the extension of the germ band (Fig. 1a,c,d,e).

Mitotic domains are groups of blastoderm cells that divide in synchrony during nuclear cycle 14, first appearing 20 min after gastrulation on the embryo’s head.^6^ Our analysis of the ectopic folds in *btd* and *eve* mutants shows that they form between mitotic domains or between the posterior Mitotic Domain 6 (MD6) and the extending germ band (Fig. 2a,b). When mitotic cells begin to divide, they lose their basal attachment, round up at the apical side, and more than double their apical area during anaphase (Extended Data Fig. 3a). This apical expansion compresses the adjacent, non-dividing cells, which are the first to fold inward (Fig. 2b). Mitotic expansions always precede ectopic folding (Fig. 2c,d, Extended Data Fig. 1a, Table 1). This suggests the formation of mitotic domains may generate buckling instability in the monolayer epithelium, contributing to the appearance of ectopic folds.

**Figure 2:**
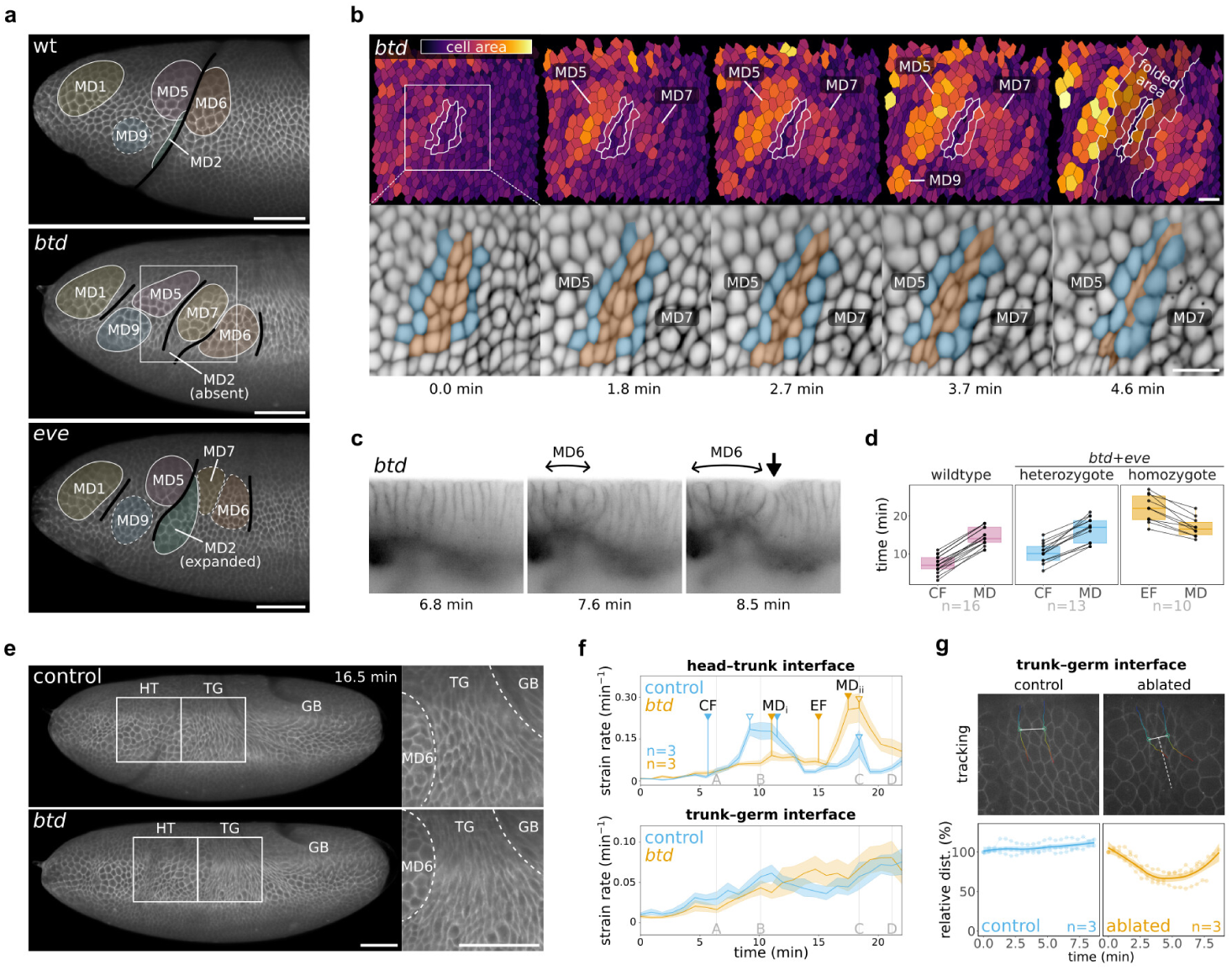
Coincidence of mitotic domains and germ band extension with ectopic folding. **a**, Position of folding (black lines) in relation to mitotic domains (colored areas) in wildtype embryos and *btd* and *eve* mutants. Ectopic folds appear between mitotic domains and between the germ band and the mitotic domain 6 (MD6). Scale bars = 50 µm. **b**, Apical area of cells between the mitotic domains MD5 and MD7 (top) until the formation of a ectopic fold; folded area is highlighted at the last frame (4.6 min). The bottom panels show an annotated subset of non-dividing cells (orange) and dividing cells (blue). Scale bars = 20 µm (approximate value since pixel resolutions vary across the projection). **c**, Ectopic folding between MD6 and germ band (not visible) in a *btd* mutant. Mitotic expansions (double arrow) always precede ectopic folding (arrow). Scale bar = 20 µm. **d**, Comparison between the timing of cephalic furrow formation (CF), mitotic domain expansion (MD), and ectopic folding (EF) in wildtype and cephalic furrow mutants (*btd* and *eve* combined). In wildtype and heterozygote embryos, the cephalic furrow forms before mitotic domains, while in homozygote embryos, the ectopic folds form after mitotic domains. **e**, Lateral view of sibling control and *btd* mutant showing the head–trunk (HT) and trunk– germ (TG) regions. The apical surface of cells at the TG, between MD6 and the tip of the germ band, exhibits a compressed profile. Scale bars = 50 µm. **f**, Strain rate analysis at the head–trunk (top) and trunk–germ (bottom) regions. Filled triangles indicate the beginning of cephalic furrow (CF), mitotic domains (MD), and ectopic folds (EF) formation. MD_i_ shows the initial metaphase expansion and MD_ii_ the moment of cell division during telophase. Empty triangles indicate the peaks of strain rate. Letters A–D show the exact frames from Extended Data Fig. 3b. The measurements combine isotropic and anisotropic strain rate. **g**, Dynamics of trunk–germ tissues after laser cuts in wildtype. We measured the distance between pairs of cell vertices over time in control and ablated embryos. Tracks are color-coded for time. Solid white line represents the distance between vertices at one timepoint. Dashed white line shows the location of the cut. The plots show a smoothed local regression of three samples.

To estimate the forces acting on the buckling tissue, we measured the rate of tissue deformation (strain rate) at the head–trunk and trunk–germ boundaries using particle image velocimetry (Fig. 2e). Control embryos exhibit a peak of strain rate around the head–trunk interface that correlates with late phase of the cephalic furrow invagination, when the initiator cells move into the yolk (Fig. 2f, Extended Data Fig. 3b, Supplementary Video 9). This is absent in *btd* mutants. Instead, the mutants show a higher peak of strain rate that coincides with the maximum expansion of the mitotic domains during telophase and ectopic folding (Fig. 2f, Extended Data Fig. 3b, Supplementary Video 9). This suggests the tissue deformation is associated with the mitotic expansions.

At the trunk–germ interface, the cells between MD6 and the extending germ band become increasingly anisotropic (Fig. 2e). We find that the strain rate in this region steadily increases over time (Fig. 2f), suggesting that the tissue is under compression. To test this hypothesis, we performed laser cuts at the trunk–germ interface in wildtype embryos. We ablated the apical membrane of multiple cells (3– 4) with cuts oriented orthogonal to the direction of the germ band extension, and then tracked the distance between non-ablated cells on each side of the cut (Fig. 2g). This distance remains constant in uncut control embryos, but decreases over the first five minutes after the cut (Fig. 2g). This is evidence that the tissue may be “collapsing on itself” after the ablation, which supports the hypothesis that the trunk–germ interface is under compression from the extending germ band.

Taken together, these analyses suggest the mitotic expansions and germ band extension are potential sources of compressive stress capable of inducing tissue buckling, and thus could contribute to the formation of ectopic folds at the head–trunk boundary during gastrulation.

### Contribution of mitotic domains and germ band

To test the individual role of mitotic domains and germ band to the mechanical stability of the blastoderm, we performed a series of perturbation experiments *in vivo*.

We first asked if mitotic expansions are required for the formation of ectopic folds. To that end, we generated a double-mutant line lacking both cephalic furrow and mitotic domains, using the loss-of-function alleles of *btd* and *string* (*stg*)—the *cdc25* phosphatase ortholog that regulates the formation of mitotic domains in *Drosophila*.^16^ The absence of mitotic domains in *stg* mutants does not affect the formation of the cephalic furrow or other morphogenetic movements of gastrulation^16^ (Fig. 3a, Extended Data Fig. 4a,b, Supplementary Video 10, Supplementary Video 11). However, the absence of mitotic domains suppresses the formation of ectopic folds in *btd*–*stg* double-mutants; these embryos show no ectopic folds at their head–trunk interface (Fig. 3b,c, Supplementary Video 12, Supplementary Video 13). This indicates that mitotic expansions are necessary for the appearance of ectopic folds in cephalic furrow mutants.

**Figure 3:**
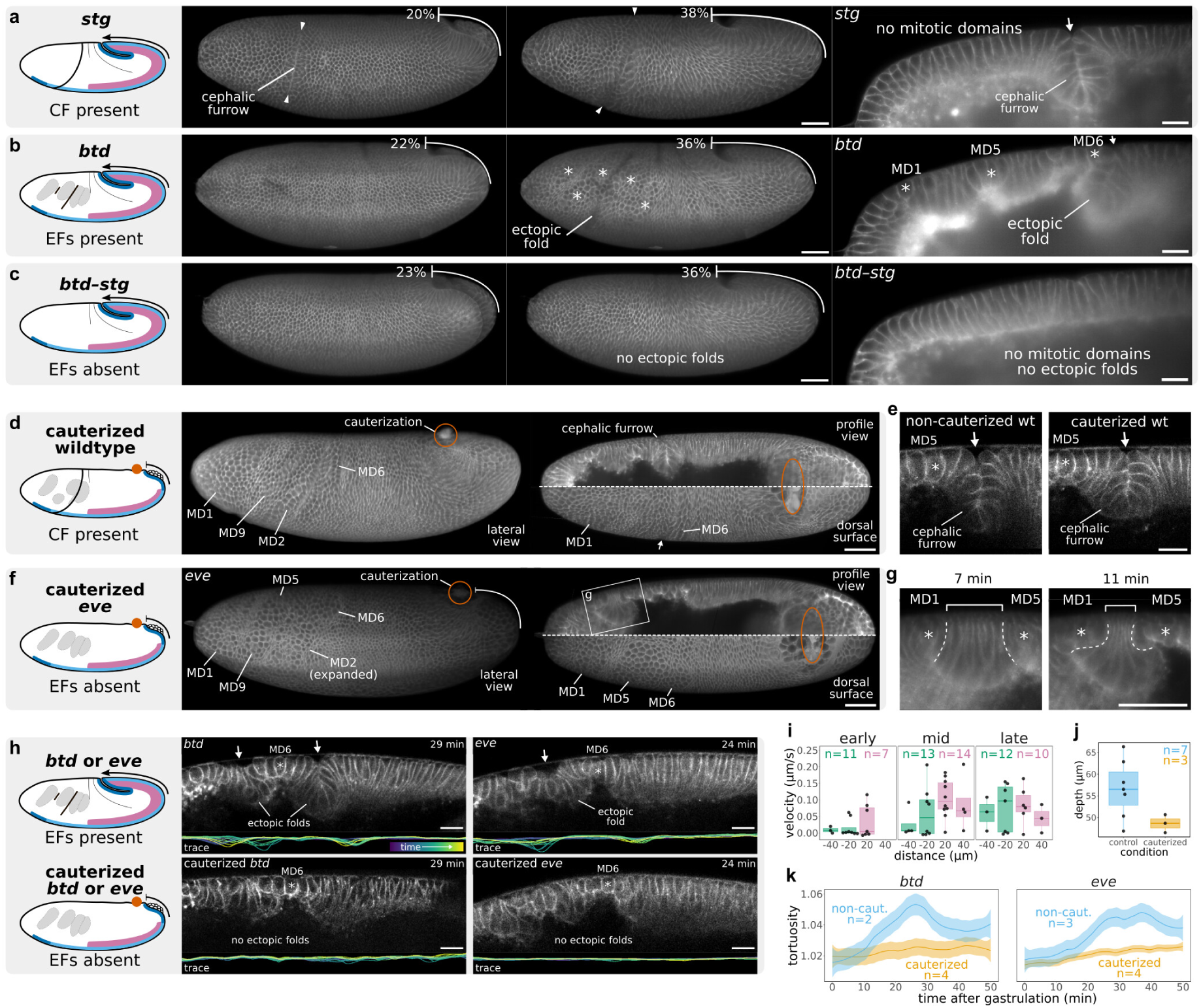
Perturbation experiments *in vivo* using cephalic furrow mutants and wildtype embryos. **a**, Lateral and profile views of *stg* mutants showing the formation of the cephalic furrow and germ band extension are unaffected. Scale bars = 50 µm and 20 µm. **b**, Lateral and profile views of *btd* mutants showing the formation of ectopic folds. Scale bars = 50 µm and 20 µm. **c**, Lateral and profile views of *btd*–*stg* double mutants, which lack both the cephalic furrow and mitotic domains, showing the absence of ectopic folds at the head–trunk interface. Scale bar = 50 µm and 20 µm. **d**, Cauterized wildtype embryo in lateral (left) and dorsal view (right). Cauterization site is outlined in orange. Cephalic furrow formation occurs normally. Scale bar = 50 µm. **e**, Profile view of a non-cauterized and cauterized wildtype embryos showing a small difference in depth. Scale bar = 20 µm. **f**, Cauterized *eve* mutant in lateral (left) and dorsal view (right) at the maximum expansion of dividing cells in mitotic domains. Cauterization site is outlined in orange. The boxed outline (right) shows panel **g**. Scale bar = 50 µm. **g**, Inset from panel **f** showing the progressive expansion of mitotic domains compressing the non-dividing cells between MD1 and MD5. Scale bar = 50 µm. **h** Profile view and epithelial trace of non-cauterized and cauterized *btd* and *eve* embryos. The trace shows the dynamics of epithelial deformations over time, colored from purple to yellow. Cauterized mutants show no ectopic folds and less deformation of the epithelium. Scale bars = 20 µm. **i**, Recoil velocity of laser ablations around the cephalic furrow at stage 6. The average recoil increases with time for anterior cuts, and reaches a peak at mid-stage 6 for posterior cuts. The average recoil velocity is higher at shorter distances from the initiator cells (−20 and 20 µm) and lower at greater distances (−40 and 40 µm). In late stage 6, we detect recoil within 40 µm anterior and posterior of the invagination. **j**, Maximum depth of the cephalic furrow in cauterized embryos. We pooled wildtype and heterozygote embryos for non-cauterized (wt=2, *btd*=1, and *eve*=3) and cauterized (wt=2 and *btd*=1) samples. The cephalic furrow in cauterized embryos is 15% shallower (p=0.0221 in Welch Two Sample t-test). **k**, Tortuosity of the epithelial traces in non-cauterized (*btd*=2, *eve*=3) and cauterized (*btd*=3, *eve*=4) embryos from **h**. We measured left and right sides of each embryo.

Next, we asked if the extension of the germ band is required for the formation of ectopic folds. To prevent the germ band from extending, we cauterized a patch of posterodorsal tissue at the onset of gastrulation to mechanically attach it to the vitelline envelope (Fig. 3d,f). Blocking the germ band in wildtype embryos does not prevent the formation of the cephalic furrow (Fig. 3d,e), as the shortening of the initiator cells occurs normally and correlates with an increase in tension around the head– trunk boundary (Fig. 3i). We only detect a mild influence of the germ band on the final depth of the invagination, since the cephalic furrow in cauterized embryos is about 15% shallower than in control embryos (Fig. 3e,j). This data corroborates the understanding that the initial formation of the cephalic furrow is an active process independent of other morphogenetic movements

We then performed the cauterization experiment in *btd* and *eve* embryos. When the germ band extension is blocked in these mutants, no ectopic folds appear at the head–trunk interface (Fig. 3f, Supplementary Video 14, Supplementary Video 15, Supplementary Video 16), and their epithelium undergoes less deformation and buckling events compared to non-cauterized mutant embryos (Fig. 3h,k). The expansion of cell apices in mitotic domains still compress the neighboring, non-dividing cells, but no buckling occurs (Fig. 3g). These experiments reveal that the germ band extension is necessary for the appearance of ectopic folds in cephalic furrow mutants.

Overall, we conclude that neither mitotic expansions nor the germ band extension can induce ectopic folds individually, but when both events happen concomitantly, the epithelial monolayer becomes unstable and buckles.

## Physical model of folding dynamics

To determine the relative contribution of mitotic domains and germ band as sources of mechanical stress on the head–trunk boundary, we created a physical model of the blastoderm and simulated the tissue mechanics of mutant and wildtype conditions *in silico*.

### Model and simulation design

Our model represents an epithelial monolayer confined inside a rigid shell. It embodies one side of a frontal slice between the midline and the dorsal apex of a *Drosophila* embryo with its typical morphological proportions (Fig. 4a, Fig. Extended Data Fig. 5a). The blastoderm is modeled by an elliptical arc of equidistant particles connected by springs and enclosed on one side by a rigid barrier representing the vitelline envelope (Fig. 4b). The total energy per unit length of this tissue (*W_T_*) is a sum of a stretching energy component (*W_s_*) and a bending energy component (*W_b_*) (Fig. 4c). Each of these components has a rigidity associated with them. *K_s_* is the stretching rigidity and *K_b_* is the bending rigidity. These two parameters can be combined into a single dimensionless bending rigidity, 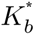 (Fig. 4c).

**Figure 4:**
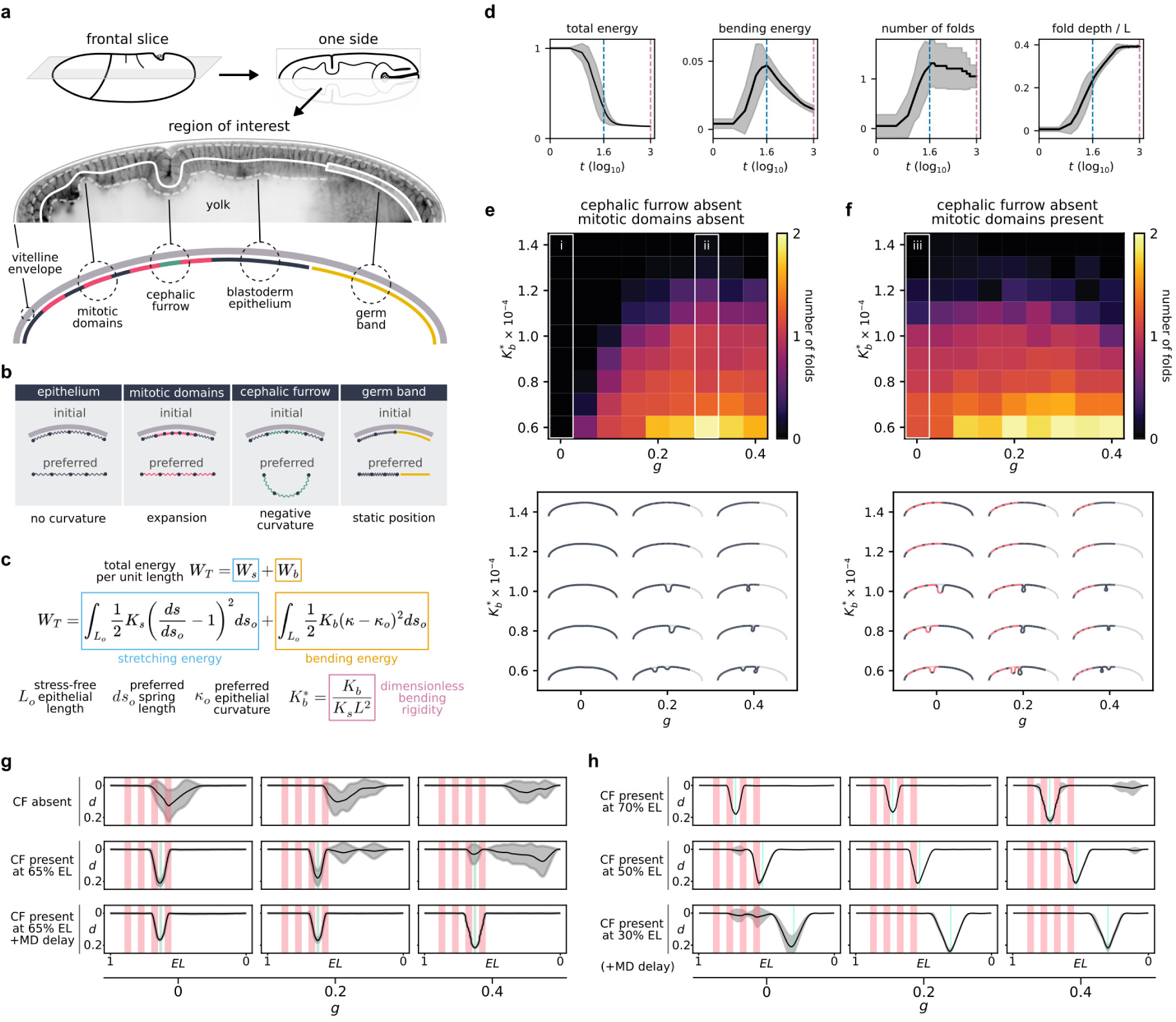
Model and simulations of the tissue mechanics at the head–trunk boundary. **a**, Region of interest of the model. From a frontal slice across a dorsal plane, we select one side of the blastoderm, following the embryo’s shape, proportions, and relative position of mitotic domains, cephalic furrow, and germ band. **b**, Characteristics of the model components which is based on particles connected by springs. Drawings exemplify the initial and final states for each component. **c**, Energy equation with a stretching and a bending component, and the dimensionless bending rigidity. Stress-free rod length (*L_o_*), total energy per unit length (*W_T_*), stretching energy per unit length (*W_s_*), bending energy per unit length (*W_b_*), stretching rigidity (*K_s_*), bending rigidity (*K_b_*), preferred spring length (*d_so_*), current spring length (*d_s_*), preferred curvature (*κ_o_*), current curvature (*κ*), semimajor embryonic axis (*L*). **d**, Plots showing the energy dynamics across iterations in a typical simulation. Total energy goes down to equilibrium. Bending energy increases drastically reaching a peak (blue dashed line) that diminishes gradually with each iteration. Energy values are normalized by the initial total energy. The number of folds stabilizes when the bending energy peaks but the fold depth continues to increase until the last iteration (pink dashed line). **e**, Parameter sweep for a mutant condition (no cephalic furrow) without mitotic domains. The heatmap shows the average number of ectopic folds for different bending rigidities and percentages of germ band extension. Ectopic folding frequency increases with lower bending rigidities (softer tissues) and with greater values of germ band extension. Outlined in white are the baseline conditions with neither mitotic domains nor germ band but only ground level noise (i), and the germ-band-only condition with more folding events (ii). A visual rendering of representative simulations are shown below. **f**, Parameter sweep for a mutant condition (no cephalic furrow) with mitotic domains. The phase diagram shows an increase in number of folds in relation to **e**. The addition of mitotic domains induces the formation of ectopic folds even without germ band extension (iii). Representative simulations are shown below. **g**, Quantification of ectopic folding to evaluate the effectiveness of the cephalic furrow. A mutant condition with mitotic domains and germ band extension show extensive ectopic folding (top row). With the presence of the cephalic furrow, ectopic folding decreases around the head–trunk region, especially in lower percentage of germ band extension (middle row). Adding a delay of *t_MD_* = 5 to mitotic domain formation to mimic the *in vivo* condition results in less frequent ectopic folding throughout the blastoderm and germ band extension (bottom row). *t_MD_* = 1 corresponds to 105 computational timesteps. **h**, Quantification of ectopic folding with the cephalic furrow at different positions along the anteroposterior axis. Displacing the cephalic furrow more anteriorly results in ectopic folding at the posterior end (top row) while displacing it more posteriorly results in more ectopic folding at the anterior region (bottom row). Placing the cephalic furrow around the central region prevents ectopic folding more broadly.

To simulate the physical interactions between mitotic domains, germ band, and cephalic furrow, we defined the mitotic domains as compressed regions which tend to expand (they contain more particles compared to the surrounding regions), and the cephalic furrow as a narrow region having an intrinsic negative curvature predisposing the tissue to invaginate (Fig. 4b). The germ band is defined by the position of the posterior end of the tissue (*g*), which is fixed at different fractions of egg length for each simulation (Fig. 4a,b). Thus, the effect of germ band extension appears as a global compression in the blastoderm. To run the simulations, we defined a ground level of random noise and iterated towards equilibrium of the total energy in the system (see Methods). We then set several simulations with different bending rigidity values and combinations of presence/absence of mitotic domains, cephalic furrow, and percentages of germ band extension to quantify the position, frequency, and depth of ectopic folding in the epithelium.

### General properties and energy dynamics

To characterize the model properties and energy dynamics, we ran simulations using initially a single bending rigidity value, without mitotic domains, and at different percentages of germ band extension. Without the germ band, the tissue is almost stress-free and no ectopic folding occurs (Fig. 4e). We begin to observe folds in the simulations with higher progression of germ band extension. Folding events are stochastic and happen at distinct iterations for each simulation. When a fold begins to form, the bending energy increases releasing a larger amount of stretching energy, which, in turn, decreases the total energy of the system over each iteration (Fig. 4d). The increase in bending energy coincides with a rapid deepening of the fold. Once the bending energy reaches a peak, we find that the fold continues to deepen more gradually, but the number of folds rarely changes afterward (Fig. 4d, Extended Data Fig. 5b). Therefore, this peak of bending energy provides an informative reference point, which we used to standardize the comparison across simulations.

### Bending rigidity sweep in mutant conditions

To obtain realistic values of the dimensionless bending rigidity 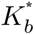, we performed a sweep across the parameter space for the cephalic furrow mutant conditions. In general, we find that the number of ectopic folds is higher in softer conditions, where the bending rigidity is lower (Fig. 4e,f, Extended Data Fig. 5c,f). In simulations without mitotic domains, we observe no folding events without the germ band (Fig. 4e, column i). With the extension of the germ band, the probability of buckling increases (Extended Data Fig. 5d) and the time to folding decreases (Extended Data Fig. 5e). The parameter sweep shows a clear transition in the phase space with the buckling probability reaching a plateau around 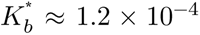 (Fig. 4e). In these stiffer conditions, the germ band, even at its maximum extension, cannot drive the formation of ectopic folds.

Adding mitotic domains to the simulations changed the phase diagram, increasing the probability of folding. We find that, in softer conditions 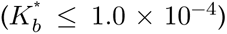, mitotic domains alone can induce ectopic folding (Fig. 4f, column iii). For lower values of germ band extension, the number of folds is higher (Extended Data Fig. 5g) and the time to folding is lower (Extended Data Fig. 5h) compared to the simulations without mitotic domains. These simulations reveal that, depending on the bending rigidity, the germ band or mitotic domains alone can drive ectopic folding, and that their combined action increases the mechanical instability of the tissues.

To determine where our embryo lies in this parameter space, we considered the insight from our experimental data that neither mitotic domain nor germ band can promote ectopic folding by themselves. With this information, we identified the bending rigidity value that recapitulates these biological behaviors in the simulations. For that, we determined at which bending rigidity the average number of folds falls below 1 in mitotic-domains-only and germ-band-only conditions (Extended Data Fig. 5c,f). The criterion is fulfilled when the bending rigidity is 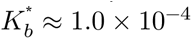.

To compare this value with existing direct measurements, we calculated * for the bending rigidity estimates in cultured cells ((*K_b_* ≈ 5 × 10^−13^*Nm* and *K_s_* ≈ 0.2*Nm*^−1^) ^17,18^ (Supplementary Note 1). Given that in elastic sheets the bending and stretching rigidity ratio is proportional to the square of the height (*K_b_*/*K_s_* ∝ *h*_2_),^19^ we adjusted the calculation to the thickness of the epithelial monolayer in culture (∼18 µm).^18^ The value we obtained in this way for the effective 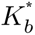 of a height-adjusted epithelial monolayer, ≈ 1.8 × 10^−4^, is within a factor of 2 of our own reference bending rigidity, indicating that the relevant region of parameter space we identified in our model is consistent with existing direct measurements of bending rigidity in other tissues. This value establishes an important bridge between our *in vivo* and *in silico* data by revealing where the embryo resides within the parameter space of the model, and by providing a biologically relevant reference for subsequent simulations.

### Impact of cephalic furrow on ectopic folding

The formation of ectopic folds in cephalic furrow mutants led us to hypothesize that the wildtype invagination may contribute to absorbing the compressive stresses generated by mitotic domains and germ band in normal conditions. To explore the role of the cephalic furrow as a mechanical buffer, we analyzed how the presence of the invagination affects the dynamics of ectopic folding in the simulations.

We programmed the cephalic furrow in our model by establishing an intrinsic negative curvature 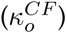 to a narrow region of the particle–spring blastoderm that matches the span of the initiator cells *in vivo* (Fig. 4a,b). Using our reference bending rigidity value 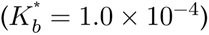, we ran a parameter sweep for different 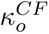 and established a baseline 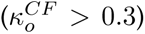 where the invagination forms in a robust manner with minimal variability, phenocopying the cephalic furrow *in vivo* (Extended Data Fig. 5i).

We first evaluated how the strength of the cephalic furrow impacts the frequency and placement of ectopic folding. In conditions without the germ band (*g*=0), we find that an active invagination reduces the spread and frequency of ectopic folding at the head–trunk boundary (Fig. 4g, Extended Data Fig. 5l) and that this reduction in variability correlates with the strength of the pull (Extended Data Fig. 5j). In conditions with an extended germ band, however, this buffering effect was diminished due to an increase in ectopic folding at the posterior region, which dominates the landscape and even inhibits the formation of the cephalic furrow in embryos with 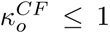 and *g* ≥ 0.02 (Extended Data Fig. 5j). Nevertheless, in general, these simulations show that higher values of 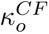 are more effective at preventing ectopic folds.

Since the cephalic furrow forms around 15 min before mitotic domains in wildtype embryos, we asked whether their relative timing of formation may influence the effectiveness of the cephalic furrow. In contrast to the simulations above where the cephalic furrow and mitotic domains form at the same time, we added a delay to the formation of mitotic domains to match the *in vivo* characteristics. In this condition, the cephalic furrow is more effective in preventing ectopic folding, even for lower 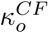 values (Fig. 4g, Extended Data Fig. 5j,k,l). Similar to our *in vivo* observations of ectopic folding in wildtype embryos (Extended Data Fig. 2i–m, Table 4), we also observe some ectopic folding in these simulations, especially at greater percentages of germ band extension (Extended Data Fig. 5k). These experiments reveal that relative timing, rather than force, is more important at preventing ectopic folding.

Finally, we tested how the position of the cephalic furrow impacts its ability to prevent ectopic folding. When the germ band is not extended (*g* = 0.0), a cephalic furrow positioned anteriorly (>70% embryo length) effectively prevents ectopic folding in the head (Fig. 4h), similar to the simulations at the wildtype position. However, when the cephalic furrow is positioned more posteriorly (<50% embryo length), ectopic folding around mitotic domains becomes more frequent. Conversely, when the germ band is extended (*g* = 0.4), a posteriorly located cephalic furrow (<30% embryo length) prevents ectopic folding in the posterior region, while an anteriorly located invagination (>50% embryo length) fails to do so (Fig. 4h). These simulations show that when the cephalic furrow is positioned between 40-60% embryo length, it is the most effective at preventing ectopic folding throughout the different phases of germ band extension.

Taken together, our physical model provides a theoretical basis that a fold such as the cephalic furrow— by forming before other morphogenetic events around the central region of the anteroposterior axis— can absorb compressive stresses and prevent, to a substantial degree, mechanical instabilities in embryonic tissues during gastrulation.

## Evolution of gene expression

As described above, our analyses suggest that the effectiveness of the cephalic furrow in preventing epithelial instabilities depends on the position and time of the invagination. In *Drosophila*, this spatiotemporal control is determined genetically by the combinatorial expression of *btd*, *eve*, and *prd* at the head–trunk boundary.^8,15^ However, it remains unclear which elements of this genetic patterning cascade are associated with the evolution of the cephalic furrow. To address this question, we first sought to identify additional genes involved in cephalic furrow formation, and then asked how the expression of these cephalic furrow genes compare between dipteran species with and without the cephalic furrow.

### Role of *sloppy paired* in cephalic furrow patterning

To identify other cephalic furrow genes, we performed a live-imaging screen using loss-of-function alleles of several candidates expressed at the head–trunk interface (Supplementary Note 2). We found that null mutants for the *sloppy paired* (*slp*) transcription factors, *slp1* and *slp2*, show a strong phenotype where the cephalic furrow is delayed and shifted towards the anterior end by ∼6% of the egg length (control = 67.6±1.4%, n=26; *slp* = 73.2±0.7%, n=7) (Fig. 5a,b, Supplementary Video 17). With this anterior shift, *slp* mutants exhibit a more prominent posterior dorsal fold and an early ectopic fold within MD6 appearing before cell divisions (Fig. 5a, Supplementary Video 17). These observations are congruent with the increase in posterior mechanical instability present in our simulations where the cephalic furrow is shifted forward (Fig. 4h).

**Figure 5:**
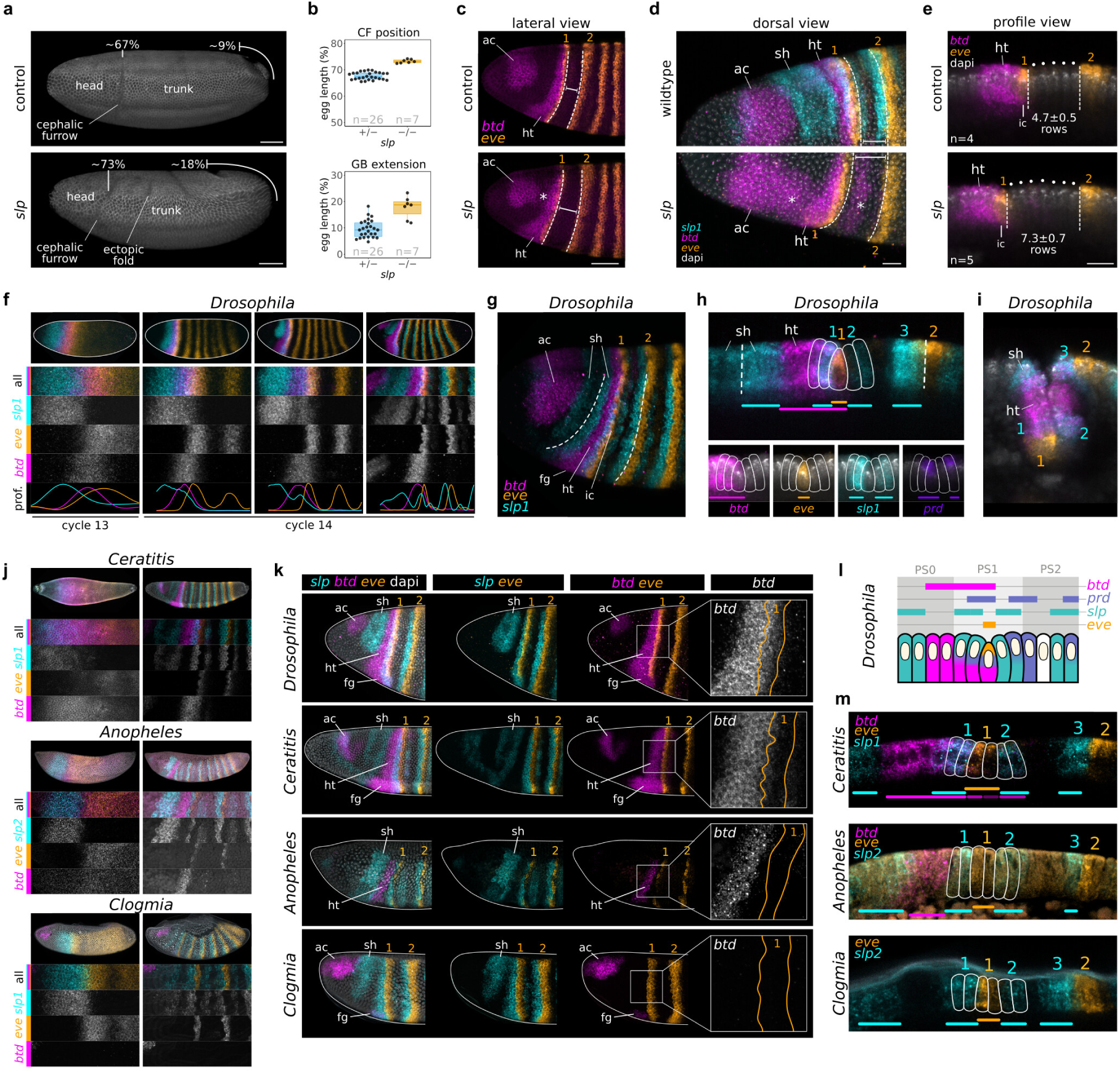
Genetic patterning of the head–trunk boundary in *Drosophila* and *Clogmia*. **a**, Lateral view of *slp* heterozygote (control) and mutant embryo at the onset of initiator cell behavior. In *slp* mutants, the cephalic furrow formation is delayed; it happens when the germ band is extended about 18% of egg length. The position of initiator cells is shifted forward in mutants to about 73% of egg length. The asterisk indicates mitotic cells. Scale bars = 50 µm. **b**, Plots showing the position of the cephalic furrow (CF) and germ band (GB) at the onset of initiator cell behavior in *slp* mutants. **c**, Lateral view of *slp* mutants in *Drosophila* embryos showing the expression of *btd* and *eve*. The distance between *eve* stripe 1 and 2 is larger in *slp* embryos. The asterisk indicates a region between the acron (ac) and head–trunk (ht) domains of *btd* where its expression is activated (or derepressed) in the absence of the *slp* head domain (sh). Scale bars = 50 µm. **d**, Dorsal view showing the ectopic expression of *btd* (asterisks) and anterior shift of *btd*–*eve* overlap in *slp* mutants. Scale bars = 20 µm. **e**, Profile view showing the increased number of cell rows between *eve* stripe 1 and 2 in *slp* mutants. Scale bars = 20 µm. **f**, Progression of the early expression of *btd*, *eve*, and *slp1* around the nuclear cycle 14 in *Drosophila*. *slp1* and *eve* initially demarcate the head–trunk boundary in broad domains which become segmented, narrow and resolve into sharp, non-overlapping stripes. *btd* transcripts are localized at the head–trunk interface. **g**, Lateral view of the expression at the head–trunk boundary in *Drosophila*. *slp1* stripes demarcate the outer edges of the cephalic furrow (dashed lines). **h**, Profile view of the gene expression in initiator cells of *Drosophila*. The *eve*-expressing row is abutted anteriorly and posteriorly by non-overlapping *slp1* stripes. *btd* and *eve* overlap by 1-cell row. *prd* expression is offset with *slp1*. Dashed lines mark the future edges of the cephalic furrow. **i**, Profile view of the gene expression after cephalic furrow invagination in *Drosophila*. The *eve*-expressing initiator cells are located deep into the yolk, while the *slp* head domain (sh) and stripe 3 demarcate the outer edges of the invagination. **j**, Expression domains during nuclear cycles 13 and 14 in *Ceratitis*, *Anopheles*, and *Clogmia*. **k**, Comparison of expression patterns at the head–trunk boundary before gastrulation between species. The expression of *slp* and *eve* are similar across all species. Both *Drosophila* and *Ceratitis*, species with a cephalic furrow, show a head– trunk domain of *btd* that overlaps with *eve* stripe 1. In *Anopheles*, the head–trunk *btd* domain does not overlap with *eve* stripe 1, and in *Clogmia* this *btd* domain is ab_3_s_1_ent. **l**, Schematic drawings showing the combinatorial molecular arrangement at the head–trunk boundary of *Drosophila*. **m**, Profile view of the gene expression at the head–trunk boundary cells in *Ceratitis*, *Anopheles*, and *Clogmia*. ac: *btd* acron domain, sh: *slp* head domain, ht: *btd* head–trunk domain, fg: *btd* foregut domain.

Since *slp1* is a known anterior repressor that positions anterior pair rule stripes,^20,21^ we wondered if the shift in the position of the cephalic furrow in *slp* mutants coincided with a shift in typical overlap between *btd* and *eve* stripe 1 at the head–trunk boundary. We find that the expression patterns and typical overlap between the two genes remains almost unaltered, except for the small ectopic expression *btd* in the head and for the wider gap between *eve* stripe 1 and 2 (Fig. 5c,d, Extended Data Fig. 6b). The anterior shift in cells expressing *btd* and *eve* corresponds to a few rows of blastoderm cells (control = 4.7±0.5 rows, n=4; *slp* = 7.3±0.7 rows, n=5) (Fig. 5e). Despite the displacement, most *slp* embryos exhibit initiator cell behaviors, suggesting that the patterning is not entirely perturbed (Extended Data Fig. 6c). However, the symmetry of the resulting fold is altered (Extended Data Fig. 6d), indicating that *slp1* may contribute not only to the positioning but also to the patterning of individual cells that give rise to the cephalic furrow.

By analyzing the expression of *slp1* relative to *btd*, *eve*, and *prd* in wildtype *Drosophila* embryos, we find that the early *slp1* and *eve* expression domains demarcate the head–trunk boundary from the onset of zygotic activation until gastrulation (Fig. 5f, Extended Data Fig. 6a). Early *slp1* transcripts are limited to the anterior end while *eve* transcripts, which are initially ubiquitous,^21^ begin to clear from the anterior end at cycle 11, and become limited to the posterior region of the body from cycle 12 (Extended Data Fig. 6a). At this stage, the two genes form broad, complementary territories that correspond to the head and trunk regions of the embryo, with the domains juxtaposed at ∼70% of the embryo length (Extended Data Fig. 6a). We first detect *btd* and *prd* transcripts at this interface (Extended Data Fig. 6a,e). During subsequent stages, the *slp1*–*eve* boundary progressively resolves into narrow abutting stripes giving rise to the row of initiator cells (Fig. 5f, Extended Data Fig. 6a). Altogether, the data suggests that *slp1* activity contributes to restricting the anterior boundary of *eve* expression during early stages of zygotic activation, an interaction that determines the site of invagination of the cephalic furrow along the anteroposterior axis.

At the onset of gastrulation, the expression of *btd*, *eve*, *slp*, and *prd* at the head–trunk boundary of *Drosophila*, forms a unique combinatorial code that coincides with the different portions of the cephalic furrow (Fig. 5g, Extended Data Fig. 6f). The central row of *eve*-expressing initiator cells are surrounded by *slp1*-expressing adjacent cells, with *prd* expression offset by a single row of cells relative to the inner *slp1* stripes (Fig. 5g,l, Extended Data Fig. 6f). Moreover, *slp1*-expressing cells also demarcate the outer edges of the invagination (Fig. 5i,l). This molecular arrangement is disrupted in mutants that exhibit cephalic furrow defects (*btd*, *eve*, and *prd*) (Extended Data Fig. 6g,h,i). The combinatorial expression suggests that each row has a unique transcriptional identity and that this specific molecular profile is important for the patterning and morphogenesis of the invagination in *Drosophila*.

### Association of *buttonhead* with cephalic furrow evolution

To uncover the differences in the head–trunk genetic patterning associated with the evolution of the cephalic furrow, we compared the expression patterns in *Drosophila* to three other dipteran species. One species, the Mediterranean fruit fly *Ceratitis capitata* (Tephritidae),^22,23^ belongs to a family known to form a cephalic furrow. The other two species, the malaria mosquito *Anopheles stephensi* (Culicidae)^24,25^ and the drain fly *Clogmia albipunctata* (Psychodidae),^14,26^ belong to families where the cephalic furrow has not been observed.^14^

The three species show early, juxtaposing domains of *slp* and *eve* demarcating the head and trunk regions in a pattern highly similar to that of *Drosophila* (Fig. 5f,j, Extended Data Fig. 7a,b,c). Moreover, the late pattern of abutting *slp* and *eve* stripes is nearly identical between the four species (Fig. 5f,j,k). Only the expression of *prd* in *Clogmia* differs from *Drosophila*, since the prd-expressing cells are not offset from *slp* and *eve* (Extended Data Fig. 7d,e) These observations suggest that the genetic interactions that establish the head–trunk boundary position in *Drosophila*, and much of the late patterning, might be conserved in other dipteran species.

However, we identified a key difference in the expression of *btd* between species with and without the cephalic furrow. While the head–trunk domain of *btd* overlaps with *eve* stripe 1 in *Drosophila* and *Ceratitis*, it does not overlap with *eve* in *Anopheles* and is entirely absent in *Clogmia* (Fig. 5k,l,m). Moreover, the *btd*–*eve* overlap is absent in *Chironomus*, another dipteran that lacks a cephalic furrow.^14^ This comparative expression data suggests the evolution of the cephalic furrow may have been associated with changes in the localization of *btd* along the head–trunk boundary and emergence of a *btd*–*eve* overlap in this region. The establishment of this regulatory framework may have been essential to stabilize the cephalic furrow as a patterned morphogenetic process, and thus enable its mechanical role in absorbing compressive stresses and preventing epithelial instabilities during dipteran gastrulation.

## Discussion

Our work investigates the developmental function and patterning evolution of the cephalic furrow—an epithelial invagination that forms at the head–trunk boundary of dipteran flies. We find that without the cephalic furrow, the head–trunk tissues become unstable during gastrulation and buckle due to compressive stresses exerted by the expansion of mitotic domains and the germ band extension.

Compression is a key mechanism driving the formation of epithelial folds in development,^3,27^ particularly when tissues are under confinement.^17^ Cell divisions can increase epithelial instability due to the in-plane, outward forces generated during the elongation phase^28^ and to the imbalance caused by the basal detachment of dividing cells.^29^ This is the case for the tracheal placode of flies^30^ and intestinal villi of mice,^31^ where mitotic rounding induces the formation of epithelial folds. Interestingly, the folding in these tissues only occur when the epithelium is under compression, similar to our findings that only the combined action of mitotic expansions and germ band extension can induce ectopic folds (Fig. 3j). Moreover, complementary experiments performed in an independent study corroborate the role of mitotic domains and the germ band as sources of mechanical stress.^14^ The consolidated data, therefore, indicates that the head–trunk tissues of flies are under increased compressive stress during gastrulation. We provide evidence that the cephalic furrow counteracts these stresses. An early invagination effectively reduces the appearance of ectopic folds *in vivo* and in simulations, supporting the hypothesis that the cephalic furrow prevents the build up of compressive stresses at the head–trunk boundary, and thus, accomplishes a mechanical role during *Drosophila* gastrulation.

This physical role raises the intriguing idea that the cephalic furrow evolved as a response to mechanical instability. In this case, we expect that increased epithelial instabilities might have a detrimental effect on embryonic development and potentially decrease the fitness of individuals. Mechanical compression can trigger ATP release,^32^ calcium signaling,^33^ and even DNA damage^34^ in mechanosensitive cells. Moreover, variable tissue buckling can disrupt short-range signaling, cell-to-cell interactions, or potentially slow down embryogenesis at the head–trunk boundary. Investigating these effects *in vivo* is challenging, and our efforts to prevent cephalic furrow formation without affecting other developmental processes have failed so far. However, there is evidence that inhibiting the invagination via optogenetics increases the frequency of a distorted ventral midline,^14^ suggesting that mechanical instability can impact the robustness of morphogenetic processes. In this sense, the appearance of a patterned head invagination may have improved developmental robustness and speed and been positively selected during evolution.

The distribution of cephalic-furrow-related traits onto the dipteran phylogeny is consistent with the hypothesis of mechanical instability as a selective pressure for the evolution of morphogenesis (Fig. 6a). Mitotic domains and the germ band extension are ancestral traits, common to Diptera, while the cephalic furrow is a derived trait—an evolutionary novelty of cyclorraphan flies^14^ (Fig. 6a). Since these sources of stress were probably present at the dawn of dipterans, their head–trunk interface may have endured mechanical instabilities long before the evolution of the cephalic furrow. Remarkably, flies without a cephalic furrow (e.g., *Clogmia* and *Chironomus*) exhibit out-of-plane cell divisions at the head–trunk boundary, suggesting that they evolved an alternative solution to mitigate the effect of tissue compression during gastrulation^14^ (Fig. 6a). Moreover, neither *Clogmia*, *Anopheles*, nor *Chironomus* exhibit the *btd*–*eve* expression overlap at the head–trunk boundary^14^ (Fig. 5), a trait essential to specify the initiator cells in species with cephalic furrow like *Drosophila*. Therefore, the establishment of a *btd*–*eve* overlap was probably a key event associated with the origin of the cephalic furrow and that might explain the divergent morphogenetic solutions to a similar selective pressure of mechanical instability (Fig. 6b).

**Figure 6:**
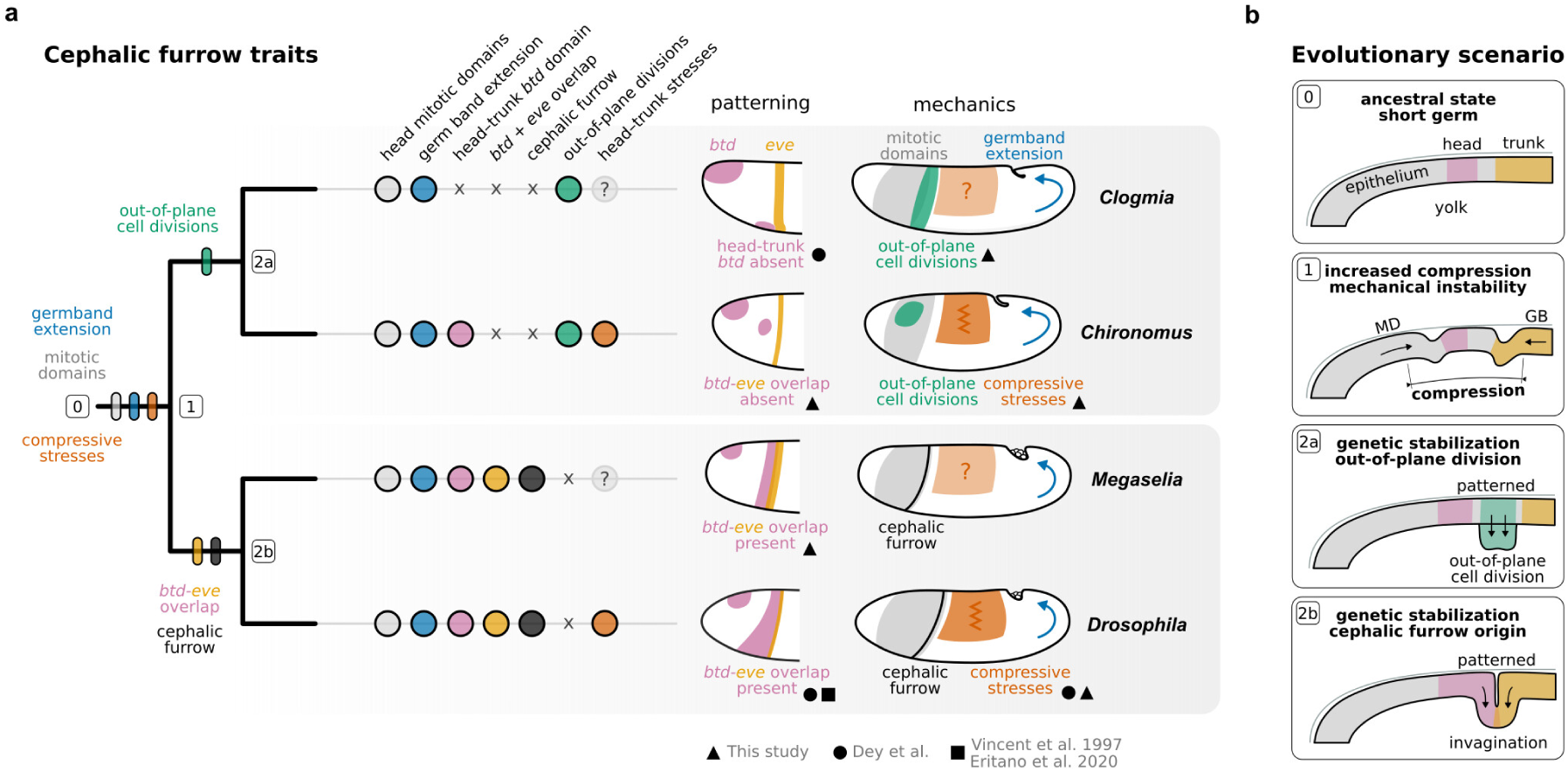
Interplay of genetics and mechanics during cephalic furrow evolution. **a**, Simplified dipteran phylogeny based on Ref^64^ with cephalic furrow traits mapped onto the tree. The germ band extension and mitotic domains are ancestral, suggesting that compressive stresses at the head–trunk boundary were present since the dawn of Diptera. The cephalic furrow is a derived trait, an evolutionary novelty of cyclorraphan flies present in the common ancestor of *Megaselia* and *Drosophila*, and the out-of-plane cell divisions at the head–trunk boundary are present in non-cyclorraphan flies such as *Clogmia* and *Chironomus*.^14^ The presence of a *btd*–*eve* overlap correlates with the presence of the cephalic furrow, and may be associated with its evolution. Data sources for the figure are annotated with geometrical symbols; this study (black circle), Dey *et al.*^14^ (black triangle), and Eritano et al.^9^ and Vincent et al.^8^ (black square). **b**, Scenario for mechanical instability as a selective pressure for the evolution of morphogenetic innovations. In the ancestral state, there was no mechanical instability at the head–trunk boundary [0]. The appearance of mitotic domains and germ band extension increased the compressive stresses and ectopic buckling events at the head–trunk boundary [1]. This mechanical instability may have had a detrimental effect on individual fitness by affecting developmental robustness or by slowing down embryogenesis. Natural selection favored the establishment of patterned processes that mitigate these compressive stresses at the head–trunk boundary. One solution, present in *Clogmia* and *Chironomus*, is the out-of-plane cell divisions, which reduce the compression load on the monolayer epithelium [2a]. Another solution, present in *Drosophila* and other cyclorraphan flies, is the formation of an out-of-plane invagination which absorbs the mechanical forces at the head–trunk boundary [2b]. These events may have happened through the stabilization of genetic interactions and cooption of existing signaling modules controlling cell and tissue morphogenesis. Tissue mechanics may have been an important factor influencing the evolution of patterned morphogenesis in early embryonic development.

Our data reveals how the interplay between genetic patterning and tissue mechanics may have shaped the evolution of morphogenetic processes in the early embryonic development of flies. These findings, however, uncover a potentially more general mechanism about how tissue mechanics can influence developmental evolution. Classical theoretical work by Newman and Müller raised the hypothesis that self-organized morphogenesis was critical to generate morphogenetic innovations at the dawn of animal evolution.^35^ Our work provides initial empirical evidence supporting this hypothesis, showing that mechanical forces might have had a critical role in generating morphogenetic innovations not only at the transition to multicellularity, but also after the establishment of developmental patterning systems across evolutionary time. We propose that the genetic resolution of mechanical conflicts between distinct embryonic processes may be a broadly occurring mechanism that contributes to generating the remarkable morphogenetic diversity of early animal embryogenesis.

## Methods

### *Drosophila* stocks and genetic crossings

To generate fluorescent cephalic furrow mutants, we performed genetic crosses using the loss-of-function alleles *btd^XA^*(FBal0030657), *eve^3^* (FBal0003885), *prd^4^* (FBal0013967), *slp^Δ34B^*(FBal0035631), and *stg^2^* (FBal0247234); the membrane fluorescent marker *Gap43-mCherry* (FBal0258719, gift from Kassiani Skouloudaki); and the green fluorescent balancers *FM7c, Kr-GFP* (FBst0005193), *CyO, twi-GFP* (gift from Akanksha Jain), and *TM3, Kr-GFP* (FBst0005195). We established stable lines balancing the loss-of-function alleles with fluorescent balancers, and used the lack of GFP signal to identify homozygous embryos in our live-imaging recordings. For genes on chromosomes 1 and 2 (*btd*, *eve*, and *prd*), we added the membrane marker on chromosome 3 ([*btd^XA^/FM7c, Kr-GFP;; Gap43-mCherry/MKRS*], [*eve^3^/CyO, twi-GFP; Gap43-mCherry/MKRS*], and [*slp^Δ34B^/CyO, twi-GFP; Gap43-mCherry/TM6B*]). For *stg*, which is located on chromosome 3, we recombined the allele with Gap (*Gap43-mCherry, stg^2^/TM3, Kr-GFP*). Since the *btd*–*stg* double mutant stable line is weak, we imaged the progeny of *btd^XA^/FM7c, Kr-GFP;; Gap43-mCherry, stg^2^/Gap43-mCherry* flies, identifying *btd* homozygozity by the GFP, and *stg* homozygozity by the lack of cell divisions after gastrulation. For laser ablations, we used a *moe-GFP* line (gift from Elizabeth Knust). The wildtype stocks contain the Gap43-mCherry marker in the Oregon-R genetic background. We obtained the founder fly stocks from the Bloomington Drosophila Stock Center and the Kyoto Stock Center and deposited the lines in the MPI-CBG stock collection. The complete list of FlyBase^36^ accession numbers and genotypes is available in the project’s data repository.^37^

### Animal husbandry and embryo collection

We maintained the *Drosophila* stocks in 50 mL hard plastic vials containing standard fly food and enclosed with a foam lid to allow air exchange. They were kept in an incubator with a constant 25° C temperature and 65% humidity and a 12:12 h light cycle. For imaging, we first amplified the stocks in larger 200 mL vials for a few weeks. We then narcotized the flies with CO_2_, and transferred them to a cage with a plate attached to one end containing a layer of apple juice agar and a slab of yeast paste on top. The flies were left to acclimatize in the cage for two days before the experiments. To guarantee the embryos are at a similar developmental stage, we exchanged the agar plate once per hour at least twice (pre-lays), and let the flies lay the eggs on the agar for one hour before collecting the plate. After filling the plate with water, we used a brush to release the eggs from the agar and transferred them to a cell strainer with 100 µm nylon mesh (VWR). To remove the chorion, we immersed the embryos in 20% bleach (sodium hypochlorite solution, Merck 1.05614.2500) for 90 s, washed abundantly with water, and proceeded to mounting for live imaging.

We maintained *Clogmia* flies in 9cm-wide plastic Petri dishes with a layer of wet cotton at room temperature and fed weekly with powder parsley. To obtain embryos for fixation, we collected the adult flies in a 200 mL hard plastic vial with wet cotton, and let them mate for 2–3 days. Then, we anesthetized the flies with CO_2_, dissected the ovaries from ripe females, and released the eggs using tweezers in deionized water, which activates embryonic development.^38,39^ We let embryos develop in deionized water at room temperature until the desired stage. To remove the chorion, we transferred the embryos to a glass vial with 0.5x PBS using a fine brush, exchanged the medium for 5% bleach in 0.5x PBS for 2 min, and washed abundantly with 0.5x PBS. Using the diluted PBS solution instead of water prevents the embryos from bursting after dechorionation.

We obtained pupae of the EgyptII wild type strain of *Ceratitis* from the Insect Pest Control Laboratory of the International Atomic Energy Agency (IAEA). Adult flies were kept at 25° C, 65% humidity, and 12:12 h light cycle, inside 49×30×30 cm plexiglass cages with the front and back ends covered by a nylon mesh. We provided water through a soaked towel and food as a 3:1 sugar:yeast mixture. As ripe females lay the eggs through the nylon mesh, we placed a plastic container with water at the back end of the cage for several hours to collect eggs. We dechorionated *Ceratitis* embryos using the *Drosophila* protocol.

We performed the collection of *Anopheles* embryos at the Center for Integrative Infectious Diseases Research (CIID) at Heidelberg University. To collect embryos, we placed a glass container with water and filter paper inside a cage with 300 mated females, which were fed a blood meal 72 h before, and put the cage in the dark for 2 h at 29° C. We then removed the container from the cage and let the embryos develop until the desired stage. To dechorionate, we collected the embryos on a cell strainer, incubated in 5% bleach for 75 s, and washed them thoroughly with deionized water.

### Embryo fixation and *in situ* hybridization

For *Drosophila* and *Ceratitis*, we transferred dechorionated embryos to a glass vial containing equal volumes of 4% formaldehyde in PBS and n-Heptane, and let the vial shaking at 215 rpm for 45 min. For *Clogmia*, we diluted the fixative in 0.5x PBS. After removing the fixative (lower phase) using a glass pipette, we added an equal volume of 100% methanol, and shook the vial vigorously by hand for 1 min. We then removed the n-Heptane (upper phase) and collected the embryos on the bottom of an Eppendorf tube and washed several times with 100% methanol. For *Anopheles*, we followed a similar protocol that includes a longer 30 min wash in water after fixation, a 30 s boiling water step followed by 15 min in ice-cold water, until the final methanol washes.^24^ All the samples were stored in 100% methanol at −20° C.

We performed the *in situ* hybridization of *btd*, *eve*, *prd*, and *slp* genes using the Hybridization Chain Reaction v3.0 (HCR^TM^)^40^ reagents, except for the probe sets, which we designed using a custom script. The oligos were obtained from Sigma-Aldrich. We selected the HCR^TM^ amplifiers to allow for triple (multiplexed) *in situ* combinations of *btd*+*eve*+*slp* or *prd*+*eve*+*slp*. Before starting, we manually devitenillized *Anopheles* embryos using fine tweezers. Then, we rehydrated the embryos in 100% methanol with a series of washes to 100% PBT. We permeabilized the embryos with 1:5000 dilution of ProteinaseK (20 mg/mL) for 5min, except for *Drosophila*. All samples were re-fixed in 4% formaldehyde for 40 min and washed thoroughly with PBT. We then followed the *“In situ HCR v3.0 protocol for whole-mount fruit fly embryos Revision 4 (2019-02-21)”* from Molecular Instruments molecularinstruments.com/hcrrnafish-protocols. After the protocol, we stained the embryos with 1:1000 DAPI in 5x SSCT solution for 2 h and mounted the embryos in 80% glycerol in 5x SSCT for imaging.

### Sample mounting for microscopy

For most of our live imaging, we used a Zeiss Lightsheet Z.1 microscope. To increase the throughput of samples imaged in one session, we optimized a mounting strategy developed previously in our laboratory.^41^ First, we cut a 22×22 mm glass coverslip (0.17 mm thickness) into 6×15 mm strips using a diamond knife, and attached a single strip to a custom sample holder using silicon glue, letting it harden for 15 min. We then coated the coverslip strip with a thin layer of heptane glue and let it dry while preparing the embryos. Using a fine brush, we transferred the embryos collected in the cell strainer onto an agar pad, and oriented them manually with a blunt cactus spine under a stereomicroscope. We aligned about 20 embryos in a single line (head to tail) along the main axis of the strip with the left or ventral sides up, depending on the experiment. To attach the embryos to the coverslip, we carefully lowered the sample holder over the agar pad until the glass coated with heptane glue touched the embryos. We placed the sample holder into the microscope chamber filled with water, and rotated it so that the samples are facing the detection objective directly, and the coverslip is orthogonal to the detection objective; this is important to prevent the lightsheet from hitting the glass edges. With the embryos oriented vertically along the coverslip, the lightsheet generated from the illumination objectives coming from the sides only needs to pass through the width of the embryo (about 200 µm). This approach gives the best results for recording lateral and dorsal views and is ideal for live imaging homozygote embryos since they are only about one fourth of the total number of imaged embryos. For fixed imaging of *in situ* samples, we used an inverted Zeiss LSM 700 Confocal microscope. We mounted the samples immersed in 80% glycerol between a slide and a glass coverslip supported by tape.

### Microscopy acquisition parameters

For the lightsheet lateral datasets, we used a Zeiss 20x/1NA Plan-Apochromat water immersion objective to acquire stacks with 0.28 µm XY-resolution and 3 µm Z-resolution covering half of the embryo’s volume in a single view. This Z-resolution was restored to 1 µm during image processing (see below). For the dorsal datasets, we used a Zeiss 40x/1NA Plan-Apochromat water immersion objective to acquire stacks with 0.14 µm XY-resolution and 3 µm Z-resolution covering a volume around in the middle section of the anterior end of the embryo. We adjusted the time resolution between 45–60 s per frame to maximize the number of embryos acquired in one session. To visualize both the membrane signal (mCherry) and the green balancer signal (GFP), we acquired two channels simultaneously using the 488 and 561 nm lasers at 3% power with an image splitter cube containing a LP560 dichromatic mirror with SP550 and LP585 emission filters. All live imaging recordings were performed at 25° C. For the confocal datasets, we used a 20x/0.8 Plan-Apochromat Zeiss air objective to acquire 4-channels using 3 tracks (405, 488 and 639, and 555 nm, respectively) with a BP575-640 emission filter and about 0.4 µm XY-resolution and 2 µm Z-resolution covering about half the embryo’s volume.

### Image processing and visualization

We converted the raw imaging datasets into individual TIFF stacks for downstream processing using a custom ImageJ macro in Fiji.^42,43^ To visualize the presence and dynamics of ectopic folds, we generated 3D renderings of the surface of embryos in lateral recordings using the plugin 3Dscript in Fiji.^44^ For analyzing the entire epithelial surface, we first improved the signal-to-noise ratio and Z-resolution of lateral datasets from 3 µm to 1 µm by training a deep learning upsampling model using CARE.^45^ Then, we created cartographic projections of the lateral recordings using the ImSAnE toolbox^46^ by loading the restored data in MATLAB,^47^ segmenting the epithelial surface using ilastik,^48^ and generating 3D cartographic projections of lateral views following a workflow established for fly embryos.^49^ To visualize *in situ* hybridization data, we performed maximum intensity projections or extracted single slices from the raw volumes. For all microscopy images, we only performed minimal linear intensity adjustments to improve their contrast and brightness.^50^

### Ectopic fold analyses

To characterize the relative timing of ectopic folding, we annotated the germ band position and the number of frames after the onset of gastrulation at the initial buckling, when the first cells disappear from the surface in the lateral 3D renderings. We defined the onset of gastrulation (T=0) as the moment immediately after the end of cellularization, and immediately before the beginning of the ventral furrow invagination. To visualize the variability of ectopic folding, we manually traced the fold outlines in lateral recordings using Fiji. Because embryos have different sizes, we first used the plugin *bUnwarpJ^51^* (imagej.net/plugins/bunwarpj) to register individual frames and then applied the same transformation to the fold traces for a standardized comparison. We analyzed the dynamics of ectopic folds by measuring the relative angle and tortuosity of the segmented line traces over time, and to visualize the kinetics we generated color-coded temporal projections using the script *Temporal Color Code* (imagej.net/plugins/temporal-color-code) with the perceptually uniform *mpl-viridis* color map (bids.github.io/colormap) bundled in Fiji.

To estimate the folded area in the cephalic furrow and ectopic folds, we annotated the region of the blastoderm before gastrulation that infolded in the cartographic projections using Fiji, and calculated the area correcting the pixel dimensions according to the coordinates in the projection. For the fold depth, we measured the distance between the vitelline envelope to the tip of the fold at the moment of maximum depth in the dorsal recordings. For the analysis of the epithelial surface, we used the plugin *MorphoLibJ^52^* (imagej.net/plugins/morpholibj) to segment, measure, and color-code the cell apical areas, and the plugin *Linear Stack Alignment with SIFT* (imagej.net/plugins/linear-stack-alignment-with-sift) to register cells between timepoints.

### Laser cauterization experiments

We performed laser cauterization experiments in two microscope setups, a Luxendo MuVi SPIM with a photomanipulation module and a Zeiss LSM 780 NLO with multiphoton excitation. For the MuVi SPIM, we embedded dechorionated embryos in 2% low-melting agarose and mounted the samples in glass capillaries to obtain *in toto* recordings. We used a pulsed infrared laser 1030–1040 nm with 200 fs pulse duration and 1.5W power to cauterize the posterior region of the dorsal embryonic surface, attaching the blastoderm to the vitelline envelope. Using an Olympus 20x/1.0NA water immersion objective, we acquired stacks with 0.29 µm XY-resolution and 1 µm Z-resolution of four different angles every one minute. For the Zeiss microscope, we attached the embryos with the dorsal side down onto coverslips using heptane glue and immersing in halocarbon oil. We cauterized the embryos sequentially using a near infrared 800 nm laser (Chameleon Vision II) through a single pixel line (210 nm/px and 100 µs/px) around the same dorsal region to block the germ band extension. We used a Zeiss 25x/0.8NA LD LCI Plan-Apochromat glycerol immersion objective to acquire every 2:38 min two different planes of the blastoderm, (i) the surface to monitor the germ band extension, and (ii) 40 µm deep in the equatorial region to monitor the occurrence of ectopic folding. The stacks had 0.21 µm XY-resolution and one minute time resolution. To obtain a quantitative measure of ectopic folding, we analyzed the degree by which the tissues deform between non-cauterized and cauterized mutants, using as a proxy the tortuosity of the epithelium outline. For that, we took the profile slices from dorsal recordings and transformed the curved vitelline envelope into a straight line using the *Straighten* tool of ImageJ (Supplementary Fig. 2a). We then cropped a 200×25 µm region along the head–trunk interface and applied Gaussian blur, thresholding, and edge detection to obtain the epithelium outline for individual timepoints covering about 50 min after gastrulation (Supplementary Fig. 2a,b). We extracted measurements from the epithelium outlines using the ImageJ plugin *Analyze Skeleton^53^* (imagej.net/plugins/analyze-skeleton), and generated the color-coded temporal projections as described above. Laser ablation experiments

We performed laser ablations in a Yokogawa CSU-X1 confocal spinning disk, an EMCCD camera (Andor iXon DU-888), and the software AndorIQ for image acquisition. We attached dechorionated embryos laterally to a MatTek glass-bottom petri dish and covered the samples with water, and performed the ablations using a Titanium Sapphire Chameleon Ultra II (Coherent) laser at 800 nm tuned down from 80 MHz to 20 kHz with a pulse-picker. The laser power measured before the microscope port was 6 mW and the pixel dwell time for scanning was 2 µs. To ensure the cut, we repeated the scan ten consecutive times along a single cell, acquiring a single slice with a 60x/1.2NA water immersion objective with 0.18 µm XY-resolution and 200 ms time-steps. We ablated each embryo just once. The temperature was maintained at 28° C. To analyze the ablation data, we created a line crossing the edges of the ablated cell perpendicular to the cut and generated a kymograph using the *Multi Kymograph* Fiji plugin (Supplementary Fig. 3). We then binarized the kymographs, measured the distance between cell edges over the first 30 s after the cut, and performed a linear fit of the data to obtain the recoil velocity (Supplementary Fig. 3).

### Strain rate analysis

To estimate the strain rates, we first performed particle image velocimetry on cartographic projections using the ImageJ plugin *iterativePIV* ^54^ (sites.google.com/site/qingzongtseng/piv). Then, we used the equation

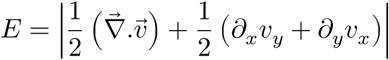

to define and calculate the magnitude of the strain rate, where is the displacement obtained in the PIV divided by the time in minutes. The measurements combine isotropic and anisotropic strain rate. We used these values to create a color-coded overlay for the strain rate (Extended Data Fig. 3b). To generate the line plots, we averaged the strain rate along the dorsoventral axis in two predefined regions, the head–trunk (canonical cephalic furrow position) and the trunk–germ (posterior to the Mitotic Domain 6) (Extended Data Fig. 3b).

### Model and simulations

Our model follows an approach similar to a previously published model of epithelial buckling under confinement.^17^ It represents the monolayer epithelium of the early *Drosophila* embryo in a cross-section as a single line through the apical–basal midline of the epithelial cells. The tissue is modeled as an elastic rod with a stretching energy per unit length *W*_*s*_ and bending energy per unit length *W*_*b*_ so that the total energy per unit length is *W*_*T*_ = *W*_*s*_ + *W*_*b*_. In full,

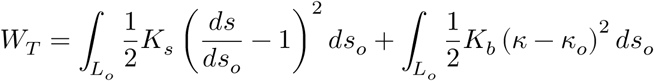

where *K_s_* is the stretching rigidity and *K_b_* is the bending rigidity of the tissue; *ds_o_* and *d_s_* are the preferred and current lengths of the curve, respectively; and is the curvature of the rod. *L_o_* is the total length of the tissue in a stress-free condition. To perform numerics, we discretize the curve into *N* particles indexed by *i*. The total energy per unit length for this discretized model is given by

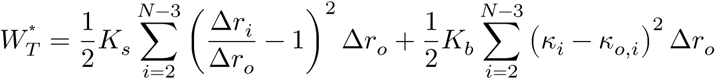

where Δ*r*_*o*_ is the preferred length of springs connecting consecutive points (equal for all springs); Δ*r_i_* is the current length between *i* and *i* + 1;*k_i_*; is the discretized curvature at point ; *i*; *k_o,i_* is the preferred curvature at point *i* (equal to 0, except when specified). The first and last two points of the curve are fixed in space. To obtain a physically meaningful dimensionless bending rigidity, we divide the bending rigidity by the factor *K*_*s*_*L^2^* as

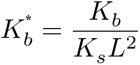

where *L* is the semi-major axis of the embryo. To minimize the total energy, we add a ground level of noise to the particles and let the particles move in the direction of the forces. The motion of the particles is governed by

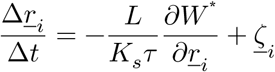

where 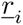 is the current position of the *i*th particle; *τ* represents an arbitrary timescale introduced here to balance dimensions (set to 1); Δ*t* are the timesteps (set to 10^−5^ × *τK_s_/L*); and 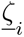 is the noise, chosen from a Gaussian distribution with mean 0 and standard distribution 10^−5^ × *L*. In our model, the position of the germ band corresponds to the position of the last particle in the curve on the semi-ellipse that represents the embryonic blastoderm. The extent of the germ band is given by *g*, which is the projection of the germ band arclength onto the mid-axis of the embryo normalized by the embryo length (2*L*). When *g*= 0 the tissue is free of stretching stress, but at any other 0 < *g* <1, the blastoderm will be compressed. The preferred lengths of the individual springs is obtained by dividing the elliptical arclength into equal segments. The length of each segment is given by 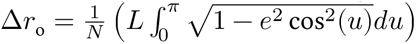. To find the initial lengths of the springs, we use

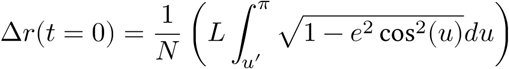

where 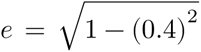 and the angle *u*′ corresponds to the position of the blastoderm end. u′ is obtained for a given value of by *u*′ = cos^−1^(1−2*g)*. Here, we obtained the initial lengths by dividing the compressed blastoderm into equal segments. For any simulation, the value of *g* is constant (blastoderm end is static in position). To model mitotic domains, we introduced new particles and springs on the mid-points between two particles in specific regions of length 0.5*L*. The new springs are given the same Δ*r*_*o*_ as the rest of the springs in the tissue. The blastoderm is confined by a rigid boundary in the shape of a semi-ellipse. Any particle that lands outside this boundary at any timestep is re-positioned onto the rigid boundary. This new position is prescribed by taking the intersection point of the rigid boundary curve and the line segment that connects the position before this iteration (which was inside or on the vitelline envelope) and the position outside the vitelline envelope. Finally, we define and count a *fold* when we find that a particle’s distance from the rigid boundary is greater than a threshold value. To calculate this threshold, we measure the maximum distance that particles can achieve when the tissue is in a stress-free state. This threshold was calculated to be 0.035*L*. The code for the model and simulations is available in a Zenodo repository.^55^

### Data visualization and figure assembly

We created illustrations and assembled the final figure plates using Inkscape v1.2.2.^56^ For microscopy videos, we exported the original stacks as AVI without compression with 10–15 fps using Fiji and post-processed them to MPEG-4 format 1080p resolution using the H.264 encoding at a constant bitrate quality factor of 18 for visualization using HandBrake v1.6.1.^57^ The high-resolution figures and videos are available in a Zenodo repository.^58^ We performed the data wrangling, statistical analyses, and plotting in R v4.2.1^59^ using RStudio v2022.7.2.576,^60^ and Python 3.10.7 using Jupyter notebooks.^61^ All the data and analyses pipelines were deposited in a Zenodo repository.^37^

## Data availability

- Main repository with data, analyses, figures, and text source files: zenodo.org/record/7781947.^37^
- Theory repository with model code and simulation outputs: zenodo.org/record/7784906.^55^
- Media repository with high-resolution figures and videos: zenodo.org/record/7781916.^58^

## Acknowledgments

We thank all current and former LoPaTs for the discussions and support during this project. Akanksha Jain and Vladimir Ulman for initial help with cartographic projections. Giulia Serafini for the help with fly crosses. Anaïs Bailles for constructive feedback. Michaela Burkon and Pavel Mejstrik for technical support. Jan Brugués and Keisuke Ishihara for the laser ablation setup. MPI-CBG’s Light Microscopy Facility and Computer Services Facility for the ample assistance with data acquisition and processing. Sven Ssykor and Cornelia Maas for the support with fly stocks. Radim Žídek for essential help in construct design. Steffen Lemke and Yu-Chiun Wang for cephalic furrow discussions and sharing unpublished data. The Lemke Lab for the generous help in setting up the *Clogmia* cultures. Freddy Frischknecht and Miriam Reinig for enabling and supporting the collection of *Anopheles* embryos. Konstantinos Bourtzis for kindly providing us with *Ceratitis* pupae. Carles Blanch-Mercader for feed-back on simulations. Juliana Roscito for the text revisions. Michele Marass for crucial editorial input. Michael Akam for drawing BCV’s attention to the cephalic furrow.

## Author contributions

BCV and PT conceived the study. BCV designed experiments, generated fly stocks, acquired microscopy data, performed *in situ* hybridization, and processed and analyzed the *in vivo* data. MBC conceived and conducted the laser ablation and cauterization experiments, and analyzed the laser ablation and strain rate data. CDM, AK, and AS designed the model. AK and AS programmed the model, performed the simulations, and analyzed the *in silico* data. BCV wrote the manuscript. All authors revised and contributed to the text.

## Extended data

**Figure Extended Data Fig. 1:**
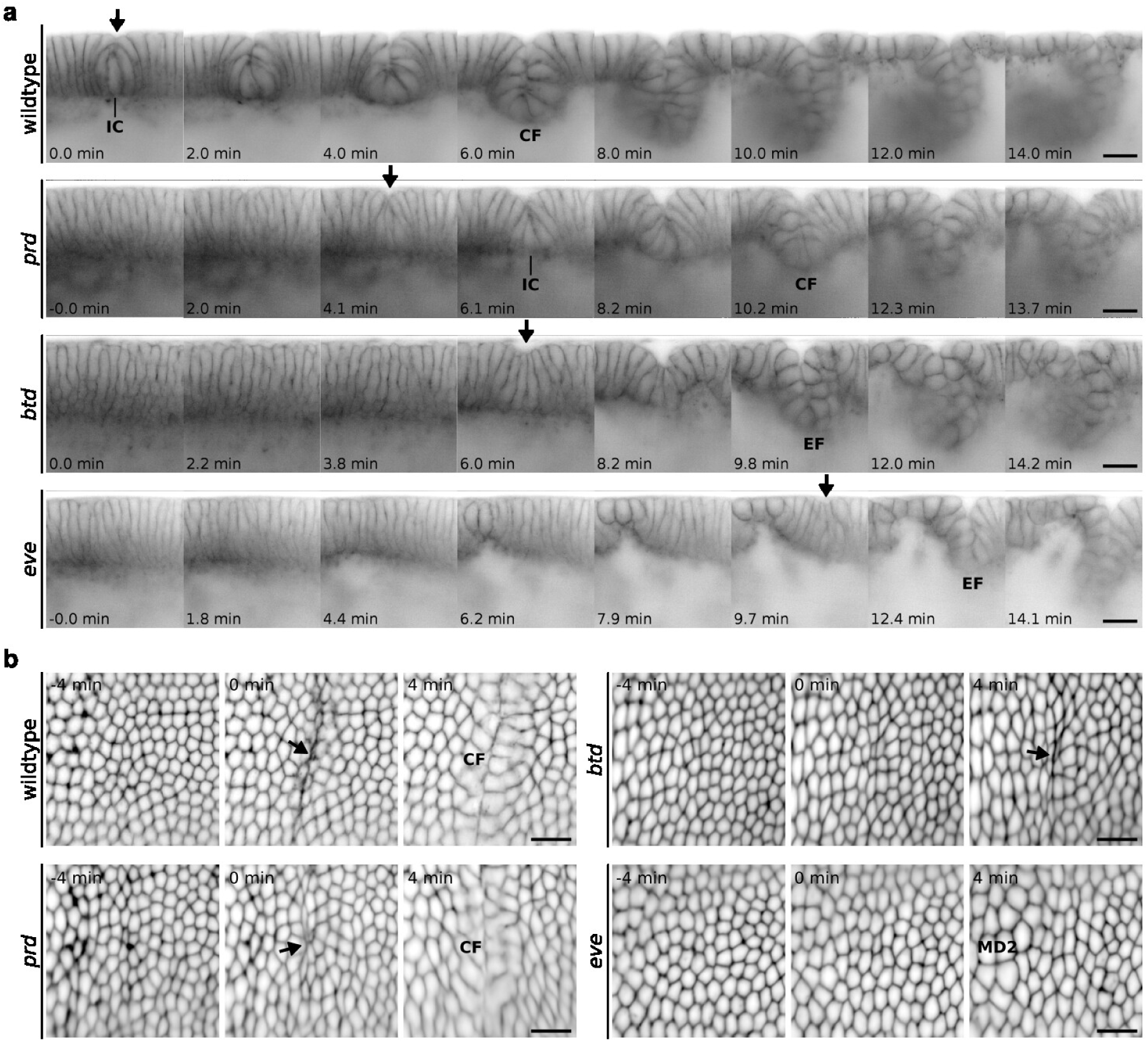
Perturbation of the initiator cell behavior in cephalic furrow mutants. **a**, Profile view showing the cephalic furrow formation in wildtype, *prd*, *btd*, and *eve* embryos. The samples are synchronized by the end of cellularization, when the cleavage furrows reach their basal position (frame 0.0 min). The arrow indicates the position and timing of the first infolding of the tissue. In wildtype, the shortening of initiator cells begins before cellularization is complete (wildtype 0.0 min). Cells adjacent to the initiators remain in close contact to the initiator row. Their apical membranes are tightly connected and become arched in an arrangement typical to the early phase of the invagination of the cephalic furrow (CF). This arrangement is perturbed in mutant embryos to different degrees. In *prd*, the initiator cells shorten and the tissue invaginates, but the infolding is delayed, and the adjacent cells do not arch over the initiator row (*prd* 6.1 min). In *btd*, there is no cell shortening, but some embryos exhibit a certain degree of anisotropic apical constriction which creates a bulge in the epithelium minutes after the end of cellularization (*btd* 6.0 min, see also **b**). This initial bulge often primes the position for the formation of ectopic folds (EF) of ectopic buckling. In *eve*, the cells show neither shortening nor apical constriction and ectopic folds appear about ten minutes after the end of cellularization (*eve* 9.7 min). Scale bars = 20 µm. **b**, Surface view of cartographic projections showing the head–trunk interface. In wildtype, the anisotropic apical constriction is localized to a narrow stripe adjacent to the initiator row. In *prd* embryos, the apical constriction occurs, but it does not form a clear line of infolding cells preceding the invagination as in wildtype embryos. In *btd* embryos, there is a similar degree of anisotropic apical constriction occurring but not all embryos form ectopic folds at this region. In *eve* embryos, the mitotic domain 2 (MD) begins expanding and there is no apical constriction behavior. Time between frames is about 4 min. Scale bars = 20 µm (approximate value).

**Figure Extended Data Fig. 2:**
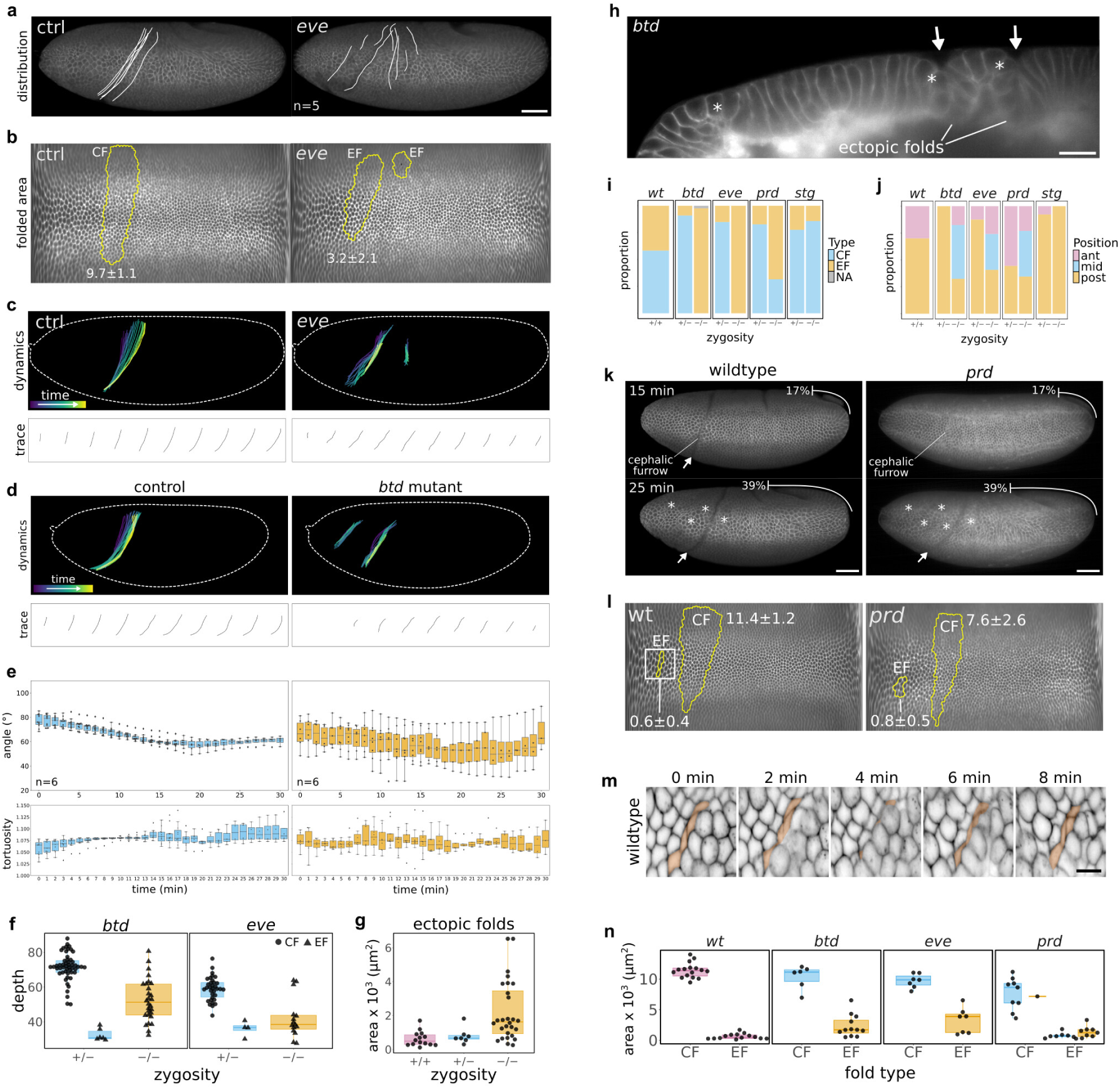
Differences between ectopic folding and cephalic furrow formation. **a**, Distribution of ectopic folds in *eve* homozygotes (right) and sibling controls (left). Scale bar = 50 µm. **b**, Folded area of the cephalic furrow (CF) and ectopic folds (EF) in *eve* embryos. The folded region is outlined in yellow on cartographic projections of a representative sibling control *eve* heterozygote (left) and of a *eve* homozygote (right). The numbers indicate the average folded area for the sample type in µm^2^×10^3^. **c**, Dynamics of cephalic furrow and ectopic fold formation in *eve* mutants. **d**, Dynamics of cephalic furrow and ectopic fold formation in *btd* mutants. **e**, Angle directionality, and tortuosity measurements comparing the cephalic furrow and ectopic fold formation. Within the first fifteen minutes after gastrulation, the cephalic furrow exhibits a typical posterior shift on the dorsal side which declines the initial angle of the invagination from 80° to about 60° in relation to the anteroposterior axis. During this period, the invagination begins as a straight line and bends showing a correspondent increase in the measured tortuosity values of the furrow outline. In contrast, ectopic folds show no obvious trend in angular direction tortuosity values over time. For both angle and tortuosity analysis, n=6. **f**, Maximum depth of the cephalic furrow and ectopic folds in *btd* and *eve* mutants. Ectopic folds are shallower than the cephalic furrow in both genetic backgrounds (*btd* p=6.0e-09 and *eve* p=1.1×10-4 in a Mann-Whitney test). Each point corresponds to a single fold; each embryo can have multiple folds. The number of embryos analyzed for *btd* is 22 heterozygotes and 6 homozygotes, and for *eve* is 14 heterozygotes and 4 homozygotes. **g**, Folded area of ectopic folds in wildtype and mutant embryos (*btd*, *eve*, and *prd*). Ectopic folds in wildtype occupy a smaller area than ectopic folds in cephalic furrow mutants (p=1.6×10e-4 in a Mann-Whitney test). **h**, Profile view of a *btd* mutant embryo, showing the presence of two ectopic folds (arrows) forming next to dividing cells (asterisks). Scale bar = 20 µm. **i**, Proportion of folding types in different genetic backgrounds. **j**, Proportion of folding positions in different genetic backgrounds. **k**, Lateral views of a wildtype (left) and a *prd* mutant (right) exhibiting ectopic folds. Scale bar = 50 µm. **l**, Folded area of the cephalic furrow (CF) and ectopic folds (EF) in the wildtype (left) and *prd* mutant (right) shown in **k**. The folded region is outlined in yellow on a cartographic projection. The numbers indicate the average folded area in µm^2^×10^3^ for the CFs and EFs separately. **m**, Developmental sequence of the wil_4_d_1_type embryo ectopic fold annotated in **l**. Four cells are temporarily infolded during the mitotic expansion of adjacent cells. Scale bar = 10 µm. **n**, Comparison of the folded area between the cephalic furrow and the ectopic folds in different genetic backgrounds.

**Figure Extended Data Fig. 3:**
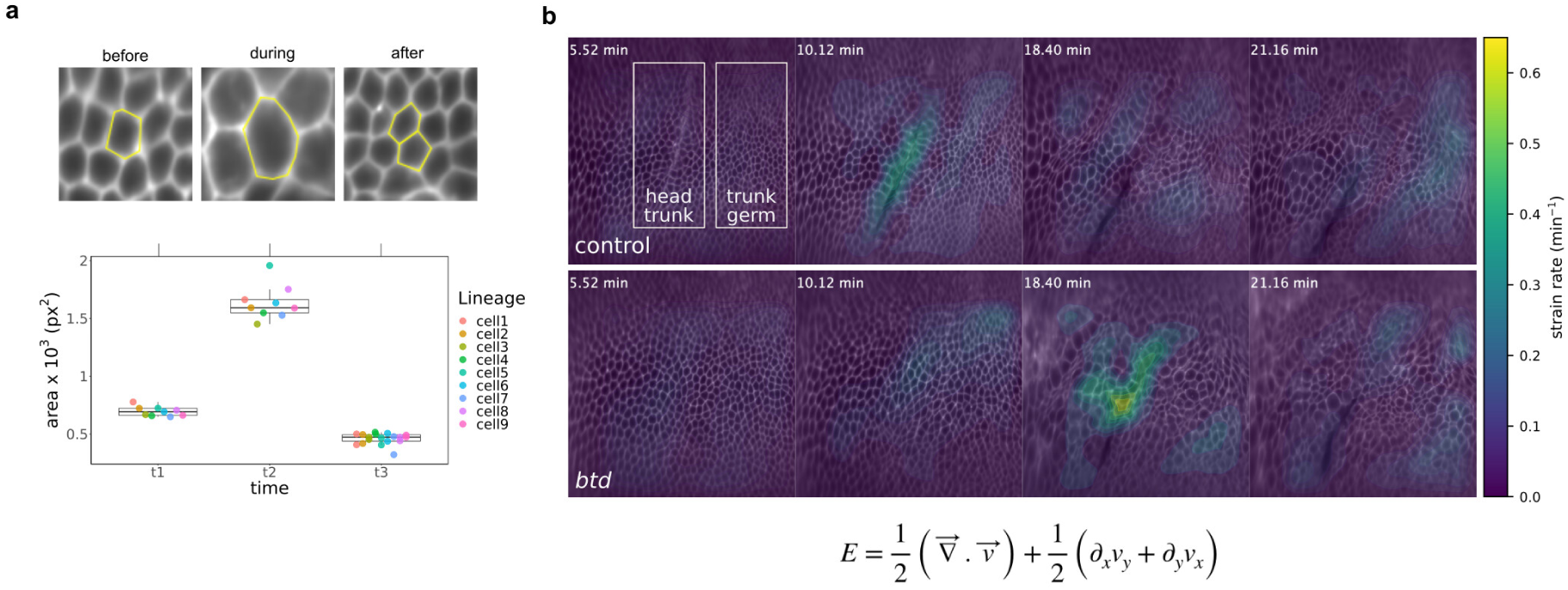
Complementary analyses of mitotic expansions and strain rate. **a**, Increase in the apical area of individual cells within mitotic domains. A dividing cell increases its apical area 2.4 times during mitotic rounding. The individual daughter cells retain 66% of the parent apical area. When summed, the apical area of the two daughter cells occupy 1.3 times the original apical area of their parent cell. **b**, Strain rate analysis in *btd* mutants. Cropped region of cartographic projections of *btd* sibling controls (top, n=3) and homozygote embryos (bottom, n=3). The membrane marker (Gap43-mCherry) is overlayed with a heatmap indicating the regions of increased strain rate in the tissue. The value is the sum of isotropic and anisotropic strain rates obtained through a particle image velocimetry analysis. We used the strain rates in the regions outlined as head–trunk and trunk–germ to generate the plot in Fig. 2f.

**Figure Extended Data Fig. 4:**
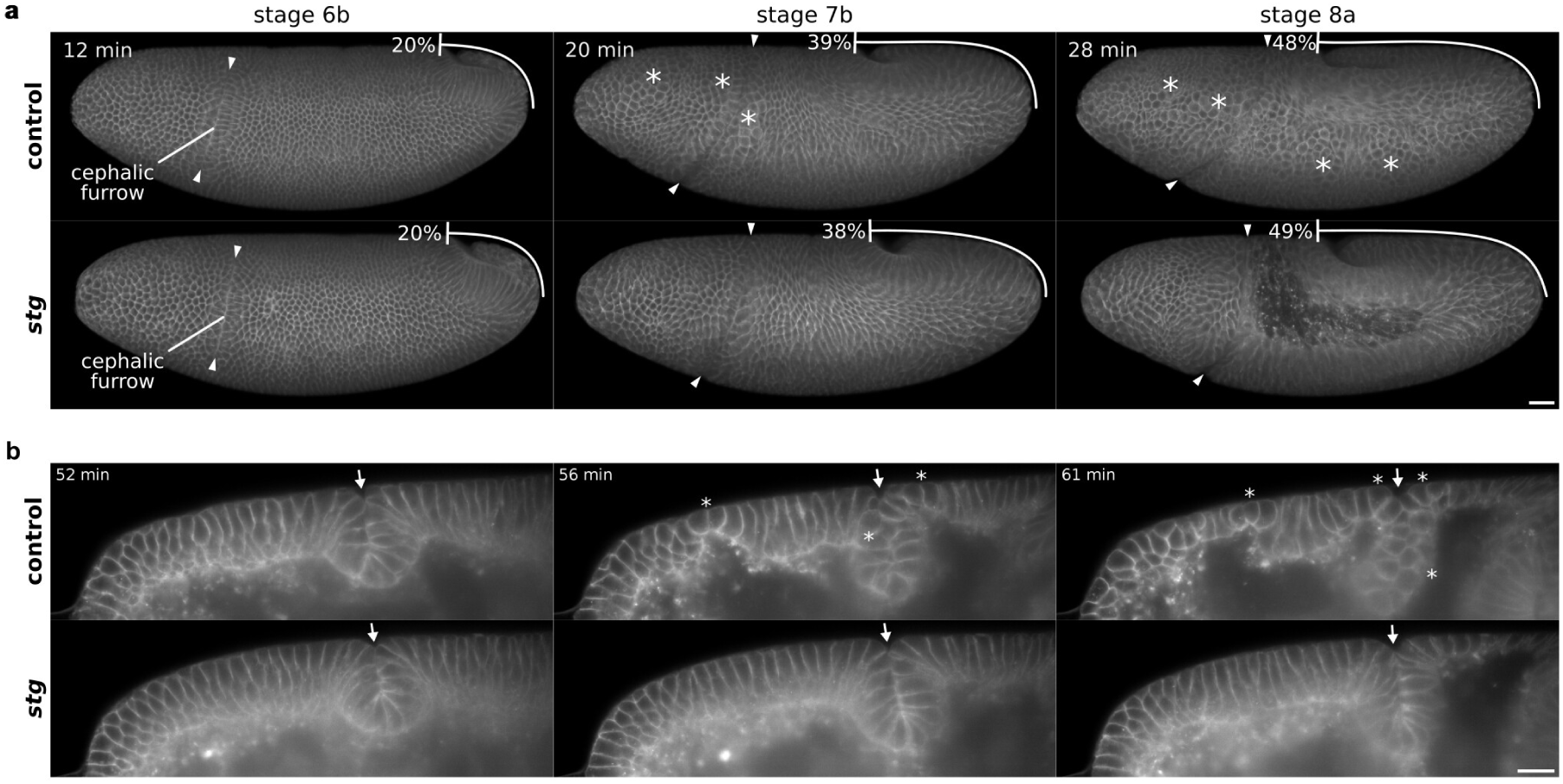
Developmental sequence of *stg* mutants. **a**, Lateral 3D renderings of *stg* homozygotes. They show no cell divisions after gastrulation, but the early morphogenetic movements of gastrulation occur normally. Asterisks indicate mitotic domains in controls. The cephalic furrow forms without delays and exhibits a similar dynamics of invagination compared to wildtype embryos. The only noticeable difference is that the dorsal portion does not shift as posteriorly as in sibling controls, which could be due to the absence of mitotic domains in the head. Scale bar = 50 µm. **b**, Optical slice of profile views in *stg* homozygotes. The initiator cell behaviors are not perturbed and the morphology of the invagination is intact. In fact, because of the lack of cell divisions, the epithelium remains more uniform during gastrulation when compared to sibling controls or wildtype embryos. Scale bar = 20 µm.

**Figure Extended Data Fig. 5:**
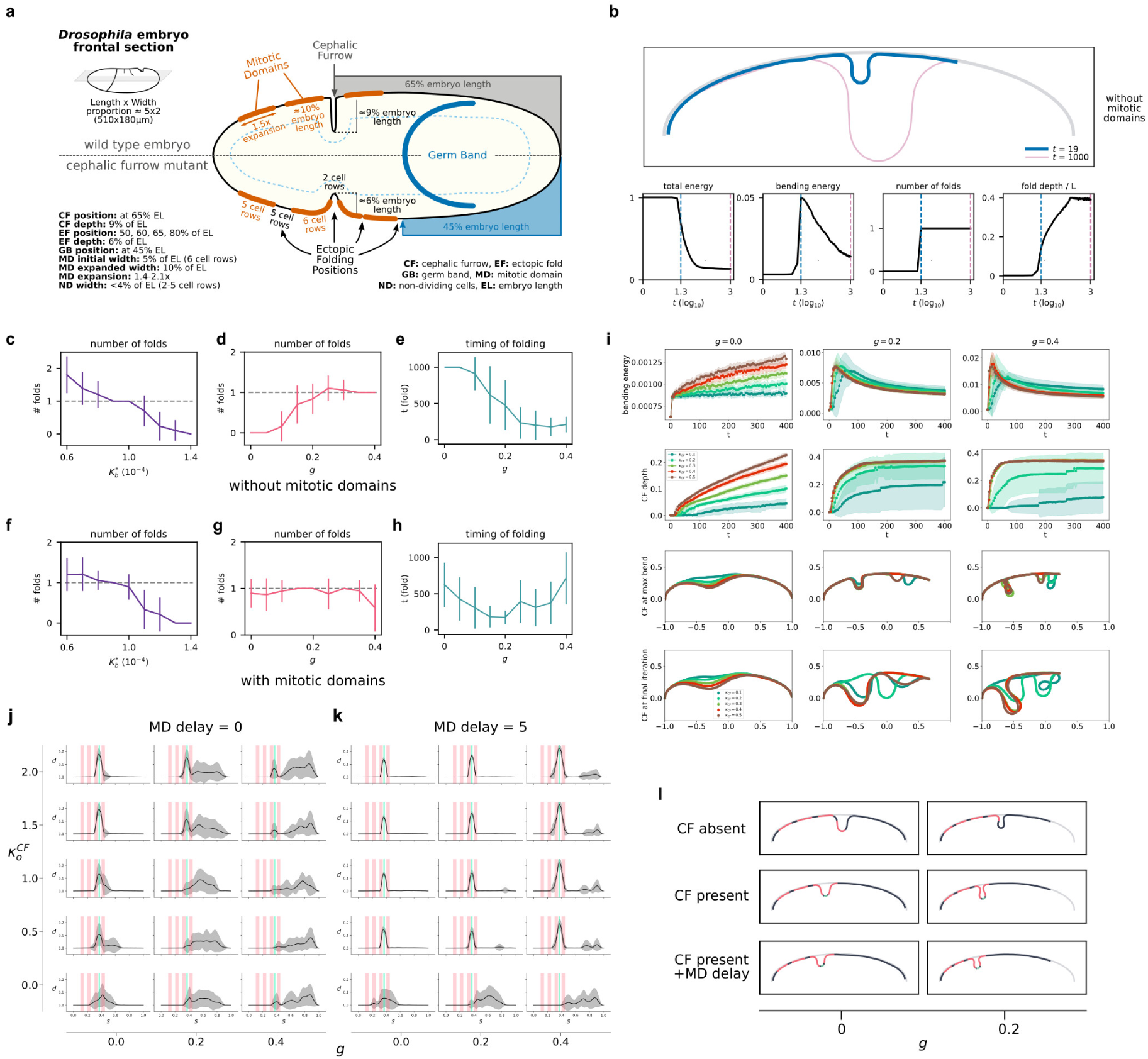
Model general properties and parameter sweeps. **a**, Embryonic proportions in wildtype and cephalic furrow mutants. Approximate relative sizes and positions between embryonic features such as mitotic domains, folds, and the germ band. All values are relative to the embryo length. We used these dimensions as a reference for creating the model. **b**, Representative simulation using 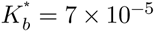 and *g* = 0.3 showing the shape of the tissue at *t* = 19 (blue) and *t* = 1000 (pink). The respective timepoints are marked in dashed lines in the descriptive plots below. They show the variation in total energy, bending energy, number of folds, and fold depth over the iterations. *t* = 1 corresponds to 10^5^ computational steps and the X axis is in *log*_10_ scale to improve the visualization. **c–e**, Parameter sweep without mitotic domains. Plots show (**c**) the number of folds by bending rigidity 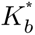 using *g* = 0.3, (**d**) the number of folds by germ band extension (*g*) using 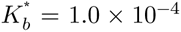, and (**e**) the timing of folding by germ band extension (*g*) using 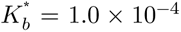 (right). **f–h**, Parameter sweep with mitotic domains and same plots as above. **i**, Parameter sweep for cephalic furrow simulations. Bending rigidity and cephalic furrow depth at different values of 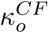 and germ band extension. Values above 0.2 exhibit a clear peak in bending energy for most conditions and the depth reaches a plateau across iterations. The cephalic furrow depth at the peak of bending energy (max bend) and at the final iteration is similar for simulations with 0% of germ band extension. At higher percentages of germ band extension the folds (both cephalic furrow and ectopic folds) exhibit a greater depth at the last iteration. **j**, Finegrained parameter sweep of ectopic folding without a delay in the formation of mitotic domains (*t_MD_* = 0). With simultaneous formation, only higher values of 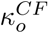 are effective in buffering the formation of ectopic folds around the cephalic furrow region. This is also limited to low percentages of germ band extension since at higher percentages there is an increase in the frequency of ectopic folding. **k**, Fine-grained parameter sweep of ectopic folding with a delay in the formation of mitotic domains (*t*_*MD*_ = 5). When a delay in mitotic formation is present, even low values of 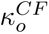 are effective in preventing the formation of ectopic folds. In this condition, the extension of the germ band increases the formation of ectopic folds, but only at the posterior regions close to the germ band tip. This suggests that the initiation of the cephalic furrow is crucial to its ability to buffer the ectopic folding. Values of 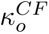 are shown in units of 1/*L*. *t_MD_* = 1 corresponds to 105 computational timesteps. **l**, Representative simulations from Fig. 4g at 0 and 20% of germ band extension.

**Figure Extended Data Fig. 6:**
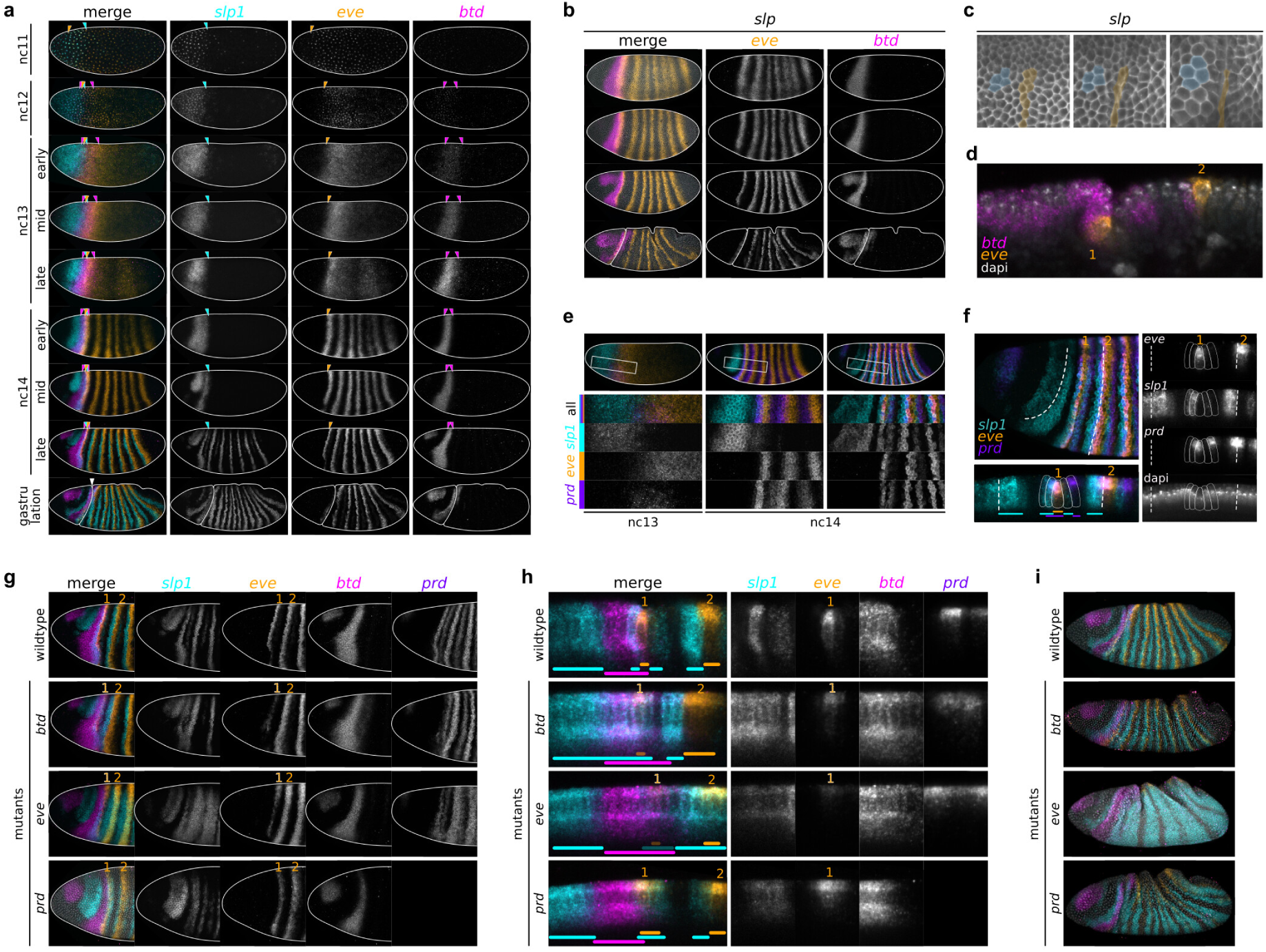
Genetic patterning of the head–trunk boundary in *Drosophila*. **a**, Expression of *slp1*, *eve*, and *btd* in wildtype from nuclear cycle (nc) 11 to gastrulation. **b**, Expression of *btd* and *eve* in *slp* mutants. **c**, Initiator cell behavior in *slp* mutants. **d**, Asymmetric cephalic furrow in *slp* mutants. **e**, Expression of *prd*, *slp1*, and *eve* in wildtype embryos in lateral view. **f**, Expression of *prd*, *slp1*, and *eve* in wildtype embryos with head and profile views. **g**, Disruption of *btd*, *eve*, and *slp1* expression patterns in the head of *btd*, *eve*, and *prd* mutant embryos. **h**, Profile views of **g** showing the gene expression at the head–trunk boundary cells. **i**, Lateral views of *btd*, *eve*, and *slp1* expression patterns in *btd*, *eve*, and *prd* mutants after gastrulation.

**Figure Extended Data Fig. 7:**
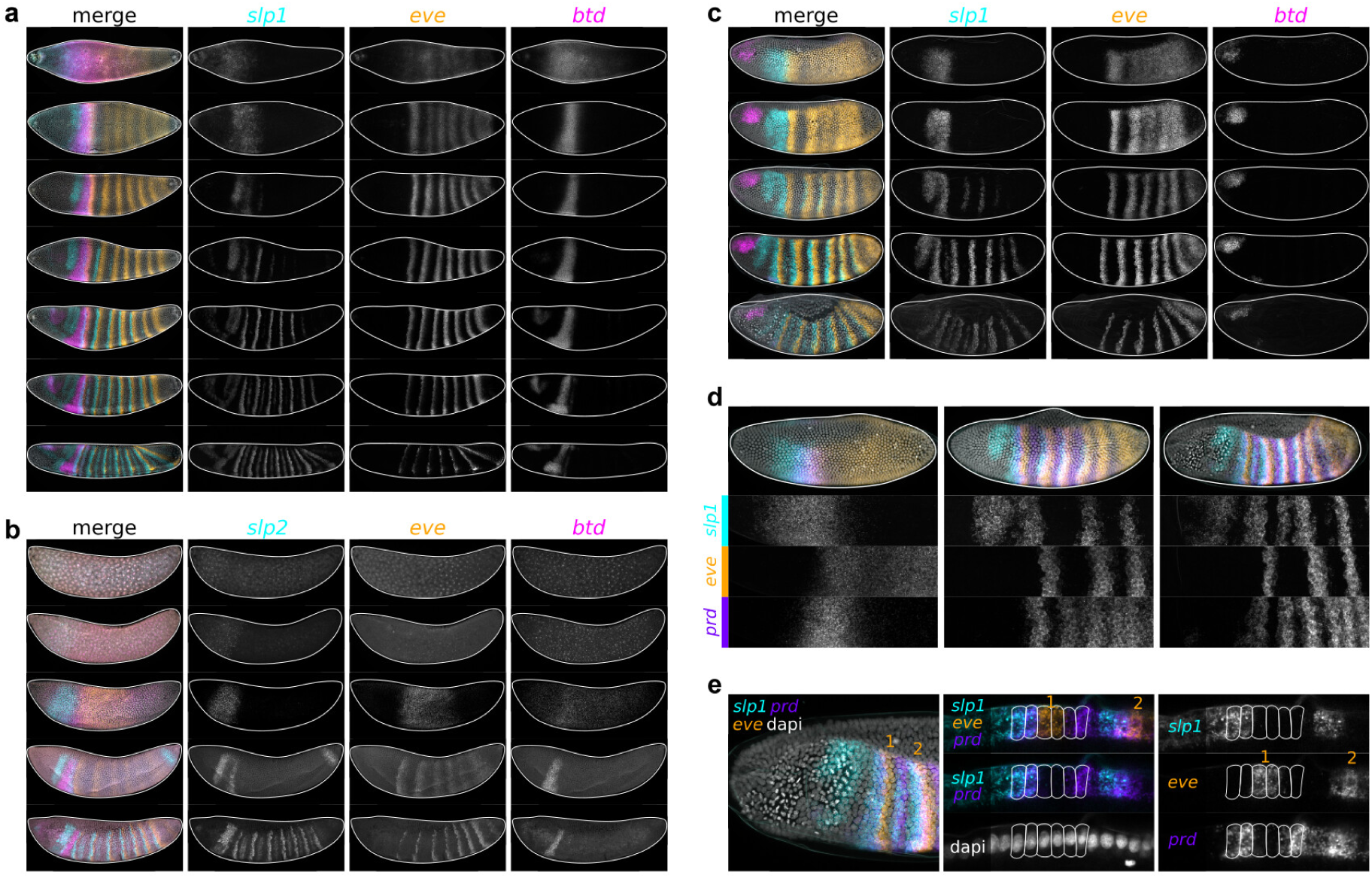
Genetic patterning of the head–trunk boundary in other dipteran species. **a**, Expression of *slp1*, *eve*, and *btd* in *Ceratitis* developmental stages. **b**, Expression of *slp2*, *eve*, and *btd* in *Anopheles* developmental stages. **c**, Expression of *slp1*, *eve*, and *btd* in *Clogmia* developmental stages. **d**, Expression of *slp1*, *eve*, and *prd* in *Clogmia* lateral views. **e**, Expression of *slp1*, *eve*, and *prd* in *Clogmia* at the head–trunk boundary in profile views.

## Supplementary information

**Figure Supplementary Fig. 1:**
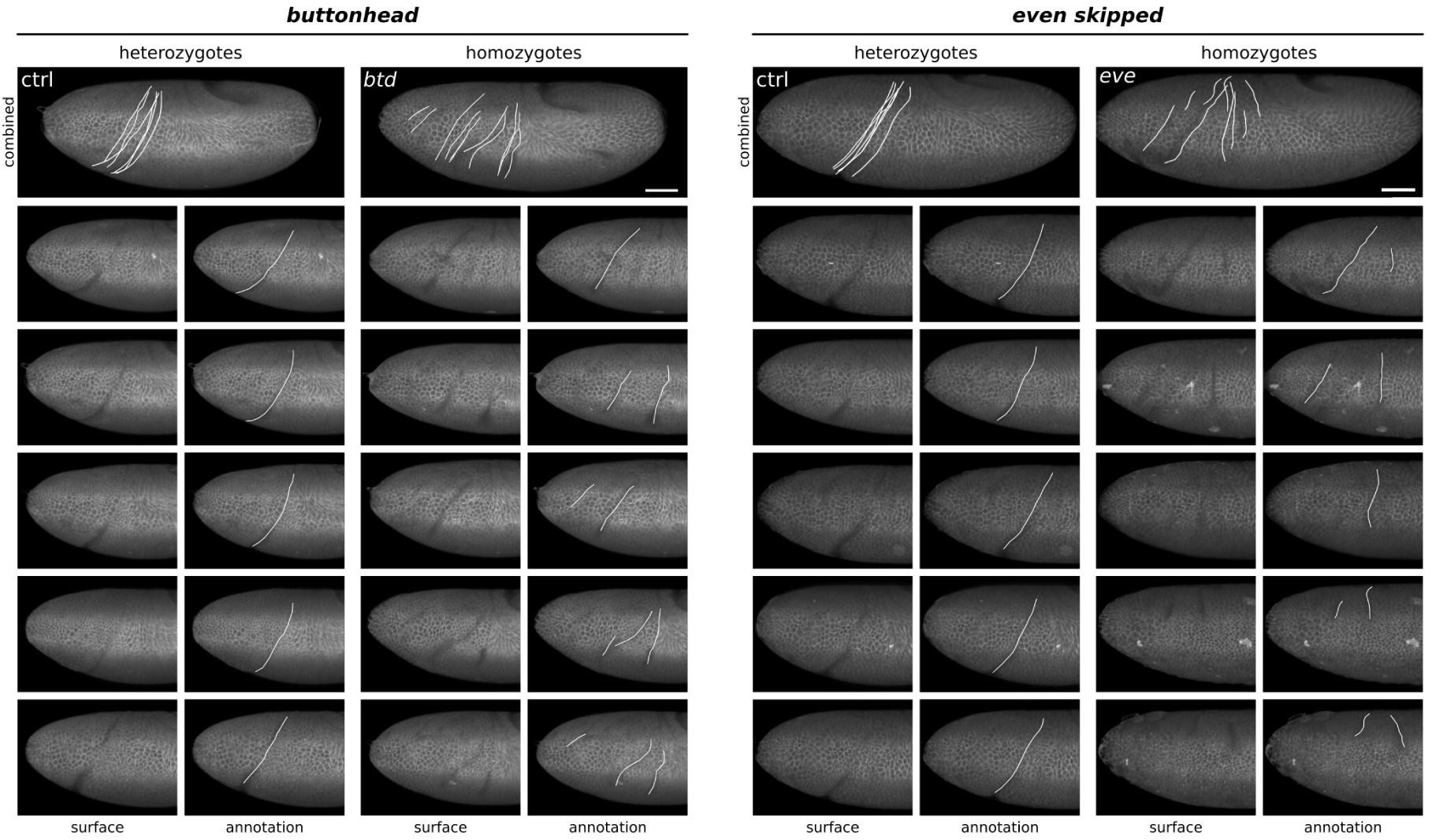
Individual variability of ectopic folding in *btd* and *eve* mutants. Cephalic furrow and ectopic folds are traced in white lines. The combined views illustrate the position of folding overlaid over a single embryo after embryo registration.

**Figure Supplementary Fig. 2:**
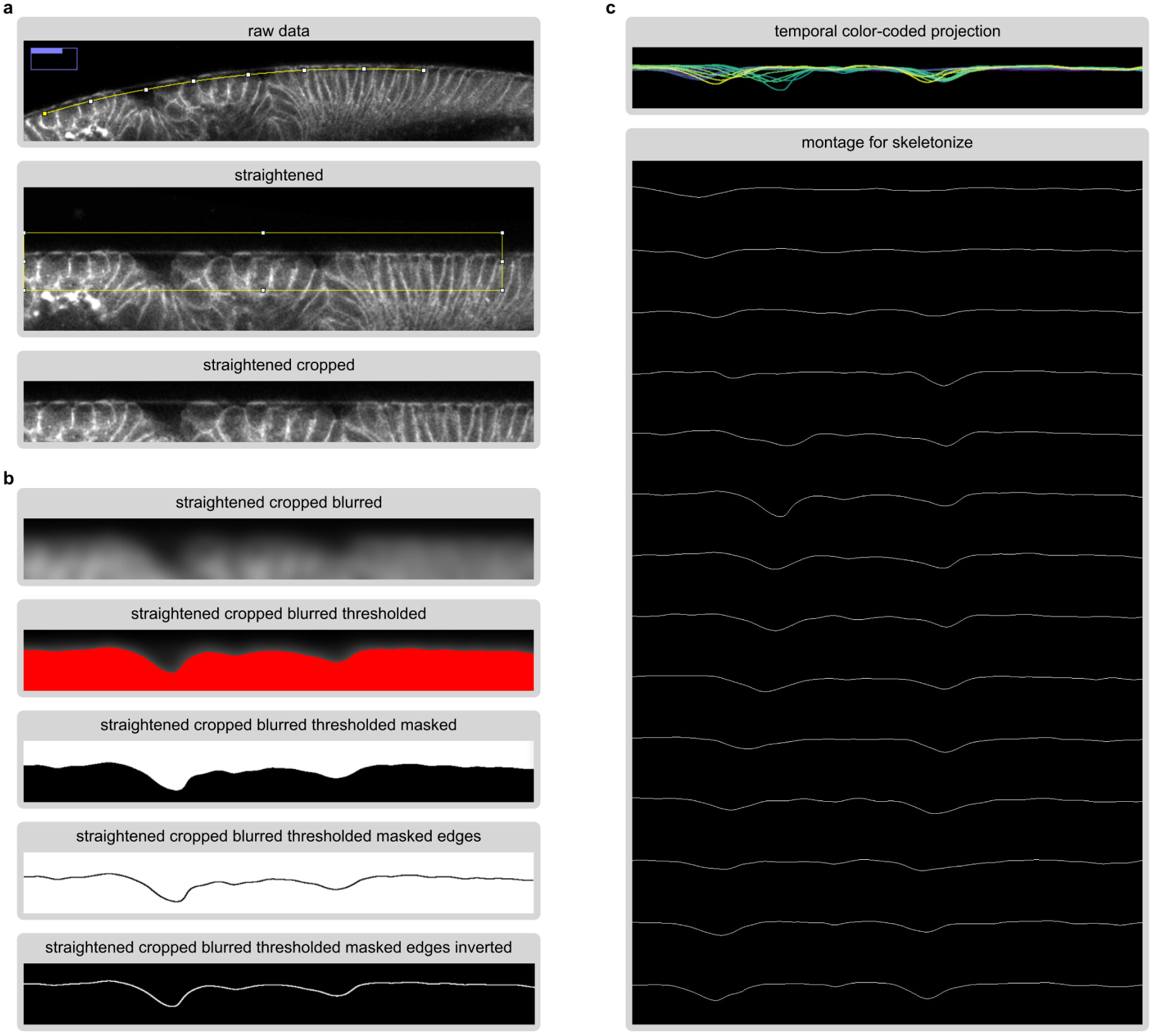
Image processing pipeline for the tortuosity analysis in cauterized mutants. **a**, We acquired a single slice in profile view of the head–trunk epithelium. First, we straightened the epithelial monolayer along the curvature of the vitelline envelope using the Straighten tool in ImageJ. We then cropped a window to standardize the size of the analyzed area for all embryos. **b**, Then, we applied a Gaussian blur which allows capturing the deformations in the epithelium caused by the ectopic folds after thresholding. We create a mask and detect edges and invert to retain a single pixel line corresponding to the outline of the epithelium. The image is inverted for downstream processing. **c**, We applied a temporal color-coded projections to visualize the epithelial dynamics over time, and created a montage with all timepoints to extract the length of the outline using the skeletonize plugin in ImageJ.

**Figure Supplementary Fig. 3:**
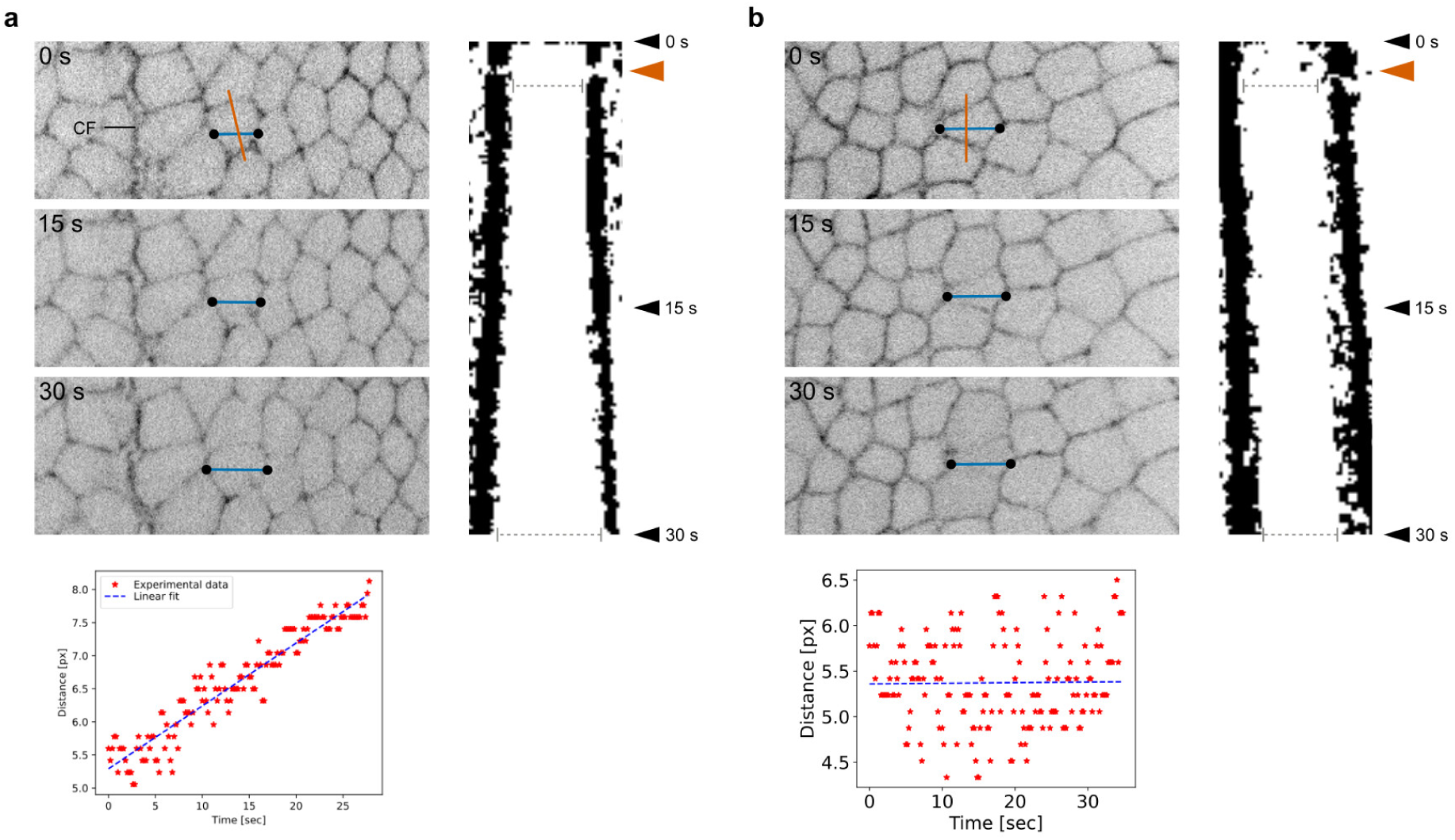
Image processing pipeline for the ablation analysis. **a**, Example of laser ablation near the cephalic furrow with the membrane signal (top left), the generated kymograph (right), and the linear fit over the distance between edges extracted from the kymograph (bottom left). The position of the laser cut is annotated in a vermilion line, the cell edges are marked in black circles, and the edge distances in a blue line. The distance between edges increases over time. **b**, Example of a laser ablation far from the cephalic furrow where the distance between edges does not increase over time.

**Figure Supplementary Video 1:**
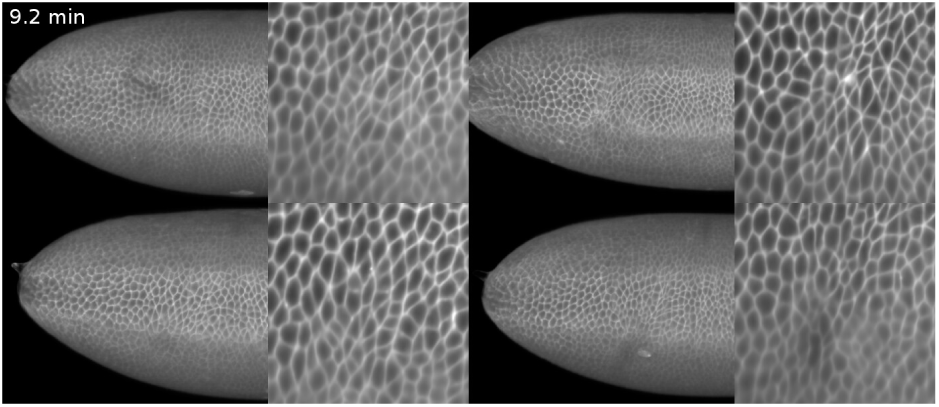
Reduced apical constriction in *btd* mutants. Lateral view (left) and cartographic projection (right) of the head–trunk interface in four individual *btd* mutants. Putative initiator cells (center) exhibit a reduced degree of apical constriction. The video is looped to highlight the changes in apical cell area. Frame rate = 10fps.

**Figure Supplementary Video 2:**
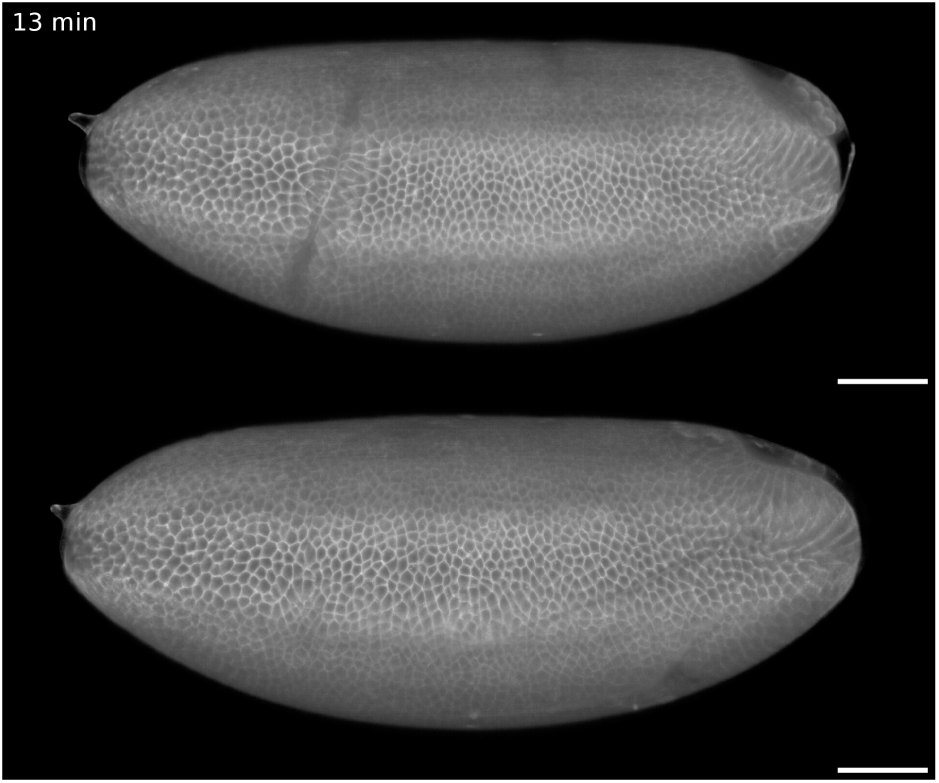
Lateral view of ectopic fold formation in *btd* mutant. The cephalic furrow forms normally in sibling controls (top) but is absent in *btd* mutants (bottom). In the mutant, no fold is present at the head–trunk interface until about 20 min, when a large ectopic fold appears and quickly unfolds at about 45 min. In the sibling control, the cephalic furrow remains partially invaginated for the period shown in the recording (about 110 min). Frame rate = 15 fps. Scale bars = 50 µm.

**Figure Supplementary Video 3:**
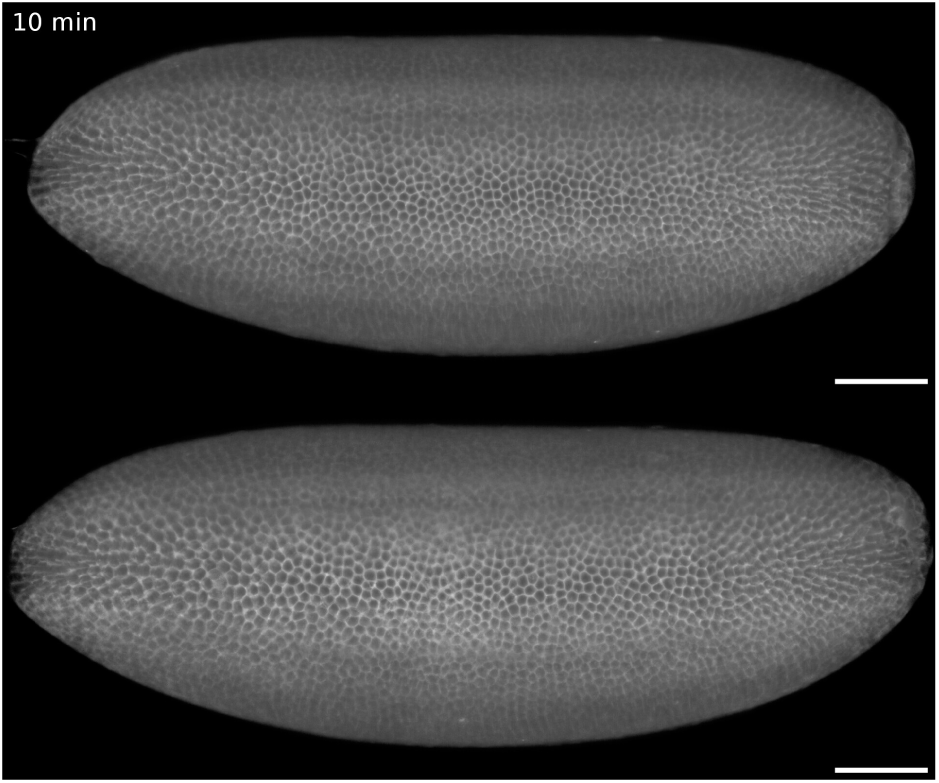
Lateral view of ectopic fold formation in *eve* mutant. The cephalic furrow forms normally in sibling controls (top) but is absent in *eve* mutants (bottom). There is no invagination at the head– trunk boundary at the onset of gastrulation, but an ectopic fold starts forming near the dorsal region as soon as the mitotic domains begin expanding around 24 min. The ectopic folds unfold almost entirely by the end of the recording (about 85 min). Additional ectopic folds appear in the trunk region. Frame rate = 10 fps. Scale bars = 50 µm.

**Figure Supplementary Video 4:**
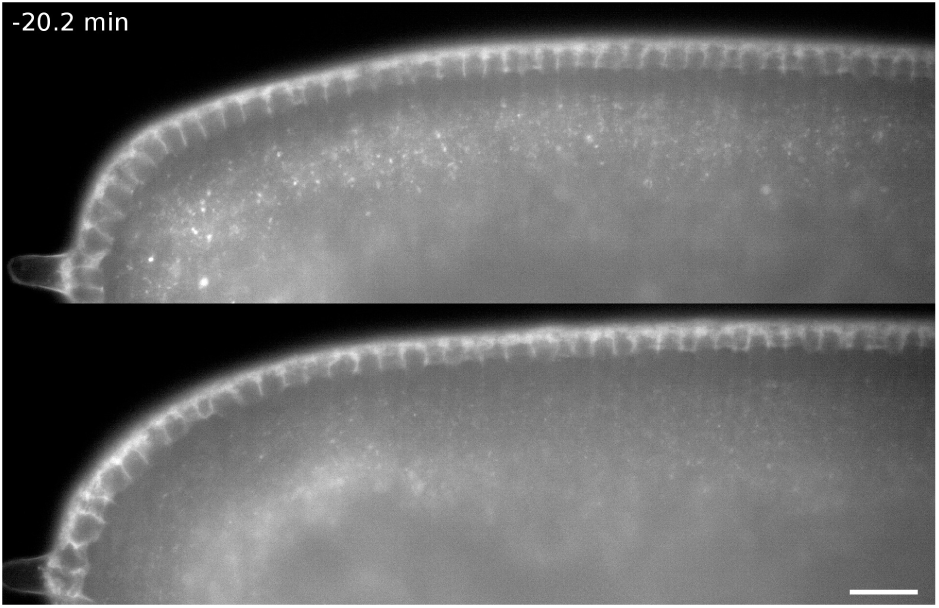
Profile view of ectopic fold formation in *btd* mutant. In sibling controls (top), the cephalic furrow initiates at the onset of gastrulation (1.5 min) and is fully invaginated when the cell divisions start (about 11 min). In *btd* mutants (bottom), no invagination initiates, but some embryos exhibit a bulging of the epithelium due to a residual apical constriction behavior (about 7 min). An ectopic fold forms at this position. Its morphology differs greatly from the cephalic furrow (see 10 min). Both the cephalic furrow and ectopic folds regress with the extension of the germ band. Frame rate = 10 fps. Scale bar = 20 µm.

**Figure Supplementary Video 5:**
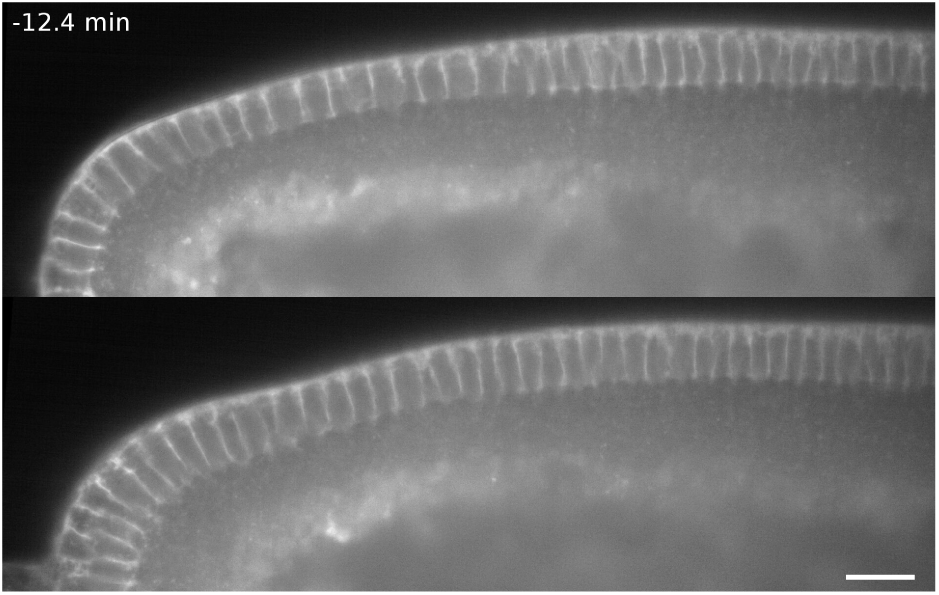
Profile view of ectopic fold formation in *eve* mutant. In sibling controls (top), the cephalic furrow initiates at the onset of gastrulation (1.8 min). In *eve* mutants, there are no folds appearing in the epithelium until the formation of mitotic domains (about 10 min). Then, a large ectopic fold appears posterior to dividing cells (15 min). The epithelium of *eve* mutants show additional folding events along the head and trunk regions. Frame rate = 10 fps. Scale bar = 20 µm.

**Figure Supplementary Video 6:**
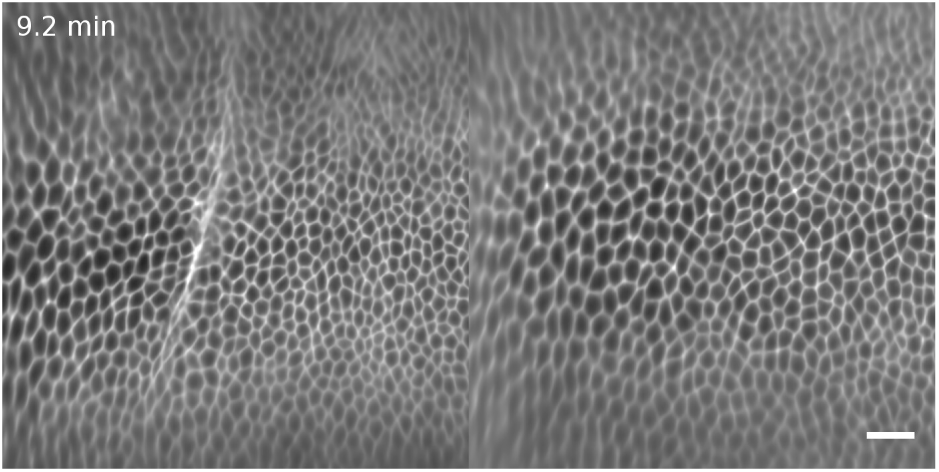
Ectopic folding between mitotic domains in *btd* mutant. Detailed view of cartographic projections of *btd* embryos, showing the formation of the cephalic furrow (left) and of an ectopic fold (right). In sibling controls, the cephalic furrow initiates from a narrow row of cells and invaginates in a progressive manner before the appearance of mitotic domains. In *btd* mutants, the ectopic folds only appear after the apical expansion of dividing cells within mitotic domains, quickly buckling and unfolding shortly after. Frame rate = 10 fps. Scale bar = 20 µm (approximate value for cartographic projection).

**Figure Supplementary Video 7:**
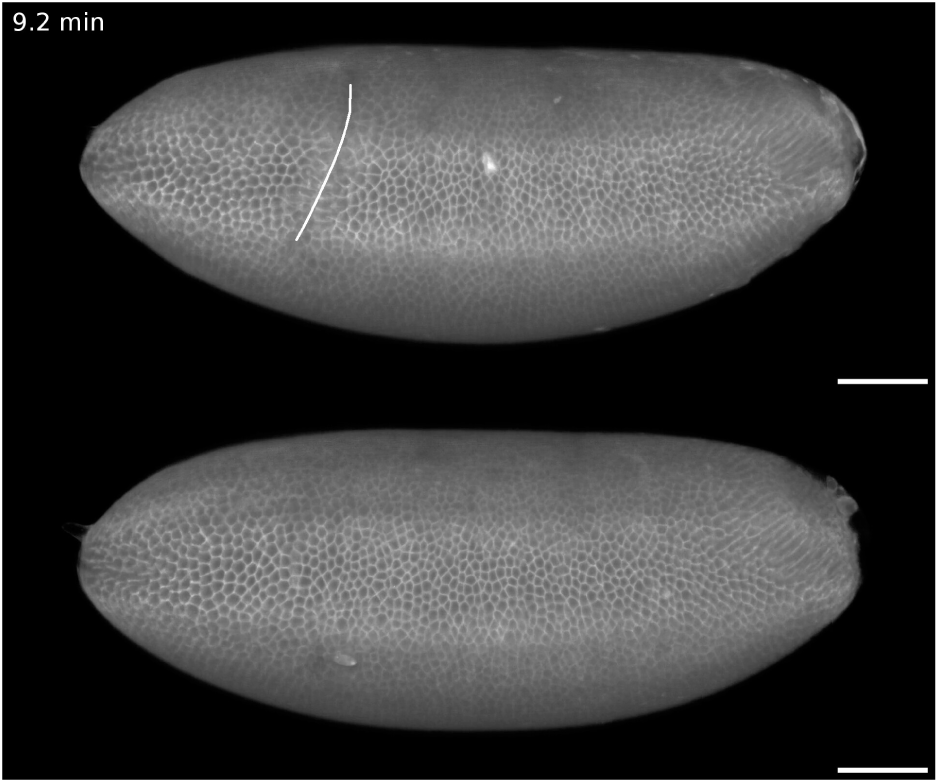
Dynamics of ectopic folding in *btd* mutant. The cephalic furrow in sibling controls (top) and the ectopic folds in *btd* mutants (bottom) are annotated in white to visualize the dynamics in position, extension, and shape during their formation. Frame rate = 10 fps. Scale bars = 50 µm.

**Figure Supplementary Video 8:**
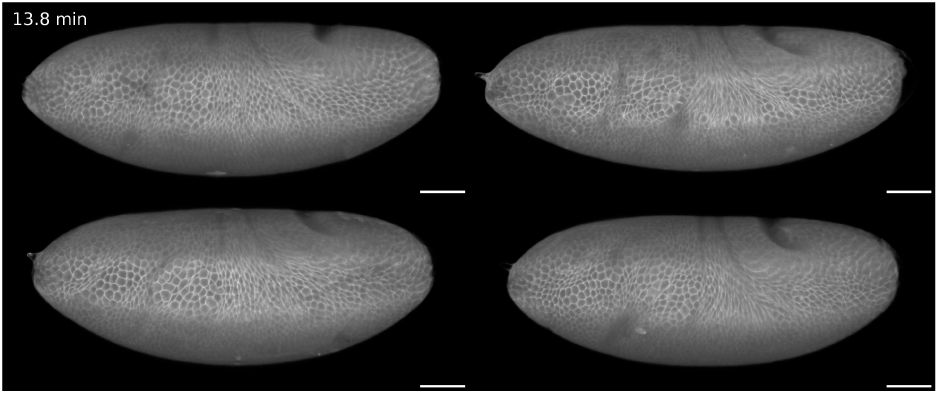
Variability of ectopic folding in *btd* mutants. The video shows four individual *btd* mutants, where each display a different pattern and number of ectopic folds at the head–trunk interface. The video is looped to highlight the dynamics of ectopic folding. Frame rate = 15 fps. Scale bars = 50 µm.

**Figure Supplementary Video 9:**
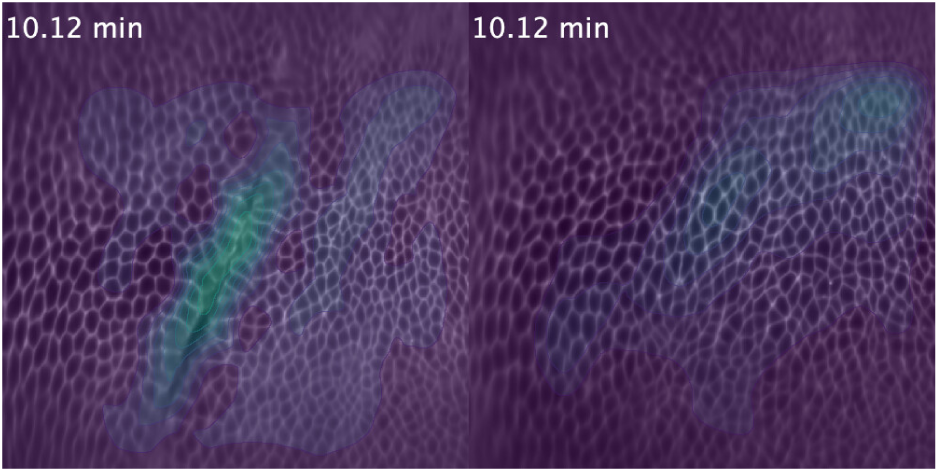
Epithelial strain rate during ectopic folding in *btd* mutant. Video from Supplementary Video 6 overlaid with the estimated strain rate across the tissues (color-coded from purple to yellow). Increase in strain rates are associated with tissue infolding and mitotic expansions. The video is looped. Frame rate = 10 fps.

**Figure Supplementary Video 10:**
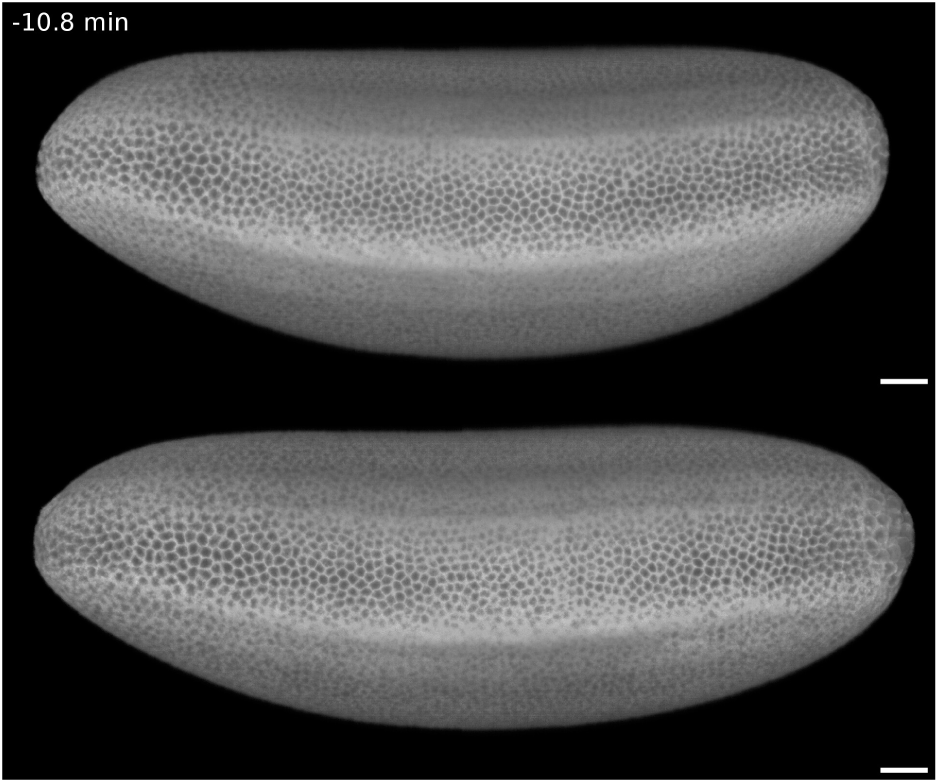
Lateral view of cephalic furrow formation in *stg* mutants. Sibling control (top) and *stg* mutant (bottom) during gastrulation. The formation of the cephalic furrow is almost identical to the control embryo. The other morphogenetic movements also occur normally until about 35 min. At this point, the cells in the *stg* mutant are notably larger than the control. Frame rate = 15 fps. Scale bars = 50 µm.

**Figure Supplementary Video 11:**
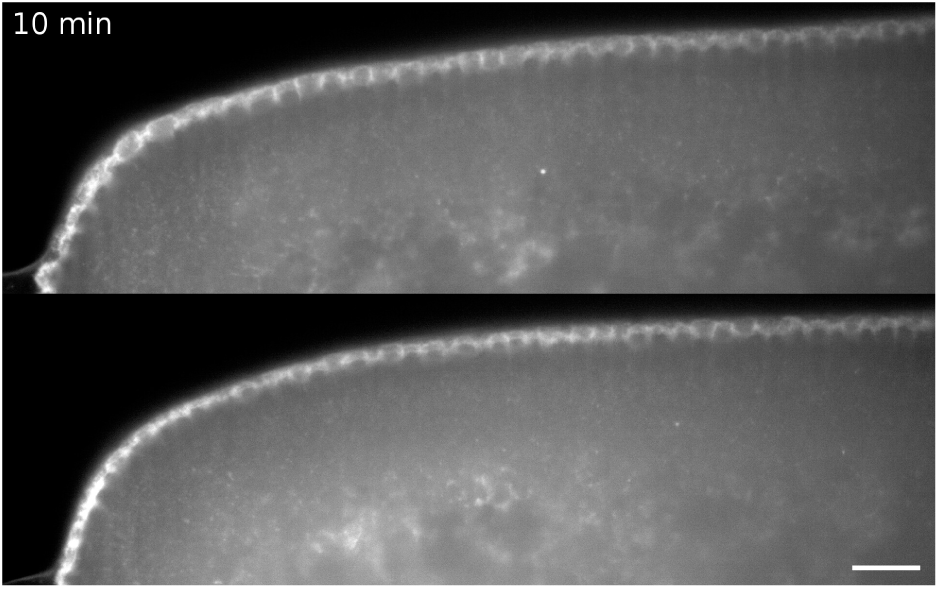
Dorsal view of cephalic furrow formation in *stg* mutants. Sibling control (top) and *stg* mutant (bottom) during gastrulation. The cephalic furrow in *stg* mutants initiates without delay and shows identical morphology to the control until cell divisions begin in the latter. The cells dividing within the cephalic furrow of control embryos alter its morphology, it becomes curved and lengthier. In contrast, the cephalic furrow in the *stg* mutant retains its initial morphology until it unfolds. Frame rate = 10 fps. Scale bar = 20 µm.

**Figure Supplementary Video 12:**
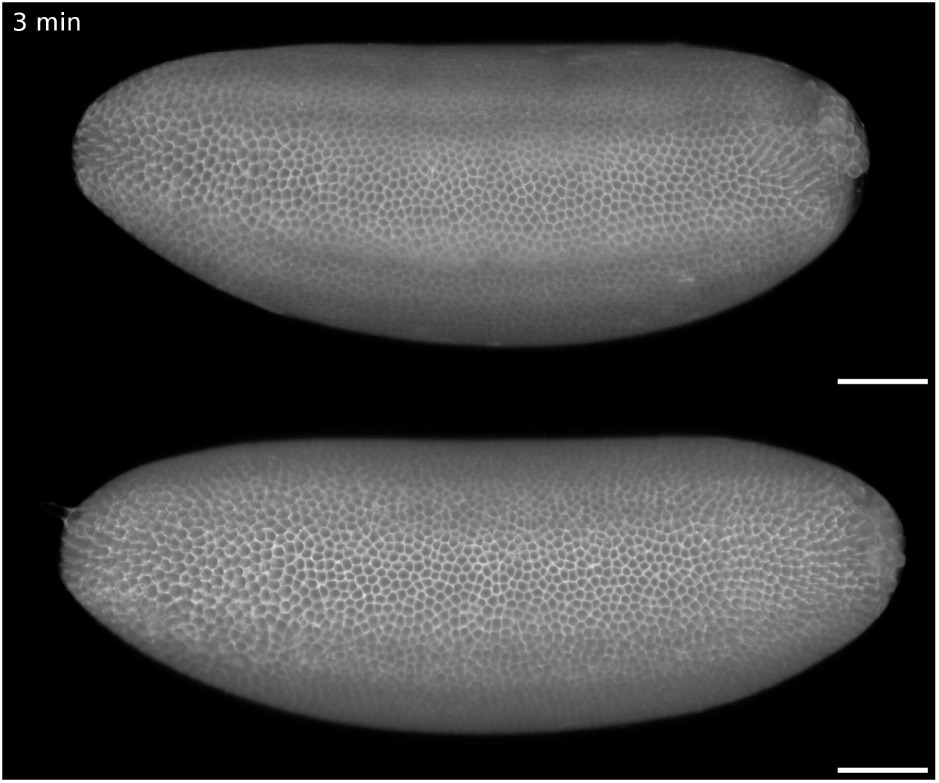
Lateral view of *btd*–*stg* double mutant. A *btd* homozygote (top) shows the formation of ectopic folds, while no ectopic folds form in the *btd*–*stg* double mutant (bottom). Frame rate = 10 fps. Scale bars = 50 µm.

**Figure Supplementary Video 13:**
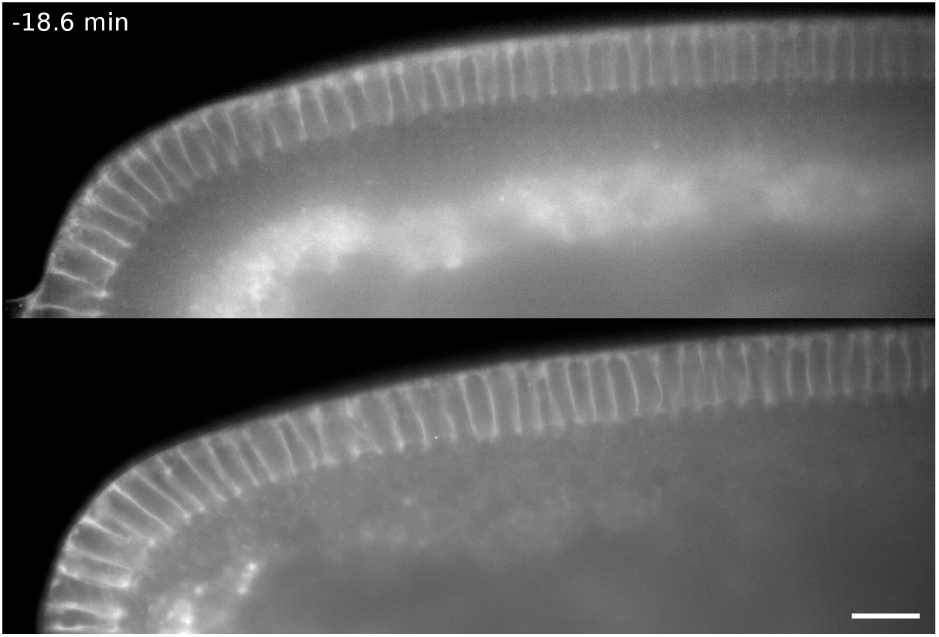
Dorsal view of *btd*–*stg* double mutant. A *btd* homozygote shows the formation of an ectopic fold (top). The *btd*–*stg* double mutant exhibits no mitotic domains and no ectopic folds (bottom). Frame rate = 10 fps. Scale bar = 20 µm.

**Figure Supplementary Video 14:**
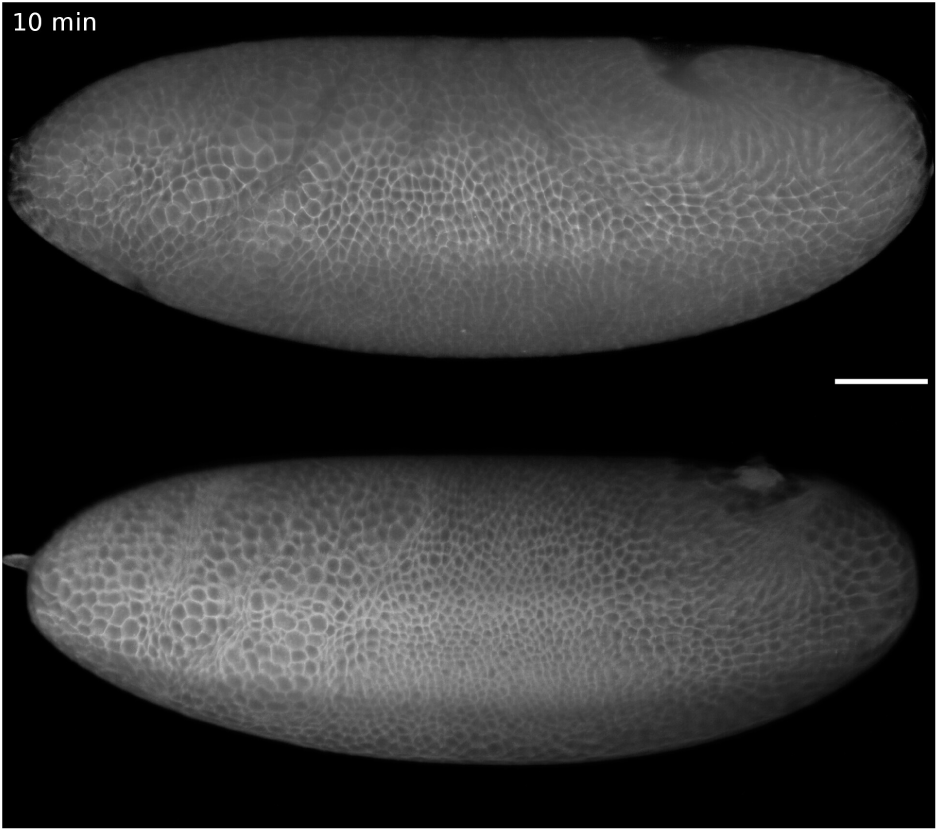
Lateral views of germ band cauterization in *eve* embryos. Non-cauterized *eve* embryo showing the formation of ectopic folds (top, same embryo from Supplementary Video 3) and a cauterized *eve* embryo where no ectopic folds appear at the head–trunk interface (bottom). The germ band extension is mechanically blocked by cauterizing the tissue to the vitelline envelope. Mitotic domains form normally, but no folding of the surface occurs. Frame rate = 10 fps. Scale bar = 50 µm.

**Figure Supplementary Video 15:**
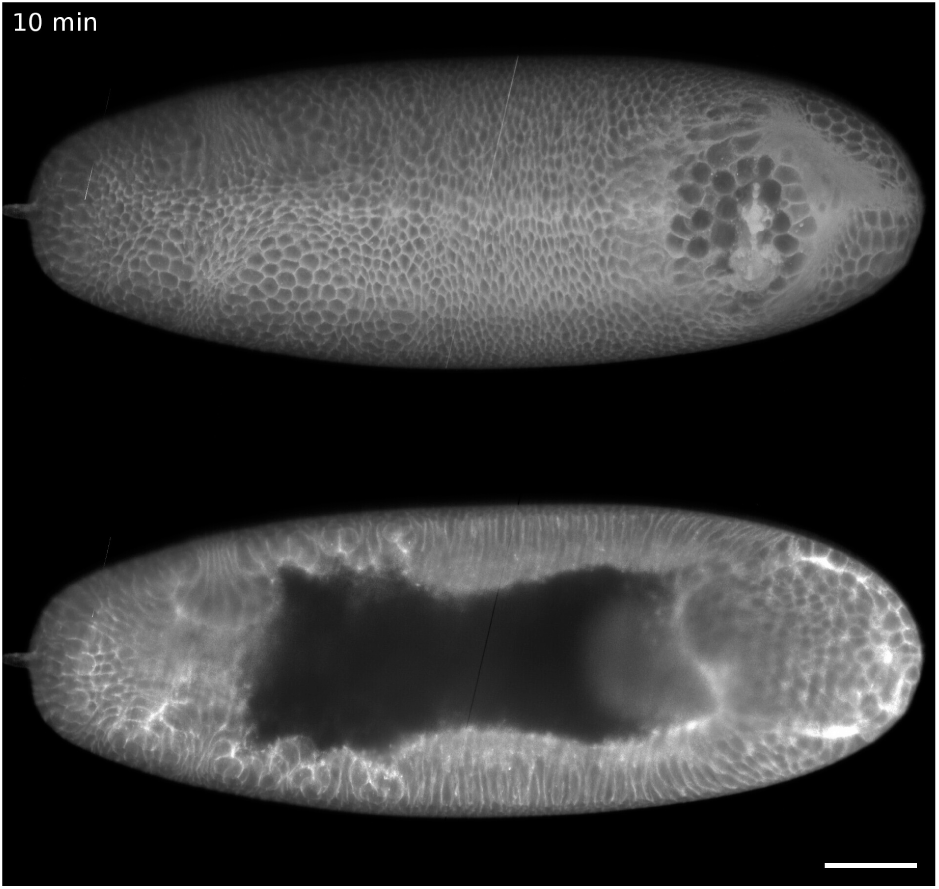
Profile views of germ band cauterization in *eve* mutant. Same embryo from Supplementary Video 14, but showing a surface and a profile view. The cauterization prevents the extension of the germ band. The mitotic domains compress non-dividing cells, but these do not buckle. Frame rate = 10 fps. Scale bar = 50 µm.

**Figure Supplementary Video 16:**
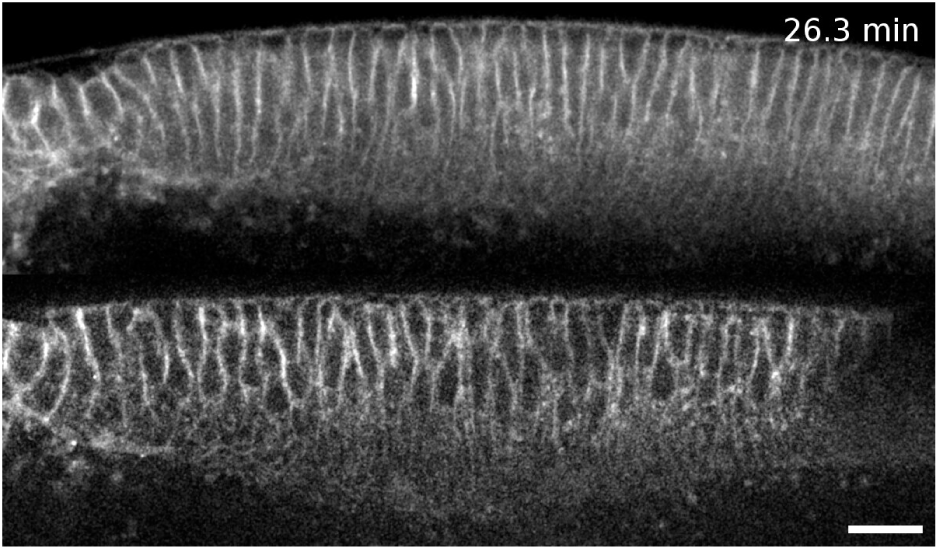
Profile views of germ band cauterizations in *btd* embryos. A non-cauterized *btd* embryo (top) showing ectopic folds and a cauterized *btd* embryo showing no ectopic folds (bottom). Frame rate = 10 fps. Scale bar = 20 µm.

**Figure Supplementary Video 17:**
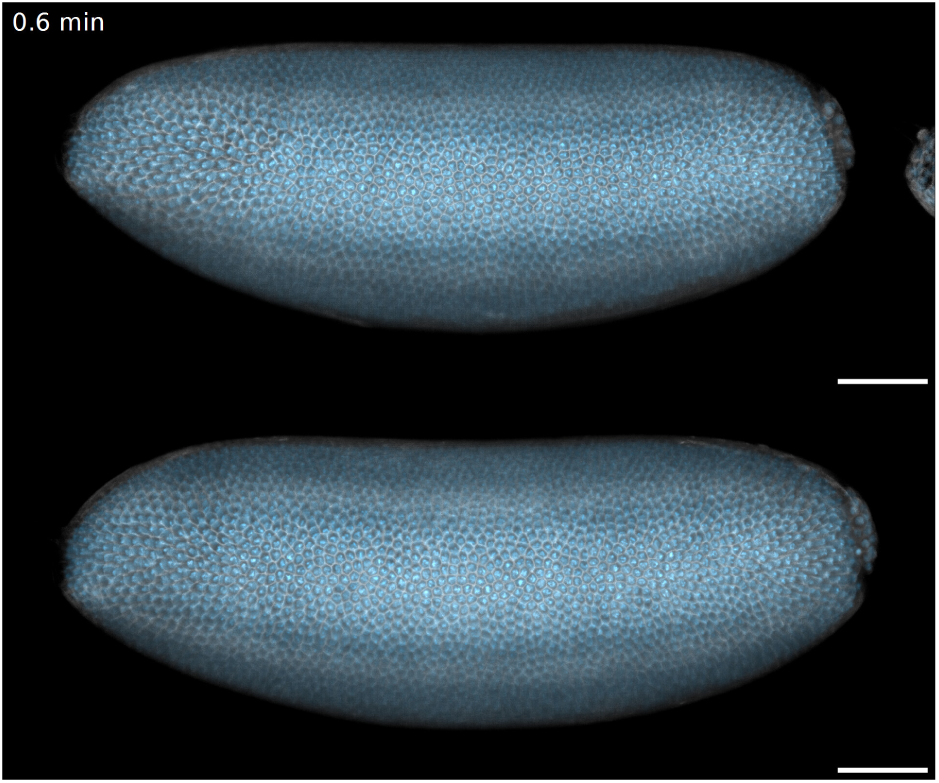
Lateral view of *slp* mutant. A *slp* heterozygote (top) shows a normal cephalic furrow formation. The *slp* homozygote (bottom) shows a delayed initiator cell behavior and cephalic furrow formation. The embryo also exhibits a more prominent posterior dorsal fold and an ectopic fold forming within the mitotic domain 6 before cell divisions. Frame rate = 10 fps. Scale bar = 50 µm.

## Supplementary Note 1

### Comparing our reference bending rigidity to direct measurements

Here we compare the reference bending rigidity established for the *Drosophila* blastoderm with our experiments to direct measurements from the literature. Trushko et al.^17^ reports that in a MDCK monolayer 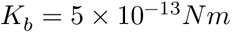 and 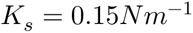. With these values, we can compute the dimensionless bending rigidity 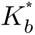 . However, the thickness of the MDCK monolayer and the *Drosophila* blastoderm are very different, and hence we need to correct for this before computing 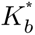 for MDCK monolayer. The bending rigidity scales with the square of the thickness of the tissue. Hence, the corrected *K*_*b*_ can be computed as 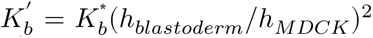. This gives us 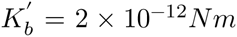. Now we can compute 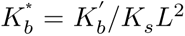 where we put *L* = 25*μm.* Here we scale 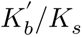 by the relevant lengthscale which is the semi-major axis of the *Drosophila* embryo. This gives us 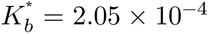. This is the estimated 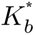 for the MDCK monolayer that has the same thickness as the *Drosophila* blastoderm (referring to the height correction) and has been put in the same geometrical conditions as the *Drosophila* blastoderm (referring to the *L* used in the calculation).

## Supplementary Note 2

### Live-imaging screen for cephalic furrow genes

To uncover other genes directly involved in cephalic furrow formation in addition to *btd*, *eve*, and *prd*, we performed a live-imaging screen in strains containing loss-of-function alleles for a selection of candidate genes expressed at the head–trunk region.^65–67^ Because the cephalic furrow is transient and leaves no trace, the live-imaging approach is critical to recognize altered phenotypes. From about 50 genes, we only detected three showing abnormal cephalic furrow formation to different degrees besides the previously described genes (see Table Supplementary Table 1). The strongest cephalic furrow phenotype was present in flies mutant for the *sloppy paired* (*slp*) genes.

---

**Table Supplementary Table 1:**
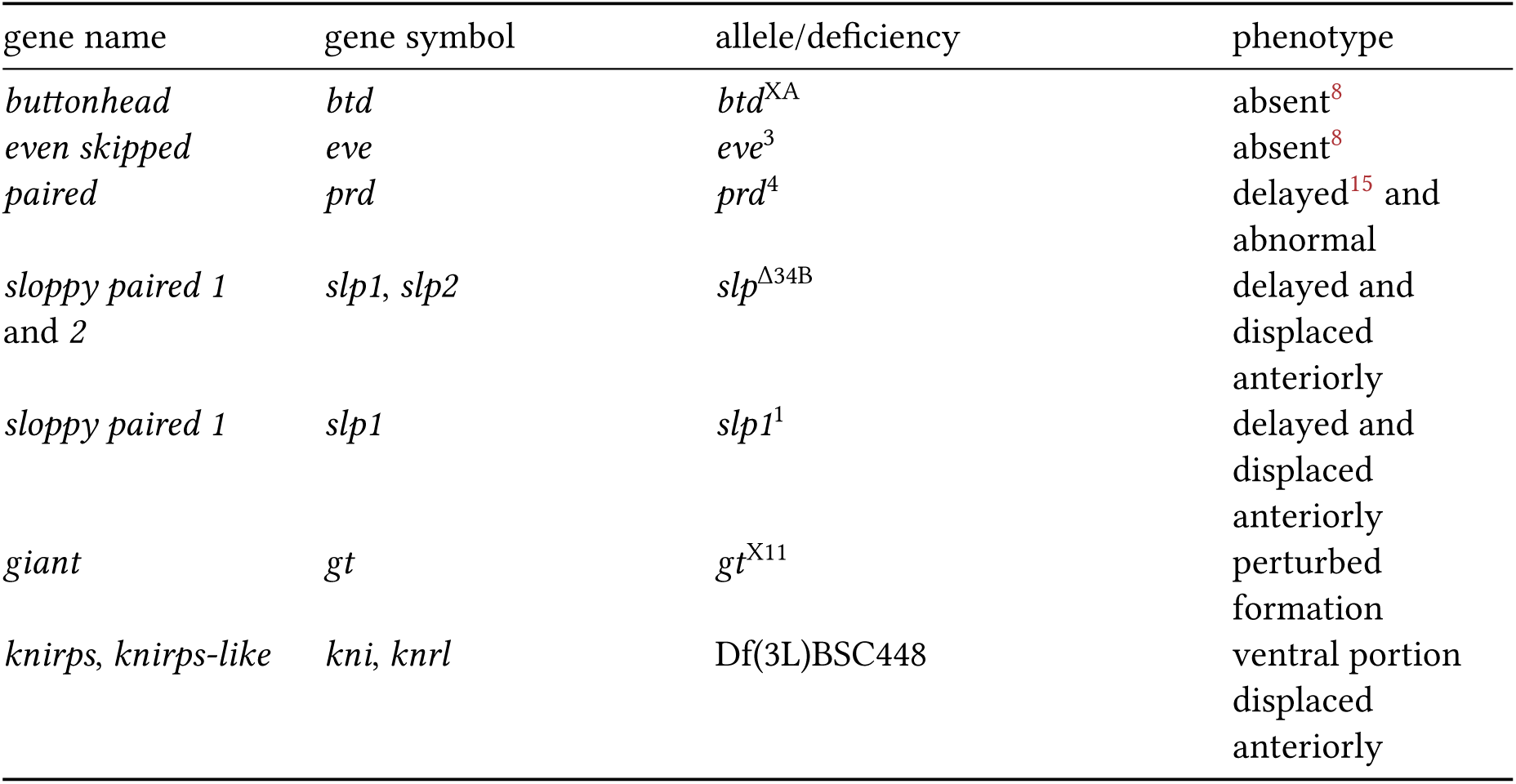
Summary of live-imaging screening results for cephalic furrow genes.

## References

1. Keller, R. Physical biology returns to morphogenesis. Science 338, 201–203 (2012).

2. Heisenberg, C.-P. & Bellaïche, Y. Forces in tissue morphogenesis and patterning. Cell 153, 948–962 (2013).

3. Collinet, C. & Lecuit, T. Programmed and self-organized flow of information during morpho-genesis. Nat. Rev. Mol. Cell Biol. 22, 245–265 (2021).

4. Hartenstein, V. & Campos-Ortega, J. A. Fate-mapping in wild-type *Drosophila melanogaster*. Wilhelm Roux Arch. Entwickl. Mech. Org. 194, 181–195 (1985).

5. Spencer, A. K., Siddiqui, B. A. & Thomas, J. H. Cell shape change and invagination of the cephalic furrow involves reorganization of F-actin. Dev. Biol. 402, 192–207 (2015).

6. Foe, V. E. Mitotic domains reveal early commitment of cells in *Drosophila* embryos. Development 107, 1–22 (1989).

7. Liu, F., Morrison, A. H. & Gregor, T. Dynamic interpretation of maternal inputs by the *Drosophila* segmentation gene network. Proc. Natl. Acad. Sci. U. S. A. 110, 6724–6729 (2013).

8. Vincent, A., Blankenship, J. T. & Wieschaus, E. Integration of the head and trunk segmentation systems controls cephalic furrow formation in *Drosophila*. Development 124, 3747–3754 (1997).

9. Eritano, A. S., Bromley, C. L., Bolea Albero, A., Schütz, L., Wen, F.-L., Takeda, M., Fukaya, T., Sami, M. M., Shibata, T., Lemke, S. & Wang, Y.-C. Tissue-scale mechanical coupling reduces morphogenetic noise to ensure precision during epithelial folding. Dev. Cell 53, 212–228.e12 (2020).

10. Popkova, A., Andrenšek, U., Pagnotta, S., Ziherl, P., Krajnc, M. & Rauzi, M. A mechanical wave travels along a genetic guide to drive the formation of an epithelial furrow during *Drosophila* gastrulation. Dev. Cell 59, 400–414.e5 (2024).

11. Turner, F. R. & Mahowald, A. P. Scanning electron microscopy of *Drosophila melanogaster* embryogenesis. II. Gastrulation and segmentation. Dev. Biol. 57, 403–416 (1977).

12. Costa, M., Sweeton, D. & Wieschaus, E. in The development of Drosophila melanogaster (eds. Bate, M. & Arias, A. M.) 1, 425–465 (Cold Spring Harbor Laboratory Press, 1993).

13. Dicko, M., Saramito, P., Blanchard, G. B., Lye, C. M., Sanson, B. & Étienne, J. Geometry can provide long-range mechanical guidance for embryogenesis. PLoS Comput. Biol. 13, e1005443 (2017).

14. Dey, B., Kaul, V., Kale, G., Scorcelletti, M., Takeda, M., Wang, Y.-C. & Lemke, S. Divergent evolutionary strategies preempt tissue collision in fly gastrulation. bioRxiv (2023). doi:10.1101/2023.10.09.561568

15. Blankenship, J. T. & Wieschaus, E. Two new roles for the *Drosophila* AP patterning system in early morphogenesis. Development 128, 5129–5138 (2001).

16. Edgar, B. A. & O’Farrell, P. H. Genetic control of cell division patterns in the *Drosophila* embryo. Cell 57, 177–187 (1989).

17. Trushko, A., Di Meglio, I., Merzouki, A., Blanch-Mercader, C., Abuhattum, S., Guck, J., Alessandri, K., Nassoy, P., Kruse, K., Chopard, B. & Roux, A. Buckling of an epithelium growing under spherical confinement. Dev. Cell 54, 655–668.e6 (2020).

18. Fouchard, J., Wyatt, T. P. J., Proag, A., Lisica, A., Khalilgharibi, N., Recho, P., Suzanne, M., Kabla, A. & Charras, G. Curling of epithelial monolayers reveals coupling between active bending and tissue tension. Proc. Natl. Acad. Sci. U. S. A. 117, 9377–9383 (2020).

19. Efrati, E., Sharon, E. & Kupferman, R. Elastic theory of unconstrained non-euclidean plates. J. Mech. Phys. Solids 57, 762–775 (2009).

20. Andrioli, L. P., Oberstein, A. L., Corado, M. S. G., Yu, D. & Small, S. Groucho-dependent repression by Sloppy-paired 1 differentially positions anterior pair-rule stripes in the *Drosophila* embryo. Dev. Biol. 276, 541–551 (2004).

21. Andrioli, L. P., Digiampietri, L. A., Barros, L. P. de & Machado-Lima, A. Huckebein is part of a combinatorial repression code in the anterior blastoderm. Dev. Biol. 361, 177–185 (2012).

22. Strobl, F., Schetelig, M. F. & Stelzer, E. H. K. In toto light sheet fluorescence microscopy live imaging datasets of *ceratitis capitata* embryonic development. Sci. Data 9, 340 (2022).

23. Strobl, F., Schmitz, A., Schetelig, M. F. & Stelzer, E. H. K. A two-level staging system for the embryonic morphogenesis of the mediterranean fruit fly (medfly) ceratitis capitata. PLoS One 19, e0316391 (2024).

24. Cheatle Jarvela, A. M., Trelstad, C. S. & Pick, L. Regulatory gene function handoff allows essential gene loss in mosquitoes. Commun. Biol. 3, 540 (2020).

25. Cheatle Jarvela, A. M., Trelstad, C. S. & Pick, L. Anterior-posterior patterning of segments in *Anopheles stephensi* offers insights into the transition from sequential to simultaneous segmentation in holometabolous insects. J. Exp. Zool. B Mol. Dev. Evol. 340, 116–130 (2023).

26. Jiménez-Guri, E., Wotton, K. R., Gavilán, B. & Jaeger, J. A staging scheme for the development of the moth midge *Clogmia albipunctata*. PLoS One 9, e84422 (2014).

27. Nelson, C. M. On buckling morphogenesis. J. Biomech. Eng. 138, 021005 (2016).

28. Gupta, V. K., Nam, S., Yim, D., Camuglia, J., Martin, J. L., Sanders, E. N., O’Brien, L. E., Martin, A. C., Kim, T. & Chaudhuri, O. The nature of cell division forces in epithelial monolayers. J. Cell Biol. 220, (2021).

29. Ko, C. S., Kalakuntla, P. & Martin, A. C. Apical constriction reversal upon mitotic entry underlies different morphogenetic outcomes of cell division. Mol. Biol. Cell 31, 1663–1674 (2020).

30. Kondo, T. & Hayashi, S. Mitotic cell rounding accelerates epithelial invagination. Nature 494, 125–129 (2013).

31. Freddo, A. M., Shoffner, S. K., Shao, Y., Taniguchi, K., Grosse, A. S., Guysinger, M. N., Wang, S., Rudraraju, S., Margolis, B., Garikipati, K., Schnell, S. & Gumucio, D. L. Coordination of signaling and tissue mechanics during morphogenesis of murine intestinal villi: A role for mitotic cell rounding. Integr. Biol. 8, 918–928 (2016).

32. Bao, L., Locovei, S. & Dahl, G. Pannexin membrane channels are mechanosensitive conduits for ATP. FEBS Lett. 572, 65–68 (2004).

33. Locovei, S., Wang, J. & Dahl, G. Activation of pannexin 1 channels by ATP through P2Y receptors and by cytoplasmic calcium. FEBS Lett. 580, 239–244 (2006).

34. Shah, P., Hobson, C. M., Cheng, S., Colville, M. J., Paszek, M. J., Superfine, R. & Lammerding, J. Nuclear deformation causes DNA damage by increasing replication stress. Curr. Biol. 31,753–765.e6 (2021).

35. Newman, S. A. & Müller, G. B. Epigenetic mechanisms of character origination. J. Exp. Zool. 288, 304–317 (2000).

36. Gramates, L. S., Agapite, J., Attrill, H., Calvi, B. R., Crosby, M. A., Dos Santos, G., Goodman, J. L., Goutte-Gattat, D., Jenkins, V. K., Kaufman, T., Larkin, A., Matthews, B. B., Millburn, G., Strelets, V. B. & the FlyBase Consortium. FlyBase: A guided tour of highlighted features. Genetics 220, iyac035 (2022).

37. Vellutini, B. C., Cuenca, M. B., Krishna, A., Szałapak, A., Modes, C. D. & Tomančák, P. Repository for: Patterned embryonic invagination evolved in response to mechanical instability. (2023). doi:10.5281/zenodo.7781947

38. Sander, K. Experimental egg activation in lower dipterans (Psychoda, Smittia) by low osmolality. Int. J. Invertebr. Reprod. Dev. 8, 175–183 (1985).

39. García-Solache, M., Jaeger, J. & Akam, M. A systematic analysis of the gap gene system in the moth midge *Clogmia albipunctata*. Dev. Biol. 344, 306–318 (2010).

40. Choi, H. M. T., Schwarzkopf, M., Fornace, M. E., Acharya, A., Artavanis, G., Stegmaier, J., Cunha, A. & Pierce, N. A. Third-generation *in situ* hybridization chain reaction: Multiplexed, quantitative, sensitive, versatile, robust. Development 145, (2018).

41. Schmied, C. & Tomancak, P. in Drosophila: Methods and protocols (ed. Dahmann, C.) 189–202 (Springer New York, 2016). doi:10.1007/978-1-4939-6371-3_10

42. Schindelin, J., Arganda-Carreras, I., Frise, E., Kaynig, V., Longair, M., Pietzsch, T., Preibisch, S., Rueden, C., Saalfeld, S., Schmid, B., Tinevez, J.-Y., White, D. J., Hartenstein, V., Eliceiri, K., Tomancak, P. & Cardona, A. Fiji: An open-source platform for biological-image analysis. Nat. Methods 9, 676–682 (2012).

43. Rueden, C. T., Schindelin, J., Hiner, M. C., DeZonia, B. E., Walter, A. E., Arena, E. T. & Eliceiri, K. W. ImageJ2: ImageJ for the next generation of scientific image data. BMC Bioinformatics 18, 529 (2017).

44. Schmid, B., Tripal, P., Fraaß, T., Kersten, C., Ruder, B., Grüneboom, A., Huisken, J. & Palmisano, R. 3Dscript: Animating 3D/4D microscopy data using a natural-language-based syntax. Nat. Methods 16, 278–280 (2019).

45. Weigert, M., Schmidt, U., Boothe, T., Müller, A., Dibrov, A., Jain, A., Wilhelm, B., Schmidt, D., Broaddus, C., Culley, S., Rocha-Martins, M., Segovia-Miranda, F., Norden, C., Henriques, R., Zerial, M., Solimena, M., Rink, J., Tomancak, P., Royer, L., Jug, F. & Myers, E. W. Content-aware image restoration: Pushing the limits of fluorescence microscopy. Nat. Methods 15, 1090–1097 (2018).

46. Heemskerk, I. & Streichan, S. J. Tissue cartography: Compressing bio-image data by dimensional reduction. Nat. Methods 12, 1139–1142 (2015).

47. Matlab. MATLAB. (The MathWorks Inc., 2015). at <https://www.mathworks.com/products/matlab.html>

48. Berg, S., Kutra, D., Kroeger, T., Straehle, C. N., Kausler, B. X., Haubold, C., Schiegg, M., Ales, J., Beier, T., Rudy, M., Eren, K., Cervantes, J. I., Xu, B., Beuttenmueller, F., Wolny, A., Zhang, C., Koethe, U., Hamprecht, F. A. & Kreshuk, A. ilastik: Interactive machine learning for (bio)image analysis. Nat. Methods (2019). doi:10.1038/s41592-019-0582-9

49. Vellutini, B. C. How to make cartographic projections using ImSAnE. (Zenodo, 2022). doi:10.5281/zenodo.7628299

50. Schmied, C. & Jambor, H. K. Effective image visualization for publications - a workflow using open access tools and concepts. F1000Res. 9, 1373 (2020).

51. Arganda-Carreras, I., Sorzano, C. O. S., Marabini, R., Carazo, J. M., Ortiz-de-Solorzano, C. & Kybic, J. in Computer vision approaches to medical image analysis 85–95 (Springer Berlin Heidelberg, 2006). doi:10.1007/11889762_8

52. Legland, D., Arganda-Carreras, I. & Andrey, P. MorphoLibJ: Integrated library and plugins for mathematical morphology with ImageJ. Bioinformatics 32, 3532–3534 (2016).

53. Arganda-Carreras, I., Fernández-González, R., Muñoz-Barrutia, A. & Ortiz-De-Solorzano, C. 3D reconstruction of histological sections: Application to mammary gland tissue. Microsc. Res. Tech. 73, 1019–1029 (2010).

54. Tseng, Q., Duchemin-Pelletier, E., Deshiere, A., Balland, M., Guillou, H., Filhol, O. & Théry, M. Spatial organization of the extracellular matrix regulates cell-cell junction positioning. Proc. Natl. Acad. Sci. U. S. A. 109, 1506–1511 (2012).

55. Krishna, A., Szałapak, A., Vellutini, B. C., Cuenca, M. B., Tomančák, P. & Modes, C. D. Model and simulations for: Patterned embryonic invagination evolved in response to mechanical instability. (2023). doi:10.5281/zenodo.7784906

56. The Inkscape Project. Inkscape: Open source scalable vector graphics editor. (2003). at <https://inkscape.org>

57. HandBrake Team. HandBrake: HandBrake’s main development repository. (Github, 2003). At <https://github.com/HandBrake/HandBrake>

58. Vellutini, B. C., Cuenca, M. B., Krishna, A., Szałapak, A., Modes, C. D. & Tomančák, P. Figures and videos for: Patterned embryonic invagination evolved in response to mechanical instability. (2023). doi:10.5281/zenodo.7781916

59. R Core Team. R: A language and environment for statistical computing. (R Foundation for Statistical Computing, 1993). at <http://www.r-project.org>

60. RStudio Team. RStudio: Integrated development environment for r. (RStudio, PBC., 2011). At <http://www.rstudio.com>

61. Granger, B. E. & Perez, F. Jupyter: Thinking and storytelling with code and data. Comput. Sci. Eng. 23, 7–14 (2021).

62. Campos-Ortega, J. A. & Hartenstein, V. The embryonic development of Drosophila melanogaster. (Springer Berlin Heidelberg, 1985). doi:10.1007/978-3-662-02454-6

63. Ashburner, M., Golic, K. G. & Hawley, R. S. Drosophila: A laboratory handbook. (Cold Spring Harbor Laboratory Press, 2005).

64. Wiegmann, B. M., Trautwein, M. D., Winkler, I. S., Barr, N. B., Kim, J.-W., Lambkin, C., Bertone, M. A., Cassel, B. K., Bayless, K. M., Heimberg, A. M., Wheeler, B. M., Peterson, K. J., Pape, T., Sinclair, B. J., Skevington, J. H., Blagoderov, V., Caravas, J., Kutty, S. N., Schmidt-Ott, U., Kampmeier, G. E., Thompson, F. C., Grimaldi, D. A., Beckenbach, A. T., Courtney, G. W., Friedrich, M., Meier, R. & Yeates, D. K. Episodic radiations in the fly tree of life. Proc. Natl. Acad. Sci. U. S. A. 108, 5690–5695 (2011).

65. Tomancak, P., Beaton, A., Weiszmann, R., Kwan, E., Shu, S., Lewis, S. E., Richards, S., Ashburner, M., Hartenstein, V., Celniker, S. E. & Rubin, G. M. Systematic determination of patterns of gene expression during *Drosophila* embryogenesis. Genome Biol. 3, research0088.1 (2002).

66. Tomancak, P., Berman, B. P., Beaton, A., Weiszmann, R., Kwan, E., Hartenstein, V., Celniker, S. E. & Rubin, G. M. Global analysis of patterns of gene expression during *Drosophila* embryogenesis. Genome Biol. 8, R145 (2007).

67. Lécuyer, E., Yoshida, H., Parthasarathy, N., Alm, C., Babak, T., Cerovina, T., Hughes, T. R., Tomancak, P. & Krause, H. M. Global analysis of mRNA localization reveals a prominent role in organizing cellular architecture and function. Cell 131, 174–187 (2007).

